# *In vivo*–Active Soluble Epoxide Hydrolase–targeting PROTACs with Improved Potency and Stability

**DOI:** 10.1101/2024.07.23.604814

**Authors:** Keita Nakane, Christophe Morisseau, Presley D. Dowker-Key, Gabrielle Benitez, Jennifer T. Aguilan, Emiko Nagai, Simone Sidoli, Bruce D. Hammock, Ahmed Bettaieb, Kosaku Shinoda, Seiya Kitamura

## Abstract

Soluble epoxide hydrolase (sEH) is a bifunctional enzyme involved in fatty acid metabolism and promising drug target. We previously reported first-generation sEH proteolysis–targeting chimeras (PROTACs) with limited degradation potency and low aqueous and metabolic stability. Herein, we report the development of next-generation sEH PROTAC molecules with improved stability and degradation potency. One of the most potent molecules (compound **8**) exhibits a half-maximal degradation concentration in the sub-nM range, is stable *in vivo*, and effectively degrades sEH in mouse livers and brown adipose tissues. Given the role played by sEH in many metabolic and nonmetabolic diseases, the presented molecules provide useful chemical probes for the study of sEH biology. They also hold potential for therapeutic development against a range of disease conditions, including diabetes, inflammation, and metabolic disorders.

---

Soluble epoxide hydrolase (sEH) is responsible for the enzymatic conversion of epoxy fatty acids (EpFAs) into their corresponding vicinal diols in the arachidonic acid cascade. These lipid mediators are important endogenous signaling molecules in glucose homeostasis, vascular regulation, and pain. Inhibition of sEH hydrolase activity by small-molecule sEH inhibitors (sEHis) is considered to be a promising approach for the treatment of diseases such as diabetes, neuropathic pain, chronic obstructive pulmonary disease (COPD), and metabolic disorders.^1–3^ Cellular mechanistic studies have demonstrated that the beneficial effects of sEHis, which stem, in part, from the reduction in endoplasmic reticulum (ER) stress caused by increased EpFA levels.^4–6^ A few sEHis, including GSK2256294A and EC5026, have been assessed in clinical trials (**Fig. 1A**), and several clinical trials have tested the efficacy of sEHis against a variety of disease indications, including impaired glucose tolerance,^7^ insulin resistance,^8,9^ COPD,^10^ and diabetic neuropathy.^11^ However, their therapeutic efficacy is limited in clinical trials, and unfortunately, no sEHis are yet in clinical use. This may be attributed to traditional, occupancy-driven inhibitors being unable to completely block enzymatic function *in vivo* because **(a)** high sEH protein levels, reaching up to the sub-µM range in the liver fraction,^12^ require high inhibitor concentrations to show efficacy, even with nM affinity; and **(b)** extremely high catalytic site occupancy is required to block the conversion of endogenous substrate due to high enzymatic efficiency (*K*_M_ = 2.0 µM, *k*_cat_ = 0.26 s^−1^ against 11,12-epoxyeicosatrienoic acid)^13^ and low concentrations of endogenous substrate epoxides (10–30 ng/g liver^14^).

**Figure 1.**
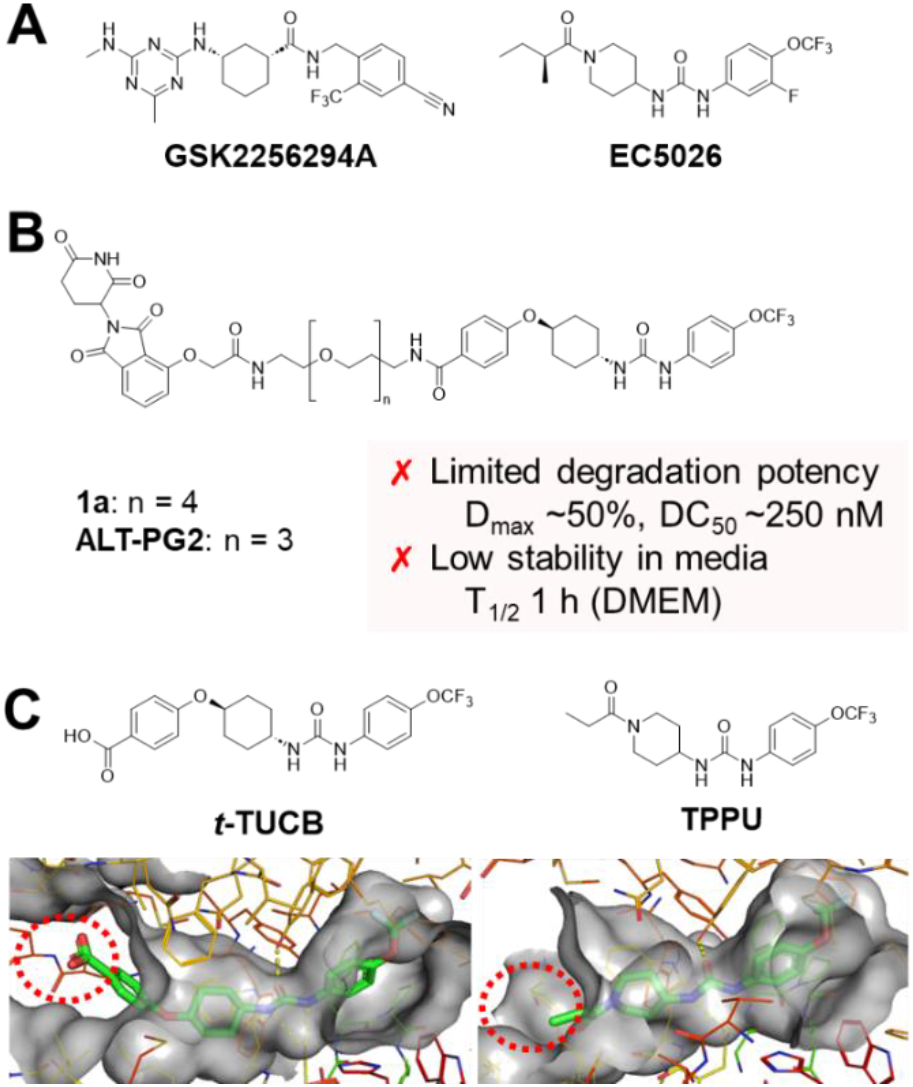
(A) Chemical structures of clinical candidate sEHis GSK2256294A and EC5026. (B) Chemical structures of first-generation sEH PROTAC compounds **1a** and **ALT-PG2**. (C) The sEHi scaffolds used in this study, *t*-TUCB and TPPU, and their X-ray crystal structures when complexed with human sEH (*Left panel, t*-TUCB, PDB: 6AUM. *Right panel*, TPPU, PDB: 4OD0). The solvent–exposed portions of the molecules are highlighted by dotted red circles.

To overcome the limitations of the conventional sEHi approach, we recently developed first-in-class small-molecule sEH degraders based on proteolysis-targeting chimera (PROTAC) technology.^15^ Optimized compound **1a** (**Fig. 1B**) induced sEH degradation in the low nM range in human hepatoma (HepG2) and kidney (HEK293T) cells. Moreover, compound **1a** exhibited superior efficacy in reducing cellular ER stress *in vitro* compared to the parent sEHi. More recently, an independent study validated that **ALT-PG2** (**Fig. 1B**), a structural analog of compound **1a**, efficiently reduced ER stress in primary cells and *in vivo*.^16^ However, these first-generation sEH PROTAC molecules demonstrate limited degradation potency with low aqueous stability, which substantially hinders their use in biological testing, including further *in vivo* studies and future clinical uses.

Herein, we report the development of sEH PROTAC molecules with improved degradation potency and stability. We performed a medicinal chemistry campaign to design and synthesize compound **1a** analogs with various E3 recruiters and linkers, followed by cellular sEH degradation measurements. This effort led to the development of metabolically stable next-generation PROTAC molecules that effectively degrade sEH *in vivo*.

While evaluating the cellular activity of first-generation sEH PROTAC molecules, compounds **1a** and **ALT-PG2** were found to be almost completely degraded in cell culture medium (DMEM) after incubation for 24 h (**Fig. S1**). We hypothesized that this was due to hydrolysis of the *O*-linked thalidomide moiety. Thus, we selected a range of cereblon-recruiting fragments and linkers that were previously reported to exhibit greater aqueous stability than the *O*-linked thalidomide moiety.^17–19^ In addition to the *t*-TUCB (4-[[*trans*-4-[[[[4-(trifluoromethoxy)phenyl]amino]carbonyl]amino]cyclohexyl]oxy]benzoic acid) scaffold, which was used as an sEH-binding moiety in the development of compounds **1a** and **ALT-PG2**, the TPPU (1-(1-propanoylpiperidin-4-yl)-3-[4-(trifluoromethoxy)phenyl]urea) scaffold was employed to determine the influence of the sEH-binding moiety on the molecule’s degradation potency and stability (**Fig. 1C**). The X-ray structures of these molecules when bound to human sEH indicated that portions of the molecules were exposed to the solvent, making them susceptible to modification (**Fig. 1C**). Subsequently, we designed eight new PROTAC candidate molecules based on *t-*TUCB and five additional molecules based on TPPU (**Tables 1** and **2**). Synthesis of these molecules is described in **Schemes 1** and **2**. Briefly, E3 ligase recruiter fragments, linkers, and sEH-binding moieties were connected via amide coupling. The TPPU scaffold was synthesized by reacting the corresponding isocyanate with Boc-protected 4-aminopiperidine, as previously described (**Scheme 2**).^20^ Details regarding the synthesis and characterization of the developed PROTAC molecules are described in the Supporting Information.

**Table 1.** Structures of *t-*TUCB-based sEH PROTAC Molecules, Cellular sEH Degradation Ability, and Biochemical Inhibitory Potency Against sEH.

| # |  | % sEH degradation<sup>a</sup> | human sEH IC<sub>50</sub> (nM)<sup>b</sup> | mouse sEH IC<sub>50</sub> (nM)<sup>b</sup> |
| --- | --- | --- | --- | --- |
| ***t*-TUCB**<br>(inhibitor) |  | - | 0.46 | 1.6 |
| **1a** |  | 35% | 0.50 | 4.0 |
| **1** |  | 30% | 1.3 | 1.6 |
| **2** |  | -8% | 1.2 | 0.64 |
| **3** |  | -2% | 1.3 | 4.7 |
| **4** |  | 10% | 0.24 | 6.0 |
| **5** |  | 32% | 1.1 | 9.3 |
| **6** |  | 79% | 1.7 | 9.9 |
| **7** |  | 56% | 5.0 | 20 |
| **8** |  | 78% | 3.8 | 210 |

<sup>a</sup> Cellular sEH degradation in HepG2 cells with 1 μM compound for 24 h, measured in triplicate by ELISA. SD values are listed in **Table S1**. The sEH concentration was normalized based on the total protein concentration, and % degradation was calculated relative to DMSO-treated samples ( $n = 3$ ).
<sup>b</sup> Biochemical inhibitory potency against recombinant purified human and mouse sEH using a fluorescent substrate for hydrolase activity.<sup>22,23</sup> Reported IC<sub>50</sub> values are the average of triplicate measurements with at least two data points above and at least two below the IC<sub>50</sub>. The fluorescent-based assay has a standard error of 10–20%, suggesting that differences of 2-fold or greater are significant.

**Table 2.**
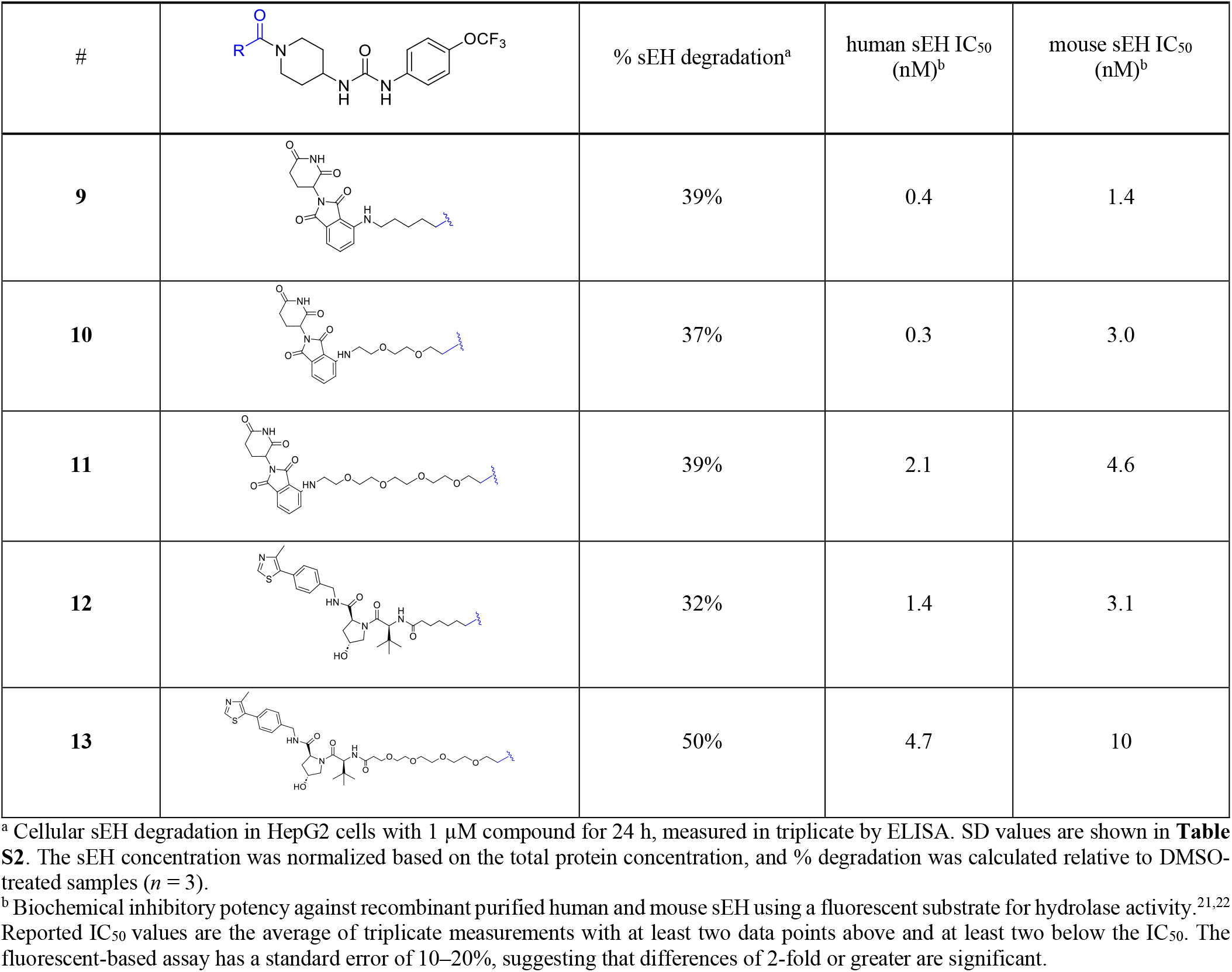
Structures of TPPU-based sEH PROTAC Molecules, Cellular s EH Degradation Ability, and Biochemical Inhibitory Potency Against sEH.

| # |  | % sEH degradation<sup>a</sup> | human sEH IC<sub>50</sub> (nM)<sup>b</sup> | mouse sEH IC<sub>50</sub> (nM)<sup>b</sup> |
| --- | --- | --- | --- | --- |
| 9 |  | 39% | 0.4 | 1.4 |
| 10 |  | 37% | 0.3 | 3.0 |
| 11 |  | 39% | 2.1 | 4.6 |
| 12 |  | 32% | 1.4 | 3.1 |
| 13 |  | 50% | 4.7 | 10 |

<sup>a</sup> Cellular sEH degradation in HepG2 cells with 1 μM compound for 24 h, measured in triplicate by ELISA. SD values are shown in **Table S2**. The sEH concentration was normalized based on the total protein concentration, and % degradation was calculated relative to DMSO-treated samples ( $n = 3$ ).
<sup>b</sup> Biochemical inhibitory potency against recombinant purified human and mouse sEH using a fluorescent substrate for hydrolase activity.<sup>21,22</sup> Reported IC<sub>50</sub> values are the average of triplicate measurements with at least two data points above and at least two below the IC<sub>50</sub>. The fluorescent-based assay has a standard error of 10–20%, suggesting that differences of 2-fold or greater are significant.

The sEH degradation potency of the synthesized molecules was first evaluated at 1 µM on HepG2 cells for 24 h. ELISA was utilized for sEH quantification given its high throughput and quantitative nature. As shown in **Tables 1** and **2**, subtle changes in linker and E3 recruiter structures appeared to influence the sEH degradation potency of the molecules. Importantly, the *t*-TUCB series of molecules with a piperazine linker (compounds **6**–**8**) demonstrated superior sEH degradation activity compared to the original PROTAC compound **1a**. These piperazine linkers and E3 recruiters have been employed in several of the first PROTAC molecules that are moving into clinical trials, such as ARV-110 and ARV471. Phenyl dihydrouracil and phenyl glutarimide are reportedly alternative cereblon binders with improved stability.^17,18^ Unfortunately, molecules with these fragments (compounds **3** and **4**) did not induce significant sEH degradation. In addition to assessing the cellular sEH degradation capability, the biochemical enzyme inhibitory potency was measured against human and mouse sEH using a fluorogenic substrate. All molecules exhibited sEHi activity ranging from sub-to-low nM potency (**Tables 1** and **2**).

**Scheme 1.**
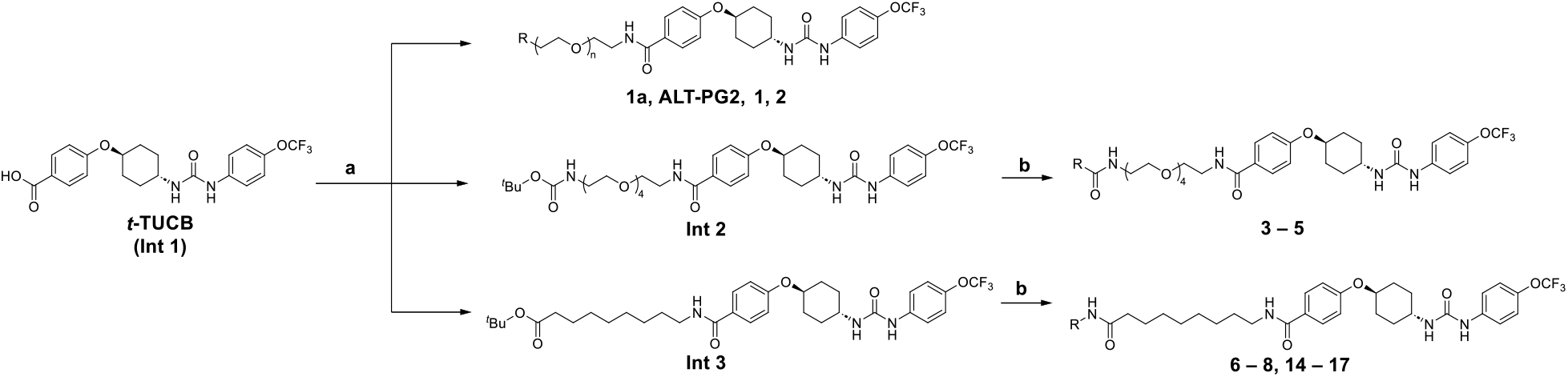
Synthesis of *t-*TUCB-based sEH PROTACs. a) HATU and DIEPA in DMF, RT; b) i) trifluoroacetic acid, RT. ii) HATU and DIEPA in DMF, RT.

**Scheme 2.**
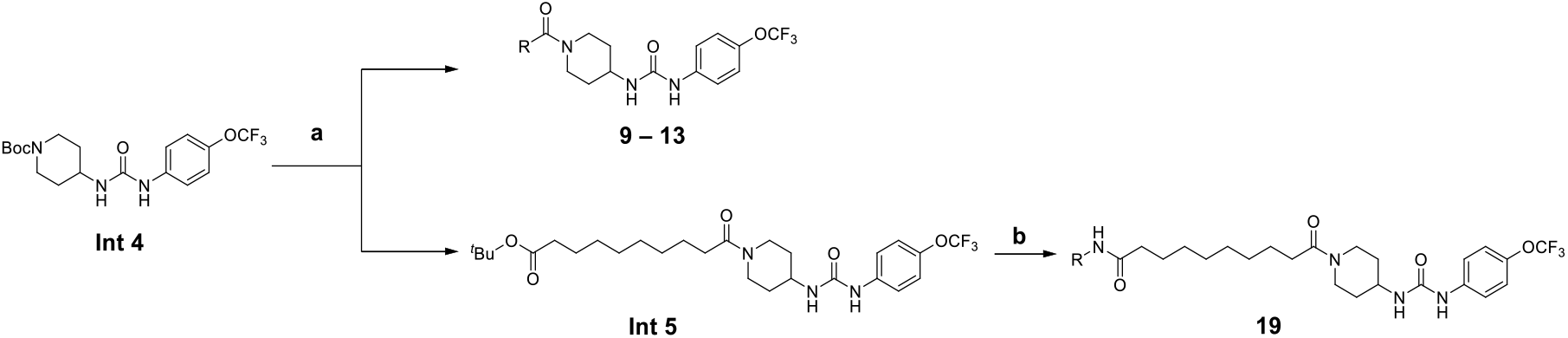
Synthesis of TPPU-based sEH PROTAC Molecules. a) i) Trifluoroacetic acid, RT; ii) HATU and DIEPA in DMF, RT.b) i) Trifluoroacetic acid, RT; ii) HATU and DIEPA in DMF, RT.

Encouraged by the initial structure–degradation relationship results (**Tables 1** and **2**), compounds **6** and **8** were chosen for further characterization. First, their aqueous stability in cell culture medium (DMEM) was measured, revealing higher stability than compound **1a** in the medium (**Fig. 2A**). In addition, the *in vitro* metabolic stability of compounds **6** and **8** was assessed with mouse liver microsomes (**Fig. 2B**), demonstrating improved stability compared to **1a**.

**Figure 2.**
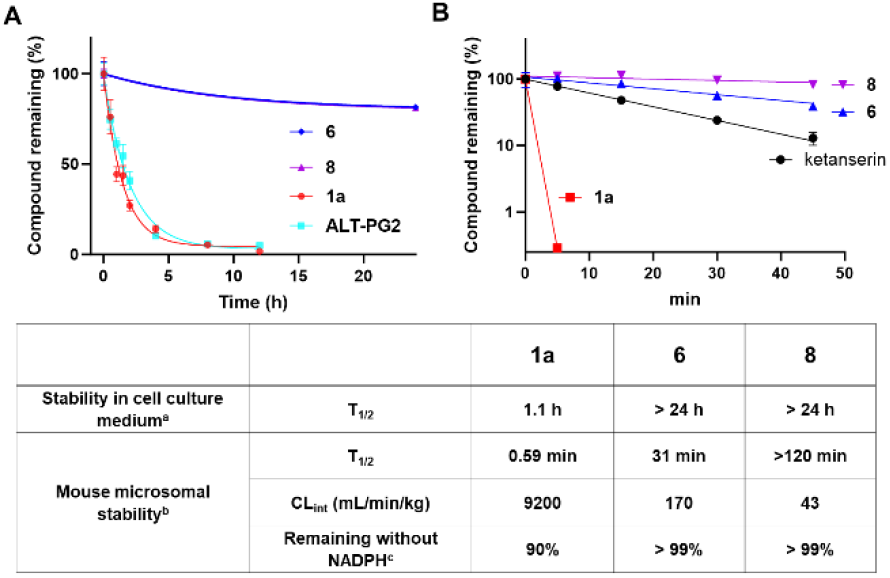
Compounds **6** and **8** have improved stability compared to first-generation sEH PROTAC compounds **1a** and **ALT-PG2**. (A) Chemical stability in cell culture medium. (B) *In vitro* metabolic stability. Ketanserin was used as an assay control. ^a^ Stability in cell culture medium was measured in DMEM. ^b^ Microsomal stability was measured by incubating 500 nM compound with mouse liver microsomes (0.25 mg/mL) and NADPH (1 mM) solution for 0–45 minutes, in duplicate (**Table S3**). ^c^ Values measured without NADPH after incubation for 45 min.

To evaluate their precise degradation potencies, we next determined the dose-dependent sEH degradation induced by compounds **6** and **8**. Compound **8** was found to have improved degradation potency compared to compound **6**, with a half-maximal degradation concentration (DC_50_) of approximately 0.5 nM (**Fig. 3A**). The dose–response experiments with compound **8** over a wide concentration range (0.01–1000 nM) revealed maximum sEH degradation at approximately 100 nM, with a maximal sEH degradation level (D_max_) of 79% (**Fig. S2**).

**Figure 3.**
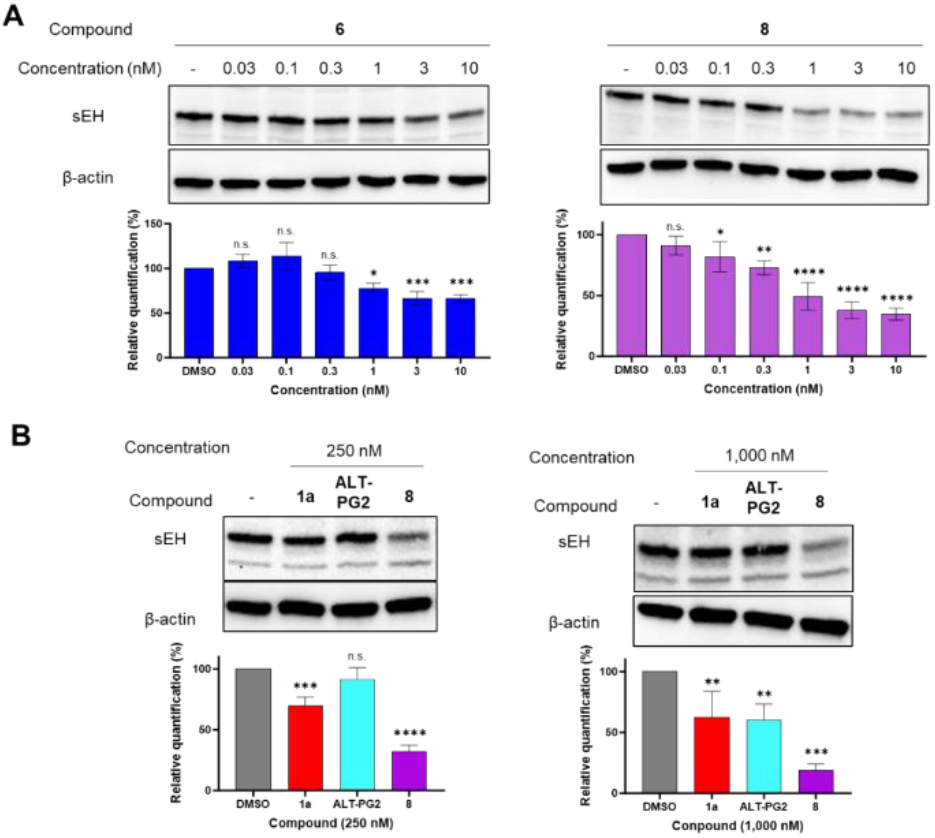
Compounds **6** and **8** have greater sEH degradation potency than first-generation sEH PROTAC compounds **1a** and **ALT-PG2**. (A) Concentration-dependent degradation of sEH induced by compounds **6** and **8** (0.03–10 nM). Differences between treatment groups were analyzed by one-way ANOVA with the Holm–Sidak test (^****^*p* < 0.0001, ^***^*p* < 0.001, ^**^*p* < 0.01, ^*^*p* < 0.04, n.s.: not significant). (B) Direct comparison of sEH degradation induced by compounds **1a** and **ALT-PG2** *vs*. compound **8**. Values represent the mean ± SD (*n* = 3). Differences between treatment groups were analyzed by one-way ANOVA with the Holm– Sidak test (^****^*p* < 0.0001, ^***^ *p* < 0.001, ^**^*p* < 0.01).

The improvement in degradation potency was further assessed by direct comparison of compound **8** with first-generation PROTAC compounds **1a** and **ALT-PG2** at two different concentrations. The results aligned with the ELISA data, demonstrating that compound **8** induced significantly higher sEH degradation than compounds **1a** and **ALT-PG2** (**Fig. 3B**). Compounds **1a** and **ALT-PG2** failed to induce 50% sEH degradation at 250 nM and 1 μM (100 nM–1 µM; DC_50_ > 1 µM, **Fig. S3**), whereas compound **8** robustly degraded >50% sEH at both concentrations. These data indicated a 500–2,000-fold improvement in DC_50_ for compound **8**, with a higher D_max_ value than first-generation PROTAC molecules. Taken together, the results demonstrated that compound **8** possessed significantly improved sEH degradation potency with improved metabolic and aqueous stability.

Based on the above findings, compound **8** was selected for further biological evaluation. First, the sEH degradation kinetics induced by compound **8** were determined (**Fig. 4A**). HepG2 cells were treated with 50 nM compound **8**, and the sEH quantity was measured at various incubation times via immunoblotting. Compound **8** induced >50% sEH degradation in as little as 4 h, with the highest degradation level achieved after incubation for 24 h. The degradation kinetics of compound **8** were faster than those of first-generation PROTAC compound **1a**, which required 16 h to achieve >50% sEH degradation.^15^

**Figure 4.**
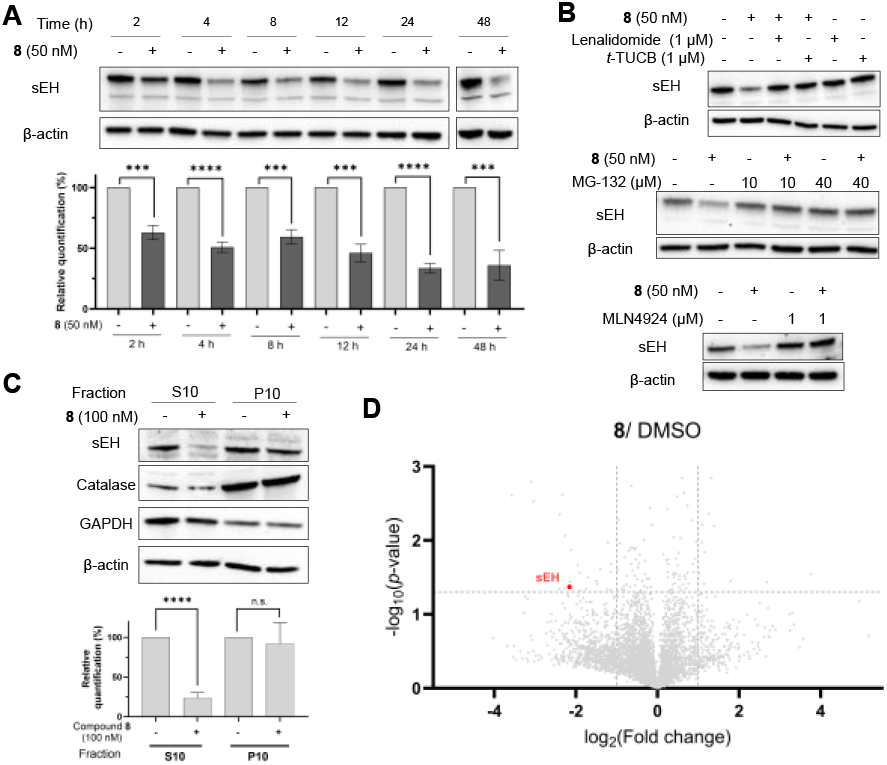
Detailed characterization of sEH degradation induced by compound **8**. (A) Degradation kinetics of sEH induced by 50 nM compound **8** treatment for 24 h. Whole-cell lysate was employed in immunoblotting. Values represent the mean ± SD (*n* = 3). Student’s *t*-test was performed, and *p*-values represent significant differences between compound treatment *vs*. vehicle control (^****^*p* < 0.0001, ^***^*p* < 0.001). (B) Validation of sEH degradation mechanism. Treatment with compound **8** alone or in combination with: *Top panel*; Lenalidomide (E3 ligase ligand) or *t-*TUCB (sEHi). *Middle panel*: Proteasome inhibitor MG-132. *Bottom panel*: NEDD8-activating enzyme inhibitor MLN4942. Whole-cell lysate was employed in immunoblotting. (C) Compound **8** selectively degraded cytosolic sEH but not peroxisomal sEH. Cytosol-selective sEH degradation was induced by 100 nM compound **8** treatment for 48 h. Cell organelles were fractionated into the cytosol-containing S10 fraction and peroxisome-containing P10 fraction, which were confirmed by immunoblotting with catalase (peroxisome marker) and GAPDH (cytoplasmic marker), respectively. Values represent the mean ± SD (*n* = 3). Statistical differences were analyzed by Student’s *t*-test. Differences were significant for S10 samples (^****^*p* < 0.0001) but not for P10 samples (n.s.). (D) Quantitative MS-based proteomic analysis indicated that sEH was one of the most significantly reduced proteins after compound **8** treatment. Data points representing sEH are shown in red. The vertical dashed lines and horizontal dashed line in the volcano plot represent log_2_ (fold change) cut-off of ±1.0 (fold change = ±2.0) and −log_10_ (*p*-value) cut-off of 1.30 (*p* = 0.05, *n* = 3).

Next, the mechanism of degradation was validated by cotreatment of compound **8** (50 nM) with *t-*TUCB (1 µM, sEHi), lenalidomide (1 µM, E3 ligase ligand), MLN4924 (1 µM, NEDD8-activating enzyme inhibitor), or MG-132 (10 or 40 µM, proteasome inhibitor). These cotreatments inhibited sEH degradation induced by compound **8**, supporting the PROTAC mechanism of degradation (**Fig. 4B**). We also explored subcellular selective sEH degradation in the cytosol and peroxisome. Aligned with the previous results of compound **1a**, compound **8** induced cytosol-selective sEH degradation over peroxisomal sEH degradation (**Fig. 4C**). Finally, quantitative MS-based proteomic analysis revealed that sEH levels were among those most significantly reduced following treatment with compound **8** (**Fig. 4D**). In addition to sEH, the expression of several proteins was significantly downregulated after compound **8** treatment, indicating potential off-target effects. The downregulated proteins did not have any known protein-protein interactions, expression regulation by sEH, or interaction with sEHis. Elucidating the mechanism responsible for the reduced levels of these proteins and improving sEH selectivity are the subjects of further study.

## Compound 8 effectively reduces cellular ER stress *in vitro*

Previous studies demonstrated that sEH is a physiological modulator of ER stress signaling,^4^ and sEH degradation reduces ER stress.^15,16^ Thus, the efficacy of PROTAC compounds in reducing ER stress was tested with HepG2 cells using the thapsigargin (Tg)-induced ER stress assay, focusing on the IRE1α and PERK signaling pathways. HEK293T cells were pretreated with compound **8** for 12 h, the medium was removed, and cells were treated with Tg to induce ER stress for an additional 24 h. Compound **8** treatment significantly reduced ER stress, as determined by decreased phosphorylation of IRE1α, PERK, and eIF2α at all tested concentrations (100 nM, 250 nM, and 1 µM) (**Figs. 5** and **S9**). By contrast, the parent sEHi, *t*-TUCB, and first-generation sEH PROTAC compound **1a** were less effective in reducing ER stress at a lower concentration (100 nM) than compound **8**. Consistent with these findings, compound **8** treatment resulted in improved cell survival and reduced Tg-induced apoptotic cell death compared to that with sEHis or compound **1a** (**Fig. S10**). These results demonstrated the improved cellular efficacy of compound **8** compared to sEHi *t*-TUCB and first-generation PROTAC compound **1a**.

**Figure 5.**
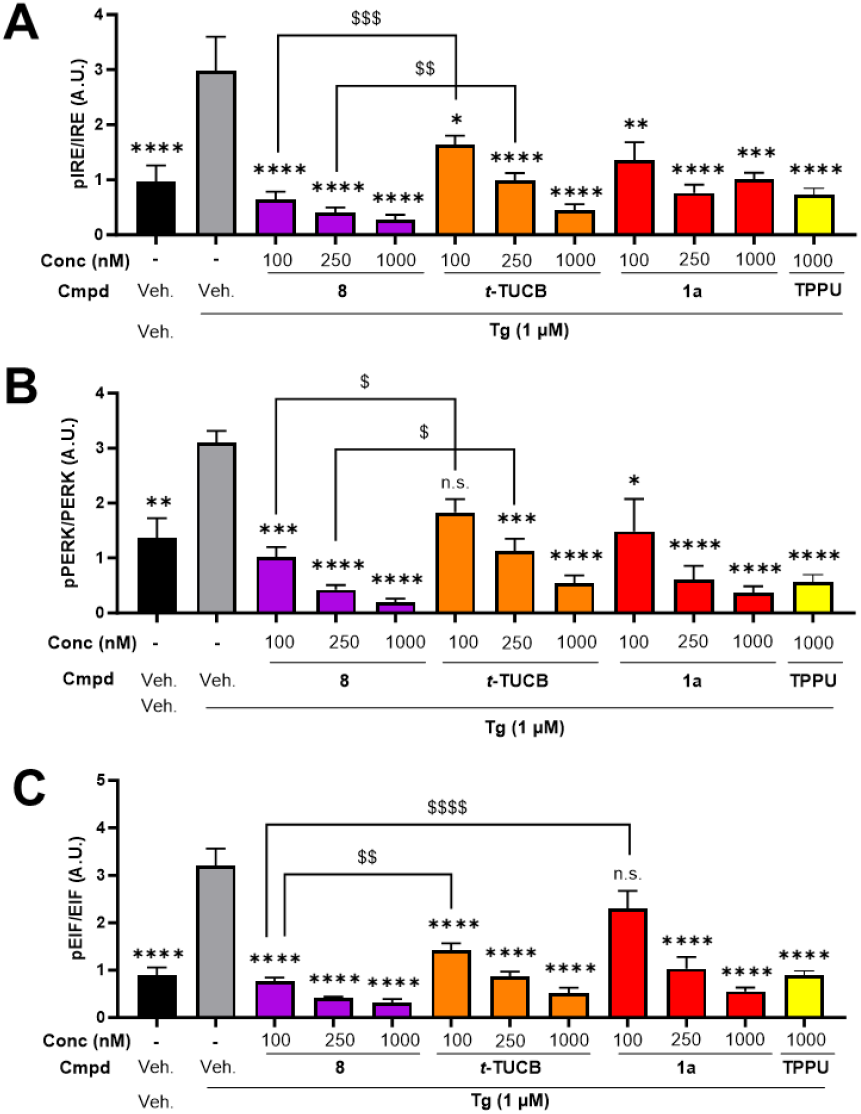
Compound **8** effectively reduces ER stress. HEK293T cells were treated with the indicated sEH modulators for 12 h prior to adding 1 µM Tg for an additional 24 h. Changes in ER stress were determined via immunoblotting with selected ER stress markers. (A) pIRE(S724)/IRE, (B) pPERK(T981)/PERK, and (C) pEIF2α(S51)/EIF2α. Values represent the mean ± SEM (*n* = 4). Differences between treatment groups were analyzed by one-way ANOVA with the Holm–Sidak test (\*\*\*\**p* < 0.0001, ^***^*p* < 0.001, ^**^*p* < 0.01, ^*^*p* < 0.04) or Student’s *t*-test (^$$$$^*p* < 0.0001, ^$$$^*p* < 0.004, ^$$^*p* < 0.009, ^$^*p* < 0.04). The original western blots are included in the Supporting Information.

Prior to preclinical *in vivo* testing of our compounds, an additional cycle of medicinal chemistry was conducted to broaden the range of candidate molecules and enhance our understanding of the structure–degradation relationship. Six additional sEH PROTAC candidate molecules were designed based on the structures of compounds **6** and **8** (**Table 3**). The sEH degradation activity of the synthesized molecules was evaluated by immunoblotting and ELISA in parallel. Compounds **17, 18**, and **19** exhibited comparable sEH degradation potency to compound **8**. Notably, compound **18**, an analog of compound **6**, with the lenalidomide 5′-piperazine fragment as an E3 ligase recruiter, was found to be relatively unstable in the medium, with 57% sEH remaining after incubation for 24 h (**Fig. S4**). Moreover, compound **18** was cytotoxic against HepG2 cells at 1 µM. By contrast, compound **17**, a structural analog of compound **8**, with the lenalidomide 5′-piperazine-4-methylpiperidine fragment as an E3 ligase recruiter, demonstrated high sEH degradation potency with better stability in the cell culture medium and no cytotoxic effects. Compound **19**, which possessed the TPPU scaffold as an sEH binder, with the E3 ligase recruiter and linker used in compound **8**, was found to have high degradation potency although it was less chemically stable than compound **8** (**Fig. S4**). Molecules with different linker positions (compounds **14** and **15**) induced less sEH degradation than the corresponding molecules with the linker at position 5 of pomalidomide, indicating the importance of the linker position in sEH degradation potency. Given these findings, we focused on compounds **17** and **19** with high sEH degradation activity and reasonable aqueous stability.

**Table 3.**
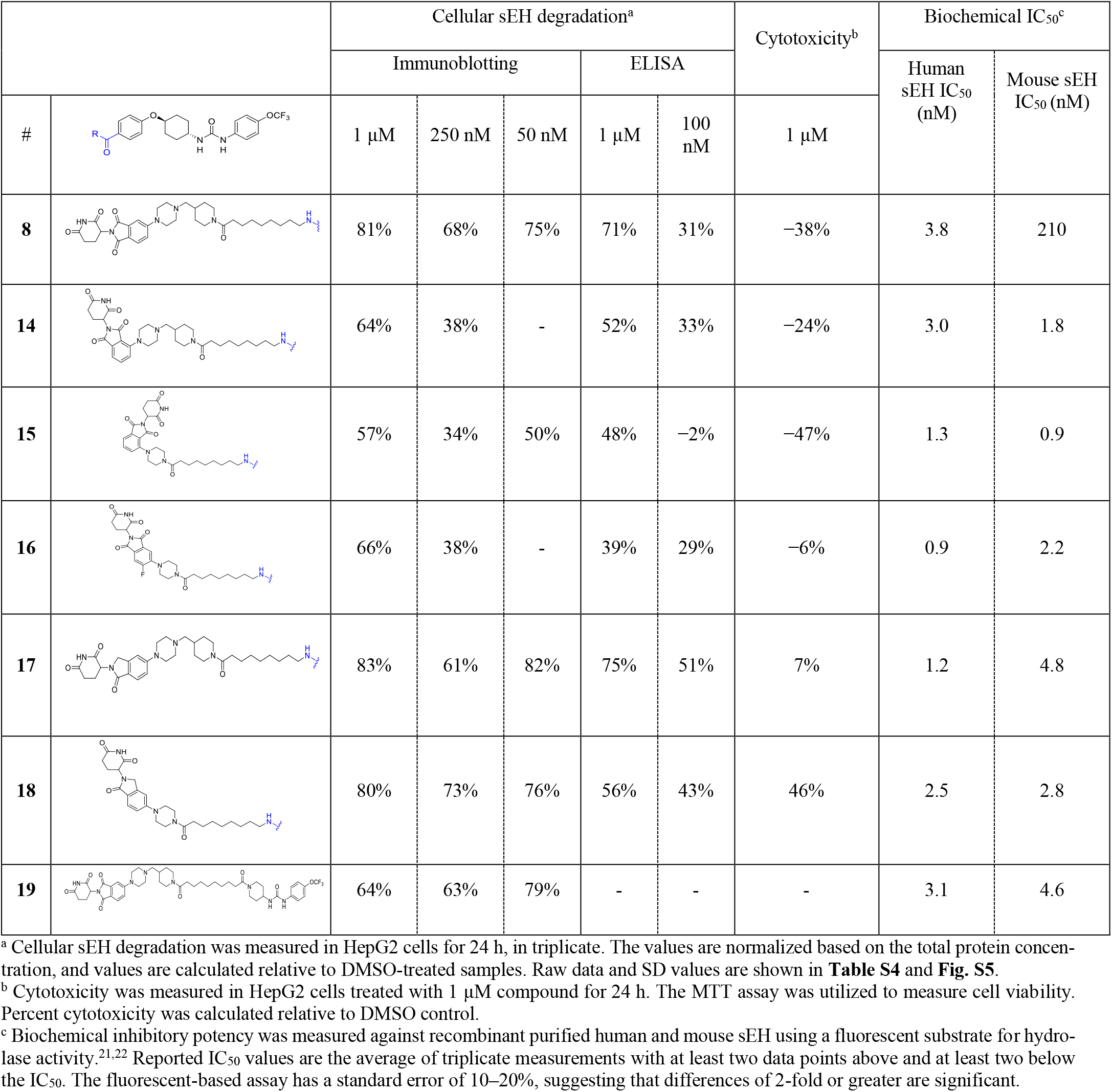
Structure–degradation Relationships of Compound 8 and its Analogs.

|  |  | Cellular sEH degradation<sup>a</sup> |  |  |  |  | Cytotoxicity<sup>b</sup> | Biochemical IC<sub>50</sub><sup>c</sup> |  |
| --- | --- | --- | --- | --- | --- | --- | --- | --- | --- |
|  |  | Immunoblotting |  |  | ELISA |  |  | Human sEH IC<sub>50</sub> (nM) | Mouse sEH IC<sub>50</sub> (nM) |
| # |  | 1 μM | 250 nM | 50 nM | 1 μM | 100 nM | 1 μM |  |  |
| 8 |  | 81% | 68% | 75% | 71% | 31% | −38% | 3.8 | 210 |
| 14 |  | 64% | 38% | - | 52% | 33% | −24% | 3.0 | 1.8 |
| 15 |  | 57% | 34% | 50% | 48% | −2% | −47% | 1.3 | 0.9 |
| 16 |  | 66% | 38% | - | 39% | 29% | −6% | 0.9 | 2.2 |
| 17 |  | 83% | 61% | 82% | 75% | 51% | 7% | 1.2 | 4.8 |
| 18 |  | 80% | 73% | 76% | 56% | 43% | 46% | 2.5 | 2.8 |
| 19 |  | 64% | 63% | 79% | - | - | - | 3.1 | 4.6 |

<sup>a</sup> Cellular sEH degradation was measured in HepG2 cells for 24 h, in triplicate. The values are normalized based on the total protein concentration, and values are calculated relative to DMSO-treated samples. Raw data and SD values are shown in **Table S4** and **Fig. S5**.
<sup>b</sup> Cytotoxicity was measured in HepG2 cells treated with 1 μM compound for 24 h. The MTT assay was utilized to measure cell viability. Percent cytotoxicity was calculated relative to DMSO control.
<sup>c</sup> Biochemical inhibitory potency was measured against recombinant purified human and mouse sEH using a fluorescent substrate for hydrolase activity.<sup>21,22</sup> Reported IC<sub>50</sub> values are the average of triplicate measurements with at least two data points above and at least two below the IC<sub>50</sub>. The fluorescent-based assay has a standard error of 10–20%, suggesting that differences of 2-fold or greater are significant.

The dose–response experiments with sEH PROTAC molecules (0.03–10 nM) in HepG2 cells showed that compounds **8** and **17** had comparable sEH degradation potencies, while compound **19** exhibited slightly less potency (**Fig. S6**). The lower sEH degradation activity of compound **19** compared to that of compound **8** may have been due to its lower stability in cell culture medium (**Fig. S4**). In addition to sEH degradation in HepG2 cells, these molecules effectively degraded sEH in HEK293T cells (**Fig. S7**). Further, compound **8** degraded sEH in human breast adenocarcinoma MDA-MB-231 cells and human bone marrow neuroblast SH-SY5Y cells (**Fig. S8**). Overall, among the synthesized compounds, compounds **8** and **17** were found to have the highest sEH degradation activity, followed by compound **19**. Importantly, these molecules shared the same linker but possessed different E3 ligase–recruiting moieties or different sEH-binding moieties, indicating the importance of the linker in sEH degradation.

## Pharmacokinetics and sEH degradation *in vivo*

Based on their degradation potencies and chemical stability, compounds **8, 17**, and **19** were selected for *in vivo* pharmacokinetic (PK) profiling. The compounds were administered to male CD1 mice (6–8 weeks, weighing 30–31 g, *n* = 3) via single i.p. injection (10 mg/kg). The plasma concentration of the compounds was monitored at various time points (**Fig. 6A**). Compound **8** was found to be most stable, with a half-life at 12 h, followed by compound **17**, with a half-life of approximately 7 h. TPPU series compound **19** was found to be least stable *in vivo*, with limited AUC values. The plasma concentration of compound **8** was maintained >1 µM (>1,000-fold higher than DC_50_) over the course of PK profiling, even 24 h after the single i.p. injection. Based on the PK data, along with the cellular degradation potency, compound **8** was selected for evaluation *in vivo*. Male C57BL/6J mice (5 weeks, weighing 22–28 g, *n* = 3) were treated with compound **8** via single i.p. injection (12 mg/kg). After 24 h, mice were sacrificed and sEH levels in the liver and brown adipose tissue (BAT) were measured via immunoblotting. Significant sEH degradation in the liver and BAT (**Fig. 6B** and **6C**) demonstrated effective *in vivo* degradation of sEH by the developed PROTAC compounds.

**Figure 6.**
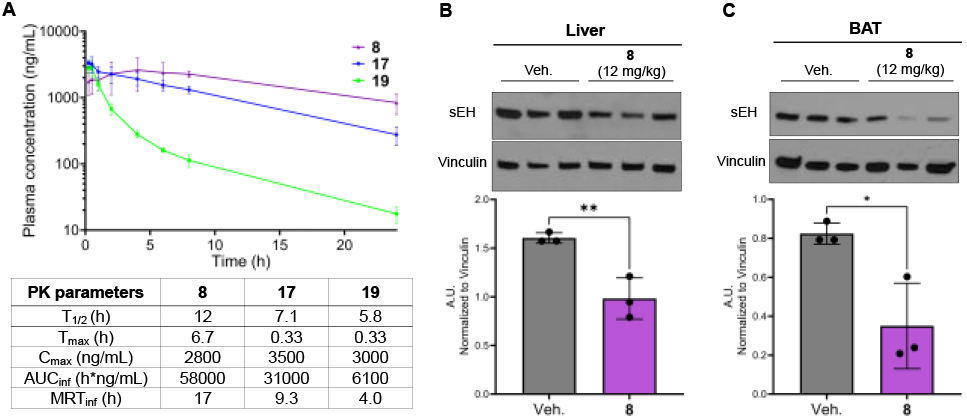
*In vivo* evaluation of compound **8**. (A) Pharmacokinetic profile (*Top panel*) and pharmacokinetic parameters (*bottom panel*) of compounds **8, 17**, and **19** (Male CD1 mice, *n* = 3 per group, 10 mg/kg i.p.). Values represent the mean ± SD (*n* = 3) (**Table S5**). (B, C) Compound **8** induces sEH degradation *in vivo*. Male C57BL/6J mice (*n* = 3 per group) received a single i.p. injection of vehicle (Veh) or compound **8** at a dosage of 12 mg/kg. After 24 h, mice were sacrificed and sEH levels in the liver (B) and BAT (C) were measured via immunoblotting. Values represent the mean ± SEM (*n* = 3). Student’s *t*-test was performed, and *p*-values represent significant differences between compound treatment *vs*. vehicle control (^**^*p* < 0.01, ^*^*p* < 0.04).

In conclusion, we conducted a medicinal chemistry campaign to develop sEH PROTAC molecules with improved degradation potency and stability, which led to the development of *in vivo–active* sEH degraders. The newly developed sEH PROTAC molecules containing piperazine linkers demonstrated high chemical and metabolic stability and potent sEH degradation activity with DC_50_ values as low as sub-nM levels. The mechanism of degradation was validated to occur via the ubiquitin–proteasome system. Administration of compound **8** to mice induced significant sEH degradation in the liver and BAT. These compounds will provide useful chemical probes for the study of sEH function, with direct therapeutic potential against various types of diseases. Furthermore, this study demonstrates the successful development of *in vivo–*active PROTAC molecules through linker optimization, yielding improved sEH degradation potency and chemical and metabolic stability.

## Supporting information

Original data of immunoblotting

## ASSOCIATED CONTENT

### Supporting Information

The Supporting Information is available free of charge on the ACS Publications website. The details of inhibitory potency of PROTAC compounds; statistical details; additional cellular data of the degradation mechanisms of sEH PROTAC compounds; synthetic methods and compound characterization; original data from immunoblotting analysis; and proteomics data are provided in the Supporting Information.

## AUTHOR INFORMATION

### Author contributions

Conceptualization: K.N. and S.K.; Synthesis and chemical characterization: K.N.; Data acquisition and analysis: K.N., C.M., P.D.K., J.T.A., G.B., S.S., A.B., K.S., and S.K.; Experimental setup: K.N., E.N., and S.K.; Project administration: B.D.H. and S.K.; Writing— original draft preparation: S.K.; Writing—figure preparation, text editing and revisions: K.N. and S.K. All authors have read, reviewed, edited, and agreed to the published version of the manuscript.

### Notes

C.M. and B.D.H. are inventors of patents related to the use of sEH inhibitors owned by the University of California. S.K., K.N. C.M., and B.D.H. hold a patent application for the compounds described in this paper (Provisional Application No. 63/642,039).

## ACKNOWLEDGMENT

We thank Dr. S. H. Hwang (University of California, Davis) for kindly providing sEH inhibitor *t*-TUCB. The authors also thank Dr. R. J. Eddy (Department of Pathology, Albert Einstein College of Medicine) for kindly providing human breast adenocarcinoma cells (MDA-MB-231) and Dr. Q. Zhang (Department of Molecular Pharmacology, Albert Einstein College of Medicine) for providing human bone marrow neuroblast cells (SH-SY5Y). Finally, we would like to thank Dr. D. W. Wolan for reviewing the manuscript and providing constructive comments.

## ABBREVIATIONS

ANOVA: analysis of variance
AUC: area under the curve
BAT: brown adipose tissue
CL_int_: intrinsic clearance
C_max_: maximum concentration
COPD: chronic obstructive pulmonary disease
DC_50_: half-maximal degradation concentration
DIEPA: *N,N*-diisopropylamine
D_max_: maximal level of degradation
DMEM: Dulbecco’s Modified Eagle Medium
DMF: *N,N*-dimethylformamide
DMSO: dimethyl sulfoxide
eIF2α: eukaryotic translation initiation factor 2 alpha
EpFA: epoxy fatty acid
ER: endoplasmic reticulum
HATU: 1-[bis(dimethylamino)methylene]-1H-1,2,3-triazolo[4,5-b]pyridinium 3-oxide hexafluorophosphate
i.p.: intraperitoneal
IC_50_: half-maximal inhibitory concentration
inf: to infinity
int: intermediate
IRE1α: inositol-requiring enzyme 1 alpha
*k*_cat_: turnover number
*K*_M_: Michaelis–Menten constant
MRT: mean residence time
n.s.: not significant
NADPH: nicotinamide adenine dinucleotide phosphate
NEDD8: neural precursor cell expressed developmentally downregulated protein 8
PDB: protein databank
PERK: protein kinase RNA-like ER kinase
PK: pharmacokinetics
PROTAC: proteolysis-targeting chimera
RT: room temperature
SD: standard deviation
sEH: soluble epoxide hydrolase
sEHi: sEH inhibitor
SEM: standard error of the mean
T_1/2_: half-life
Tg: thapsigargin
T_max_: time of maximum concentration
Veh: vehicle.

## Table of Contents

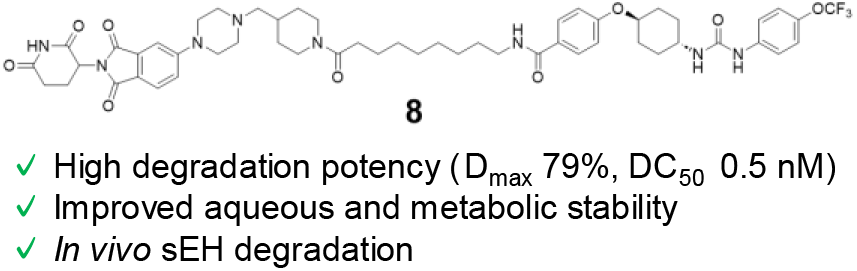

## REFERENCES

1. Wagner, K. M.; McReynolds, C. B.; Schmidt, W. K.; Hammock, B. D. Soluble Epoxide Hydrolase as a Therapeutic Target for Pain, Inflammatory and Neurodegenerative Diseases. Pharmacol. Ther. 2017, 180, 62–76. DOI: 10.1016/j.pharmthera.2017.06.006

2. Wang, W.; Yang, J.; Zhang, J.; Wang, Y.; Hwang, S. H.; Qi, W.; Wan, D.; Kim, D.; Sun, J.; Sanidad, K. Z.; Yang, H.; Park, Y.; Liu, J. Y.; Zhao, X.; Zheng, X.; Liu, Z.; Hammock, B. D.; Zhang, G. Lipidomic Profiling Reveals Soluble Epoxide Hydrolase as a Therapeutic Target of Obesity-Induced Colonic Inflammation. Proc. Natl. Acad. Sci. U. S. A. 2018, 115 (20), 5283–5288. DOI: 10.1073/pnas.1721711115

3. Fishbein, A.; Wang, W.; Yang, H.; Yang, J.; Hallisey, V. M.; Deng, J.; Verheul, S. M. L.; Hwang, S. H.; Gartung, A.; Wang, Y.; Bielenberg, D. R.; Huang, S.; Kieran, M. W.; Hammock, B. D.; Panigrahy, D. Resolution of Eicosanoid/Cytokine Storm Prevents Carcinogen and Inflammation-Initiated Hepatocellular Cancer Progression. Proc. Natl. Acad. Sci. U. S. A. 2020, 117 (35), 21576– 21587. DOI: 10.1073/pnas.2007412117

4. Bettaieb, A.; Nagata, N.; AbouBechara, D.; Chahed, S.; Morisseau, C.; Hammock, B. D.; Haj, F. G. Soluble Epoxide Hydrolase Deficiency or Inhibition Attenuates Diet-Induced Endoplasmic Reticulum Stress in Liver and Adipose Tissue. J. Biol. Chem. 2013, 288 (20), 14189–14199. DOI: 10.1074/jbc.M113.458414

5. Inceoglu, B.; Bettaieb, A.; Haj, F. G.; Gomes, A. V.; Hammock, B. D. Modulation of Mitochondrial Dysfunction and Endoplasmic Reticulum Stress Are Key Mechanisms for the Wide-Ranging Actions of Epoxy Fatty Acids and Soluble Epoxide Hydrolase Inhibitors. Prostaglandins Other Lipid Mediat. 2017, 133, 68–78. DOI: 10.1016/j.prostaglandins.2017.08.003

6. Jiang, X. S.; Xiang, X. Y.; Chen, X. M.; He, J. L.; Liu, T.; Gan, H.; Du, X. G. Inhibition of Soluble Epoxide Hydrolase Attenuates Renal Tubular Mitochondrial Dysfunction and ER Stress by Restoring Autophagic Flux in Diabetic Nephropathy. Cell Death Dis. 2020, 11 (5), 385. DOI: 10.1038/s41419-020-2594-x

7. Chen, D.; Whitcomb, R.; MacIntyre, E.; Tran, V.; Do, Z. N.; Sabry, J.; Patel, D. V.; Anandan, S. K.; Gless, R.; Webb, H. K. Pharmacokinetics and Pharmacodynamics of AR9281, an Inhibitor of Soluble Epoxide Hydrolase, in Single- and Multiple-Dose Studies in Healthy Human Subjects. J. Clin. Pharmacol. 2012, 52 (3), 319–328. DOI: 10.1177/0091270010397049

8. Luther, J. M.; Ray, J.; Wei, D.; Koethe, J. R.; Hannah, L.; DeMatteo, A.; Manning, R.; Terker, A. S.; Peng, D.; Nian, H.; Yu, C.; Mashayekhi, M.; Gamboa, J.; Brown, N. J. GSK2256294 Decreases SEH (Soluble Epoxide Hydrolase) Activity in Plasma, Muscle, and Adipose and Reduces F2-Isoprostanes but Does Not Alter Insulin Sensitivity in Humans. Hypertension 2021, 78 (4), 1092–1102. DOI: 10.1161/HYPERTENSIONAHA.121.17659

9. Lazaar, A. L.; Yang, L.; Boardley, R. L.; Goyal, N. S.; Robertson, J.; Baldwin, S. J.; Newby, D. E.; Wilkinson, I. B.; Tal-Singer, R.; Mayer, R. J.; Cheriyan, J. Pharmacokinetics, Pharmacodynamics and Adverse Event Profile of GSK2256294, a Novel Soluble Epoxide Hydrolase Inhibitor. Br. J. Clin. Pharmacol. 2016, 81 (5), 971–979. DOI: 10.1111/bcp.12855

10. Yang, L.; Cheriyan, J.; Gutterman, D. D.; Mayer, R. J.; Ament, Z.; Griffin, J. L.; Lazaar, A. L.; Newby, D. E.; Tal-Singer, R.; Wilkinson, I. B. Mechanisms of Vascular Dysfunction in COPD and Effects of a Novel Soluble Epoxide Hydrolase Inhibitor in Smokers. Chest 2017, 151 (3), 555–563. DOI: 10.1016/j.chest.2016.10.058

11. Hammock, B. D.; McReynolds, C. B.; Wagner, K.; Buckpitt, A.; Cortes-Puch, I.; Croston, G.; Lee, K. S. S.; Yang, J.; Schmidt, W. K.; Hwang, S. H. Movement to the Clinic of Soluble Epoxide Hydrolase Inhibitor EC5026 as an Analgesic for Neuropathic Pain and for Use as a Nonaddictive Opioid Alternative. J. Med. Chem. 2021, 64 (4), 1856–1872. DOI: 10.1021/acs.jmed-chem.0c01886

12. Li, D.; Cui, Y.; Morisseau, C.; Gee, S. J.; Bever, C. S.; Liu, X.; Wu, J.; Hammock, B. D.; Ying, Y. Nanobody Based Immunoassay for Human Soluble Epoxide Hydrolase Detection Using Polymeric Horseradish Peroxidase (PolyHRP) for Signal Enhancement: The Rediscovery of PolyHRP? Anal. Chem. 2017, 89 (11), 6248–6256. DOI: 10.1021/acs.analchem.7b01247

13. Morisseau, C.; Inceoglu, B.; Schmelzer, K.; Tsai, H. J.; Jinks, S. L.; Hegedus, C. M.; Hammock, B. D. Naturally Occurring Monoepoxides of Eicosapentaenoic Acid and Docosahexaenoic Acid Are Bioactive Antihyperalgesic Lipids. J. Lipid Res. 2010, 51 (12), 3481–3490. DOI: 10.1194/jlr.M006007

14. Diani-Moore, S.; Ma, Y.; Gross, S. S.; Rifkind, A. B. Increases in Levels of Epoxyeicosatrienoic and Dihydroxyeicosatrienoic Acids (EETs and DHETs) in Liver and Heart In Vivo by 2,3,7,8-tetrachlorodibenzo-p-dioxin (TCDD) and in Hepatic EET:DHET Ratios by Cotreatment with TCDD and the Soluble Epoxide Hydrolase Inhibitor AUDA. Drug Metab. Dispos. 2014, 42 (2), 294–300. DOI: 10.1124/dmd.113.055368

15. Wang, Y.; Morisseau, C.; Takamura, A.; Wan, D.; Li, D.; Sidoli, S.; Yang, J.; Wolan, D. W.; Hammock, B. D.; Kitamura, S. PROTAC-Mediated Selective Degradation of Cytosolic Soluble Epoxide Hydrolase Enhances ER Stress Reduction. ACS Chem. Biol. 2023, 18 (4), 884–896. DOI: 10.1021/acschembio.3c00017

16. Peyman, M.; Barroso, E.; Turcu, A. L.; Estrany, F.; Smith, D.; Jurado-Aguilar, J.; Rada, P.; Morisseau, C.; Hammock, B. D.; Valverde, Á.M.; Palomer, X.; Galdeano, C.; Vázquez, S.; Vázquez-Carrera, M. Soluble Epoxide Hydrolase-Targeting PROTAC Activates AMPK and Inhibits Endoplasmic Reticulum Stress. Biomed. Pharmacother. 2023, 168, 115667. DOI: 10.1016/j.biopha.2023.115667

17. Jarusiewicz, J. A.; Yoshimura, S.; Mayasundari, A.; Actis, M.; Aggarwal, A.; McGowan, K.; Yang, L.; Li, Y.; Fu, X.; Mishra, V.; Heath, R.; Narina, S.; Pruett-Miller, S. M.; Nishiguchi, G.; Yang, J. J.; Rankovic, Z. Phenyl Dihydrouracil: An Alternative Cereblon Binder for PROTAC Design. ACS Med. Chem. Lett. 2023, 14 (2), 141–145. DOI: 10.1021/acsmedchemlett.2c00436

18. Min, J.; Mayasundari, A.; Keramatnia, F.; Jonchere, B.; Yang, S. W.; Jarusiewicz, J.; Actis, M.; Das, S.; Young, B.; Slavish, J.; Yang, L.; Li, Y.; Fu, X.; Garrett, S. H.; Yun, M. K.; Li, Z.; Nithianantham, S.; Chai, S.; Chen, T.; Shelat, A.; Lee, R. E.; Nishiguchi, G.; White, S. W.; Roussel, M. F.; Potts, P. R.; Fischer, M.; Rankovic, Z. Phenyl-Glutarimides: Alternative Cereblon Binders for the Design of PROTACs. Angew. Chem. Int. Ed. Engl. 2021, 60 (51), 26663–26670. DOI: 10.1002/anie.202108848

19. Hamilton, E. P.; Schott, A. F.; Nanda, R.; Lu, H.; Keung, C. F.; Gedrich, R.; Parameswaran, J.; Han, H. S.; Hurvitz, S. A. ARV-471, an Estrogen Receptor (ER) PROTAC degrader, Combined with Palbociclib in Advanced ER+/Human Epidermal Growth Factor Receptor 2–Negative (HER2^−^) Breast Cancer: Phase 1b Cohort (Part C) of a Phase 1/2 Study. J. Clin. Oncol. 2022, 40 (16_suppl), TPS1120–TPS1120. DOI: 10.1200/JCO.2022.40.16_suppl.TPS1120

20. Rose, T. E.; Morisseau, C.; Liu, J. Y.; Inceoglu, B.; Jones, P. D.; Sanborn, J. R.; Hammock, B. D. 1-Aryl-3-(1-acylpiperidin-4-yl)urea Inhibitors of Human and Murine Soluble Epoxide Hydrolase: Structure–Activity Relationships, Pharmacokinetics, and Reduction of Inflammatory Pain. J. Med. Chem. 2010, 53 (19), 7067– 7075. DOI: 10.1021/jm100691c

21. Jones, P. D.; Wolf, N. M.; Morisseau, C.; Whetstone, P.; Hock, B.; Hammock, B. D. Fluorescent Substrates for Soluble Epoxide Hydrolase and Application to Inhibition Studies. Anal. Biochem. 2005, 343 (1), 66–75. DOI: 10.1016/j.ab.2005.03.041

22. Morisseau, C.; Sahdeo, S.; Cortopassi, G.; Hammock, B. D. Development of an HTS Assay for EPHX2 Phosphatase Activity and Screening of Nontargeted Libraries. Anal. Biochem. 2013, 434 (1), 105–111. DOI: 10.1016/j.ab.2012.11.017

