## Supplemental Information for "*In vivo*–Active Soluble Epoxide Hydrolase–targeting PROTACs with Improved Potency and Stability"

### Table of contents

**Tables**

| | |
|---|---|
| **Table S1.** | **The mean and SD values in Table 1.** |
| **Table S2.** | **The mean and SD values in Table 2.** |
| **Table S3.** | **The mean and SD values of the mouse microsomal stability assay in Figure 2B.** |
| **Table S4.** | **The mean and SD values in Table 3.** |
| **Table S5.** | **The mean and SD values in Figure 6A.** |
| **Table S6.** | **List of antibodies used in immunoblotting assay for detection of ER stress.** |

**Figures**

| | |
|---|---|
| **Fig. S1.** | **Chemical stability of 1a and ALT-PG2.** |
| **Fig. S2.** | **Concentration–dependent degradation of sEH induced by compound 8 (0.01–1000 nM).** |
| **Fig. S3.** | **Concentration–dependent degradation of sEH induced by ALT-PG2.** |
| **Fig. S4.** | **Chemical stability of compounds 8, 14 - 19.** |
| **Fig. S5.** | **Immunoblot data in Table 3.** |
| **Fig. S6.** | **Concentration–dependent degradation of sEH induced by 17 and 19 (0.03–10 nM).** |
| **Fig. S7.** | **Degradation of sEH in HEK293T cells induced by sEH PROTACs.** |
| **Fig. S8.** | **Degradation of sEH in MDA-MB-231 and SH-SY5Y cells induced by compound 8.** |
| **Fig. S9.** | **Immunoblot data in Figure 5.** |
| **Fig. S10.** | **Chromatin condensation assay in HEK293T cells treated with compounds** |

**Experimental procedure**

**NMR spectra**

**HPLC traces**

**Reference**

**Table S1. The mean and SD values in Table 1.**

| | % sEH degradation | |
|---|---|---|
| Compound | **Mean** | **SD** |
| **1a** | 35 | 8.4 |
| **1** | 30 | 17 |
| **2** | -8 | 16 |
| **3** | -2 | 15 |
| **4** | 10 | 5.4 |
| **5** | 32 | 6.3 |
| **6** | 79 | 0.7 |
| **7** | 56 | 1.7 |
| **8** | 78 | 2.4 |

**Table S2. The mean and SD values in Table 2.**

| | % sEH degradation | |
|---|---|---|
| Compound | **Mean** | **SD** |
| **9** | 39 | 2.6 |
| **10** | 37 | 5.0 |
| **11** | 39 | 2.9 |
| **12** | 32 | 13 |
| **13** | 50 | 9.8 |

**Table S3. The mean and SD values of the mouse microsomal stability assay in Figure 2B.**

| Compound | Time (min) | Area ratio #1 | Area ratio #2 | % Normalized value #1 | % Normalized value #2 | % Mean of normalized value | SD |
|---|---|---|---|---|---|---|---|
| Ketanserin | 0 | 2.95 | 3.04 | 98.5 | 102 | 100 | 1.50 |
| | 5 | 2.32 | 2.35 | 77.5 | 78.5 | 78.0 | 0.501 |
| | 15 | 1.41 | 1.46 | 47.1 | 48.7 | 47.9 | 0.835 |
| | 30 | 0.720 | 0.728 | 24.0 | 24.3 | 24.2 | 0.134 |
| | 45 | 0.331 | 0.438 | 11.1 | 14.6 | 12.8 | 1.79 |
| | 45 (without NADPH) | 3.34 | 3.39 | 112 | 113 | 112 | 0.835 |
| **1a** | 0 | 1.03 | 1.00 | 102 | 98.5 | 100 | 1.48 |
| | 5 | 0.003 | 0.003 | 0.3 | 0.3 | 0.3 | 0 |
| | 15 | 0.001 | 0.001 | 0.1 | 0.1 | 0.1 | 0 |
| | 30 | 0.001 | 0.002 | 0.1 | 0.2 | 0.1 | 0.0182 |
| | 45 | 0.002 | 0.003 | 0.2 | 0.3 | 0.2 | 0.0764 |
| | 45 (without NADPH) | 0.925 | 0.911 | 91.1 | 89.8 | 90.4 | 0.690 |
| **6** | 0 | 0.041 | 0.059 | 82.2 | 118 | 100 | 17.8 |
| | 5 | 0.052 | 0.049 | 105 | 98.7 | 102 | 2.92 |
| | 15 | 0.044 | 0.042 | 88.2 | 83.6 | 85.9 | 2.32 |
| | 30 | 0.028 | 0.028 | 56.6 | 56.2 | 56.4 | 0.201 |
| | 45 | 0.019 | 0.020 | 38.3 | 40.3 | 39.3 | 1.01 |
| | 45 (without NADPH) | 0.042 | 0.069 | 85.2 | 138 | 112 | 26.6 |
| **8** | 0 | 0.081 | 0.074 | 104 | 95.6 | 100.0 | 4.40 |
| | 5 | 0.085 | 0.087 | 110 | 113 | 111.1 | 1.68 |
| | 15 | 0.086 | 0.089 | 112 | 116 | 114 | 2.01 |
| | 30 | 0.074 | 0.075 | 95.7 | 96.5 | 96.1 | 0.389 |
| | 45 | 0.066 | 0.062 | 85.1 | 79.7 | 82.4 | 2.72 |
| | 45 (without NADPH) | 0.084 | 0.074 | 109 | 95.3 | 102 | 6.74 |

**Table S4. The mean and SD values in Table 3.**

| | sEH degradation | | | | Cytotoxicity | |
|---|---|---|---|---|---|---|
| | ELISA (1 µM) | | ELISA (100 nM) | | | |
| Compound | Mean (%) | SD | Mean (%) | SD | Mean (%) | SD |
| **8** | 71 | 14 | 31 | 8.6 | -38 | 16 |
| **14** | 52 | 7.6 | 33 | 5.9 | -24 | 2.7 |
| **15** | 48 | 6.1 | -2 | 8.7 | -47 | 46 |
| **16** | 39 | 9.0 | 29 | 7.5 | -6 | 20 |
| **17** | 75 | 1.5 | 51 | 2.9 | 7 | 5.0 |
| **18** | 56 | 6.2 | 43 | 2.8 | 46 | 5.5 |

**Table S5. The mean and SD values in Figure 6A.**

| Compound | Sampling time (h) | Concentration (ng/mL) | | | Mean (ng/mL) | SD |
|---|---|---|---|---|---|---|
| | | Mouse #1 | Mouse #2 | Mouse #3 | | |
| **8** | 0.25 | 2510 | 1410 | 1330 | 1750 | 660 |
| | 0.5 | 2740 | 1590 | 1310 | 1880 | 760 |
| | 1 | 2460 | 1540 | 1470 | 1820 | 551 |
| | 2 | 3620 | 1380 | 1840 | 2280 | 1180 |
| | 4 | 4170 | 1700 | 1930 | 2600 | 1370 |
| | 6 | 3510 | 1810 | 1840 | 2390 | 976 |
| | 8 | 2550 | 2170 | 2070 | 2260 | 255 |
| | 24 | 1090 | 903 | 531 | 839 | 282 |
| **17** | 0.25 | 3690 | 3290 | 3000 | 3320 | 345 |
| | 0.5 | 4180 | 3050 | 2440 | 3220 | 885 |
| | 1 | 3020 | 2200 | 2150 | 2460 | 487 |
| | 2 | 3000 | 1820 | 1990 | 2270 | 640 |
| | 4 | 2390 | 1700 | 1670 | 1920 | 407 |
| | 6 | 1850 | 1450 | 1330 | 1540 | 276 |
| | 8 | 1170 | 1490 | 1290 | 1320 | 159 |
| | 24 | 344 | 304 | 180 | 276 | 86.0 |
| **19** | 0.25 | 3150 | 2770 | 2360 | 2760 | 397 |
| | 0.5 | 3740 | 2660 | 2180 | 2860 | 800 |
| | 1 | 1800 | 1680 | 1250 | 1580 | 290 |
| | 2 | 808 | 670 | 543 | 673 | 132 |
| | 4 | 305 | 296 | 239 | 280 | 35.9 |
| | 6 | 171 | 164 | 146 | 160 | 12.8 |
| | 8 | 87.6 | 135 | 116 | 113 | 24.1 |
| | 24 | 15.1 | 23.2 | 14.5 | 17.6 | 4.88 |

**Table S6. List of antibodies used in immunoblotting assay for detection of ER stress.**

| Antibodies | Source | Catalog Number | Observed MW (kDa) | Host | Dilution |
|---|---|---|---|---|---|
| eIF2α | Cell Signaling Technology | 5324 | 40 | Rabbit | 1:1,000 |
| IRE1α | Cell Signaling Technology | 3294 | 115 | Rabbit | 1:1,000 |
| PERK | Cell Signaling Technology | 3192 | 140 | Rabbit | 1:1,000 |
| Phospho-eIF2α$^{S751}$ | Cell Signaling Technology | 3398 | 40 | Rabbit | 1:1,000 |
| Phospho-IRE1α$^{S724}$ | Abcam | ab 48187 | 115 | Rabbit | 1:10,000 |
| Phospho-PERK$^{T980}$ | Santa Cruz Biotechnology | sc-32577 | 160 | Rabbit | 1:1,000 |
| sEH | Santa Cruz Biotechnology | sc-166961 | 61 | Mouse | 1:500 |
| β-actin | Santa Cruz Biotechnology | sc-47778 | 44 | Mouse | 1:20,000 |

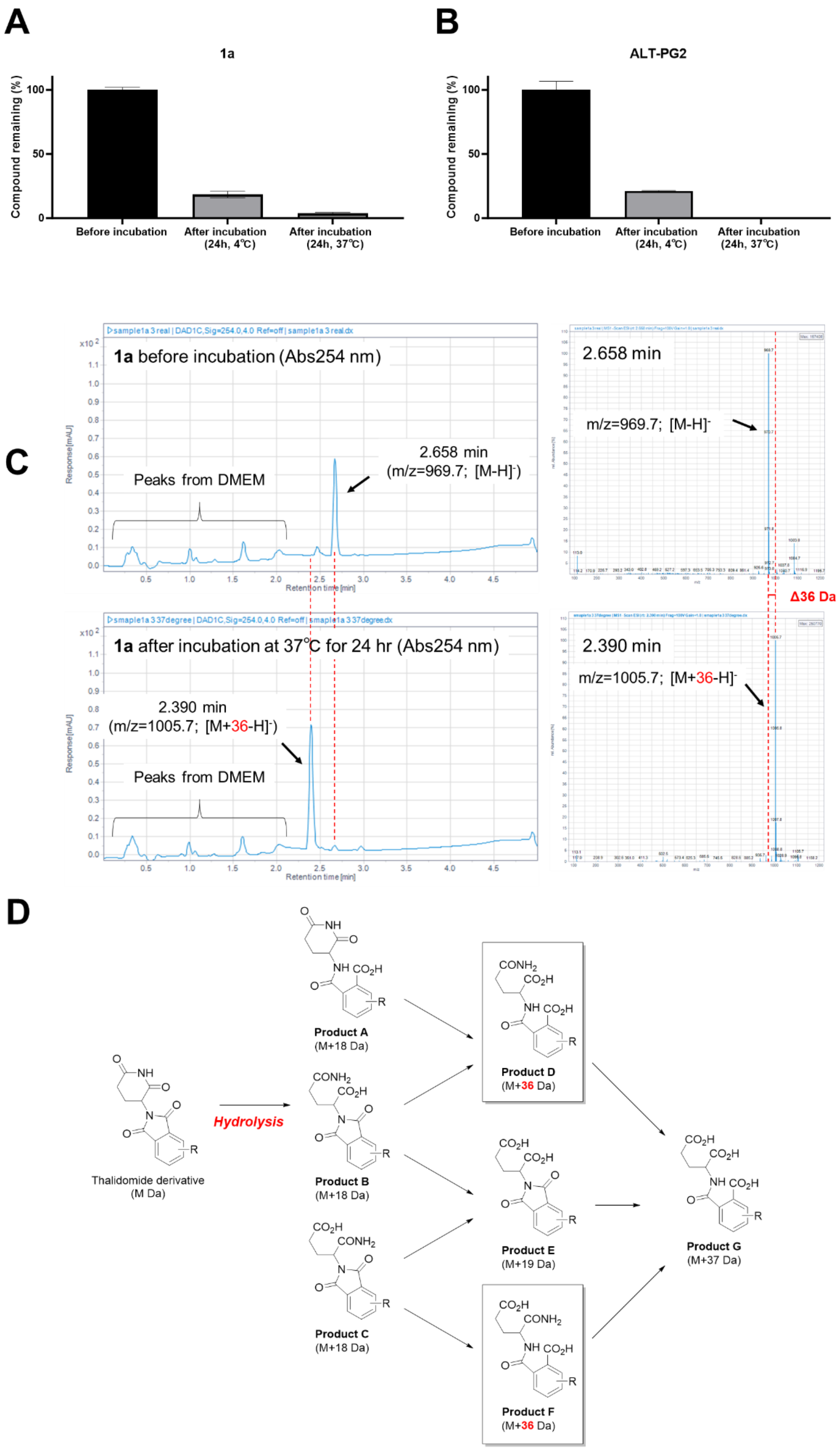


**Figure S1. Chemical stability of 1a and ALT-PG2.** (A) Stability of sEH-PROTACs **1a** and **ALT-PG2** in cell culture medium (DMEM). Data are the mean ± SD of three individual experiments. The absorbance of 254 nm of compound solutions was measured by LC-MS using a $H_2O$-MeCN gradient (100:0 to 5:95, v/v, 0.1% Formic acid (FA)). (B) Stability of **ALT-PG2** in the cell culture medium. (C) Chromatogram of **1a** in the cell culture medium. (D) Predicted hydrolysis structure of E3 recruiter.[1]

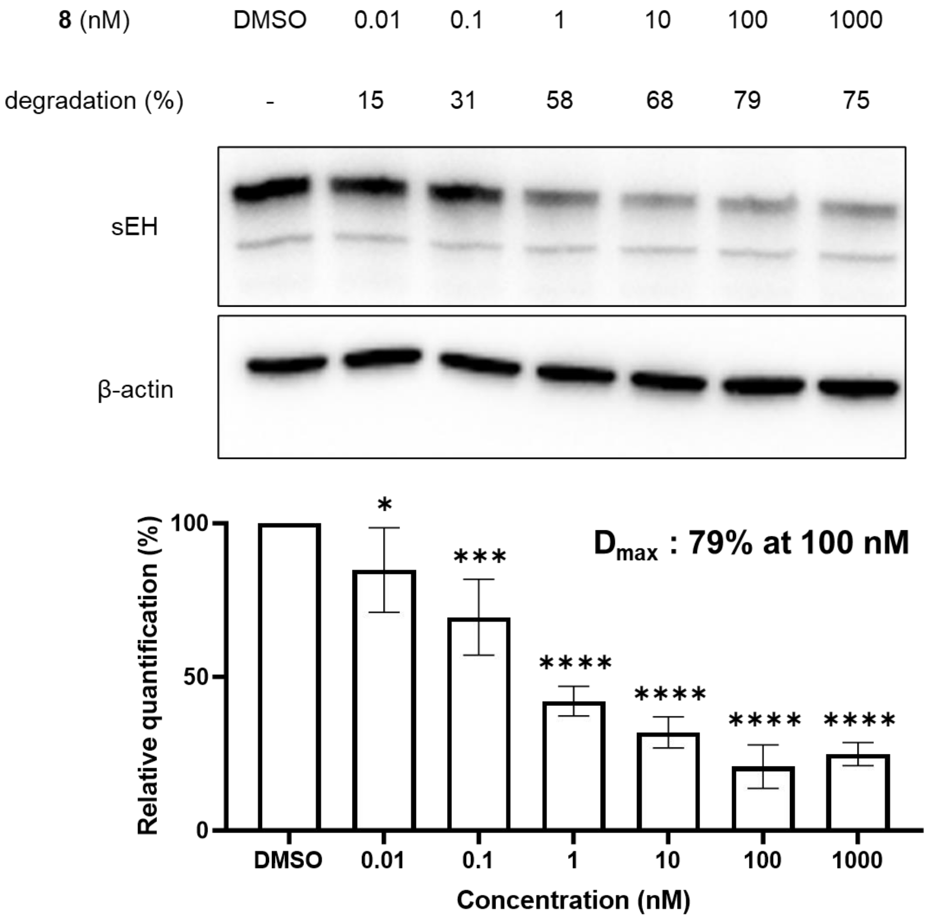


**Figure S2. Concentration–dependent degradation of sEH induced by compound 8 (0.01–1000 nM).** HepG2 cells were treated with the indicated concentration of **8** for 24 hours and sEH levels in whole cell lysate were measured by immunoblot. Data are the mean ± SD of three individual experiments. The difference between treatment groups was analyzed by one-way ANOVA with the Holm–Sidak test (****$p < 0.0001$, *** $p < 0.001$, *$p < 0.04$).

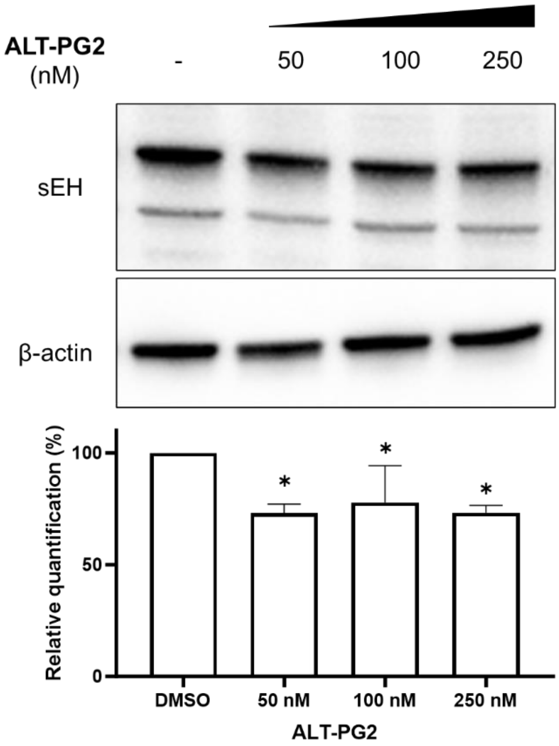


**Figure S3. Concentration–dependent degradation of sEH induced by ALT-PG2.** HepG2 cells were treated with the indicated concentration of **ALT-PG2** at 50–250 nM for 24 hours, and sEH levels in whole cell lysate were measured by immunoblot. Data are the mean ± SD of three individual experiments. The difference between treatment groups was analyzed by one-way ANOVA with the Holm–Sidak test (*$p < 0.04$).

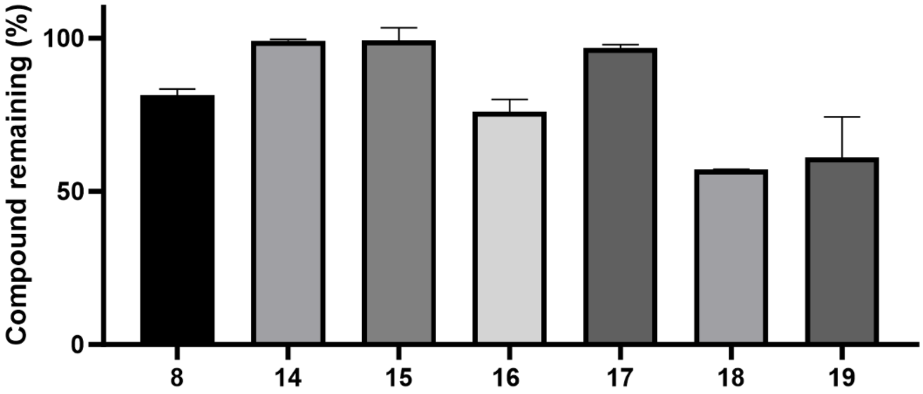


**Figure S4. Chemical stability of compounds 8, 14–19.** Each compound (100 μM) was incubated in DMEM at 37 °C for 24 hours. Mean ± SD of three individual experiments is shown.

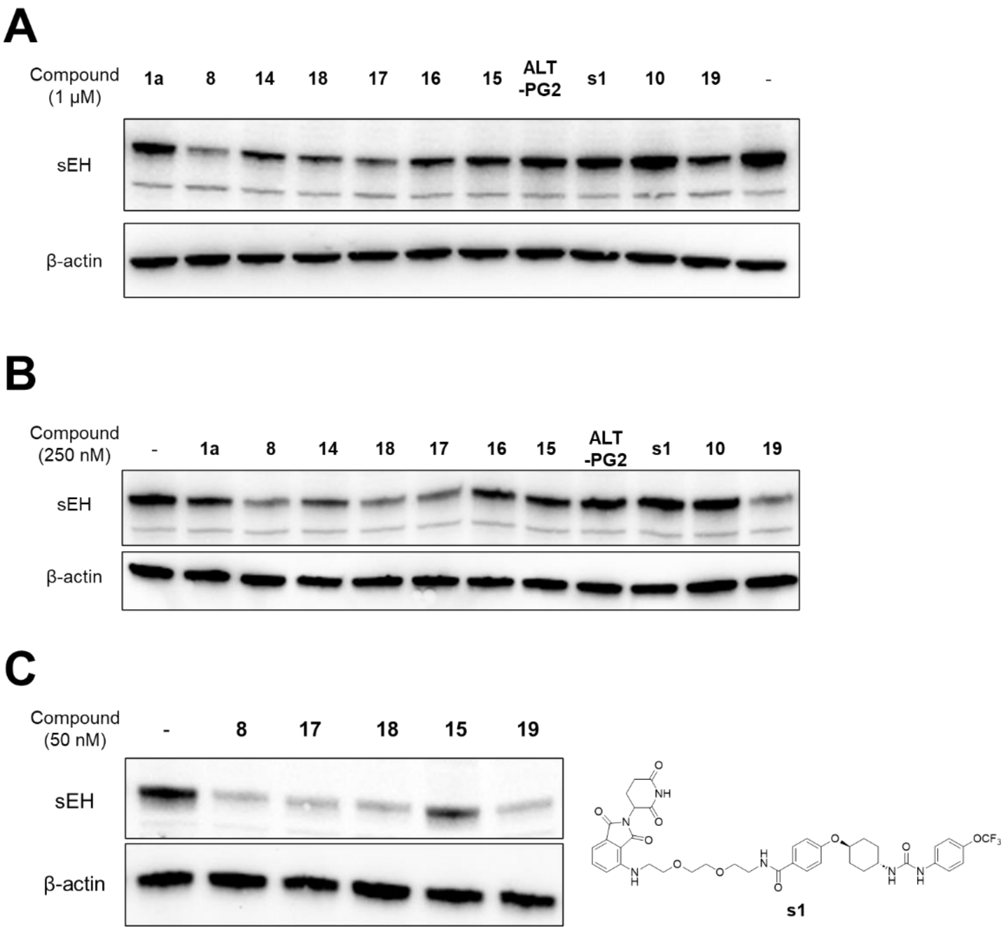


**Figure S5. Immunoblot data in Table 3.** HepG2 cells were treated with the indicated compound in triplicates at 1 μM (A), 250 nM (B), or 50 nM (C) for 24 hours, and the sEH level was measured by immunoblot.

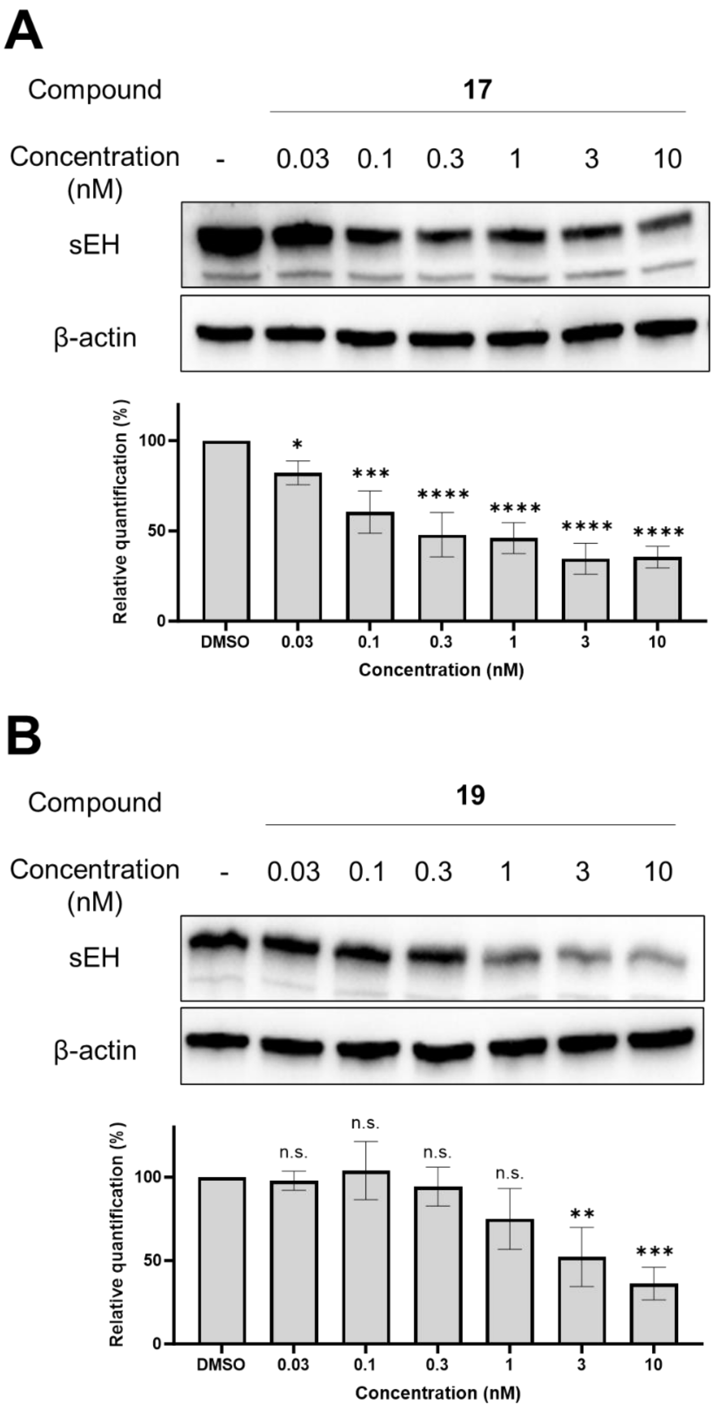


**Figure S6. Concentration–dependent degradation of sEH induced by 17 and 19 (0.03–10 nM).** HepG2 cells were treated with **17** or **19** at the indicated concentration for 24 hours and sEH levels were measured by immunoblot. Data are the mean ± SD of three individual experiments. The difference between treatment groups was analyzed by one-way ANOVA with the Holm–Sidak test (****$p < 0.0001$, *** $p < 0.001$, ** $p < 0.01$, *$p < 0.04$).

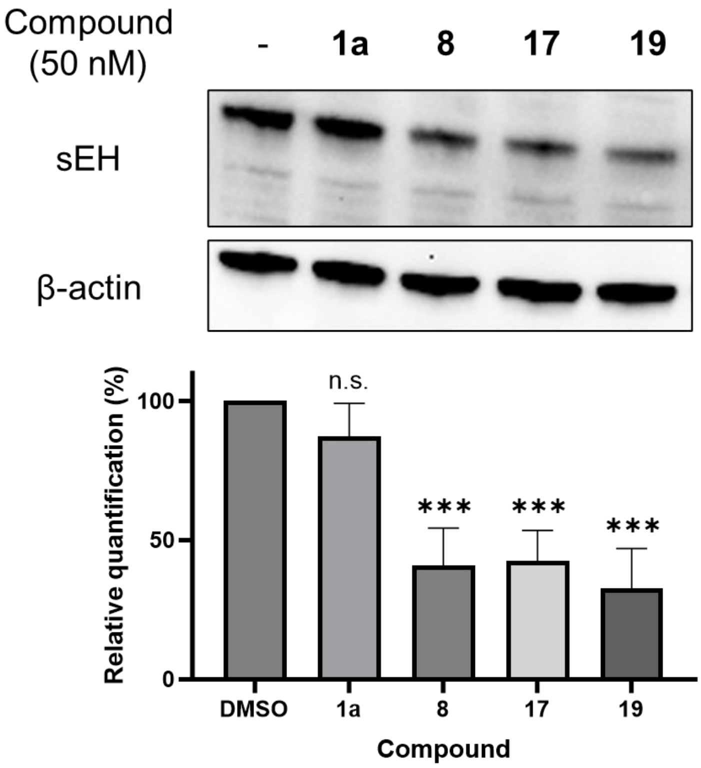


**Figure S7. Degradation of sEH in HEK293T cells induced by sEH PROTACs.** HEK293T cells were treated with the indicated compound at 50 nM for 24 hours, and the sEH level was measured by immunoblot. Data are the mean ± SD of three individual experiments. The difference between treatment groups was analyzed by one-way ANOVA with the Holm–Sidak test (*** $p < 0.001$).

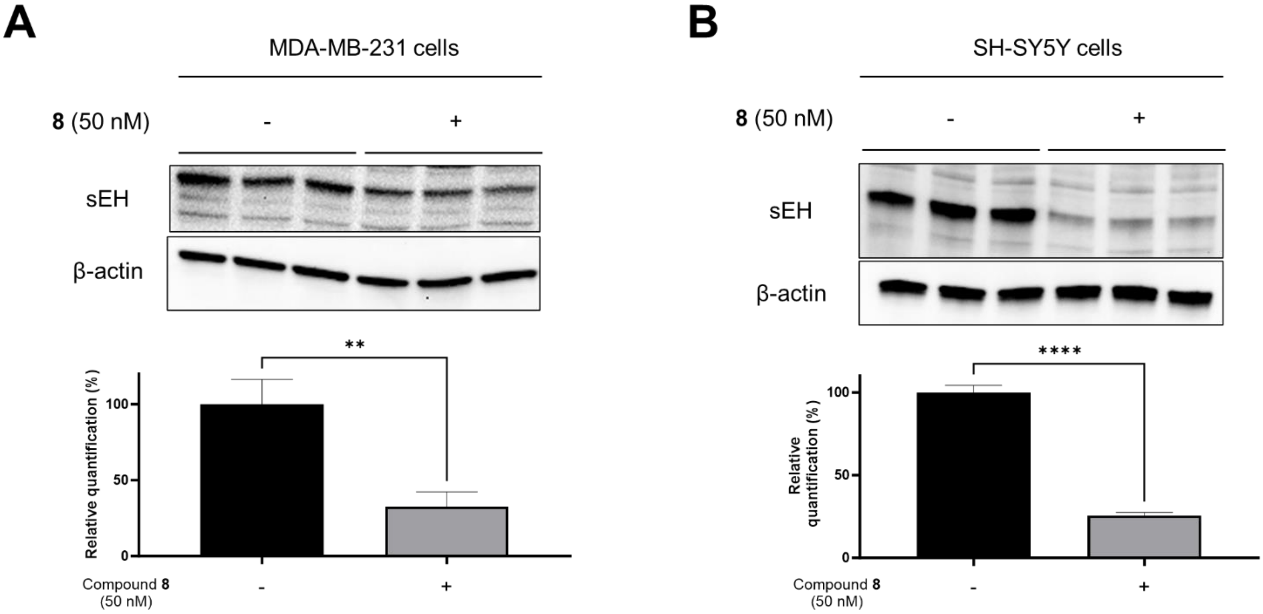


**Figure S8. Degradation of sEH in MDA-MB-231 and SH-SY5Y cells induced by compound 8.** (A) MDA-MB-231 and (B) SH-SY5Y cells were treated with compound **8** at 50 nM for 24 hours, and the sEH level was measured by immunoblot. Data are the mean ± SD of three individual experiments. The difference between treatment groups was analyzed by Student's t-test (****$p < 0.0001$, ** $p < 0.01$).

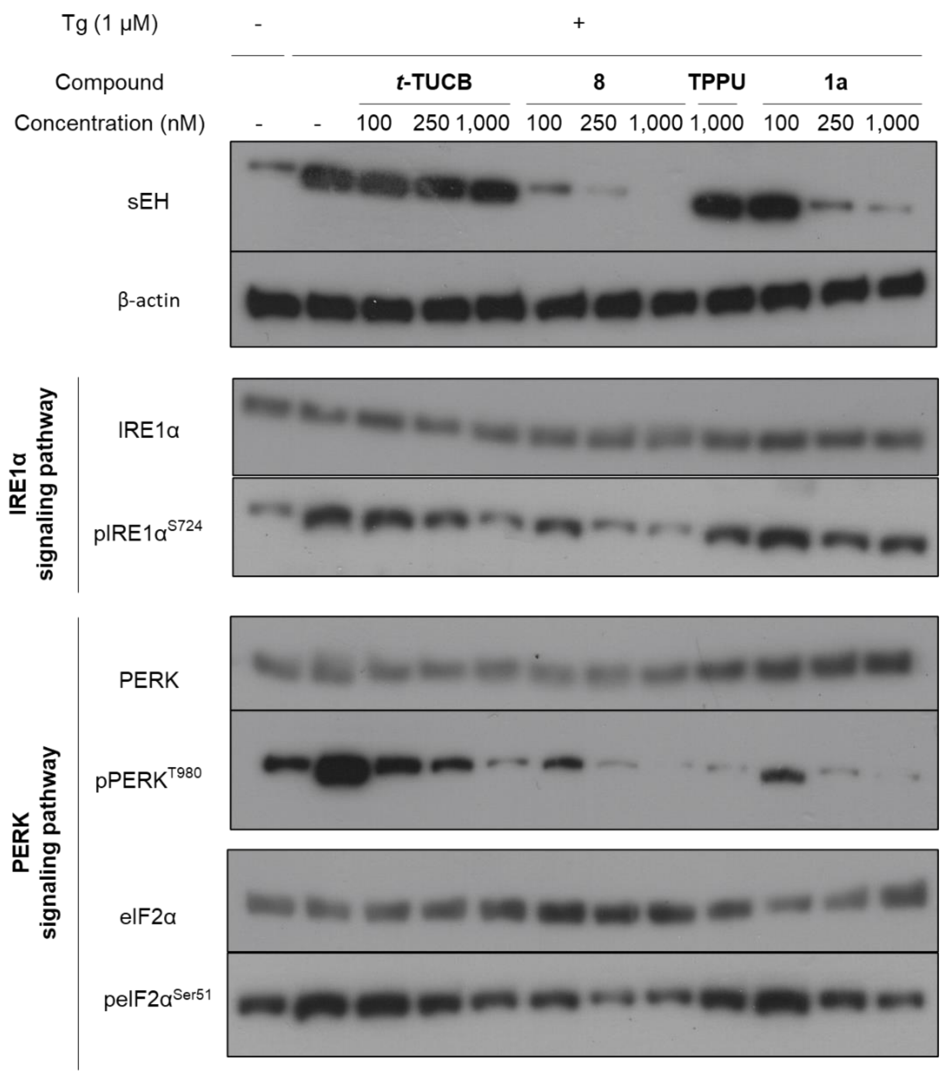


**Figure S9. Immunoblot data in Figure 5.** HEK293T cells were pre-treated with the indicated sEH modulators prior to adding 1 μM of Tg for an additional 24 hours ($n = 4$).

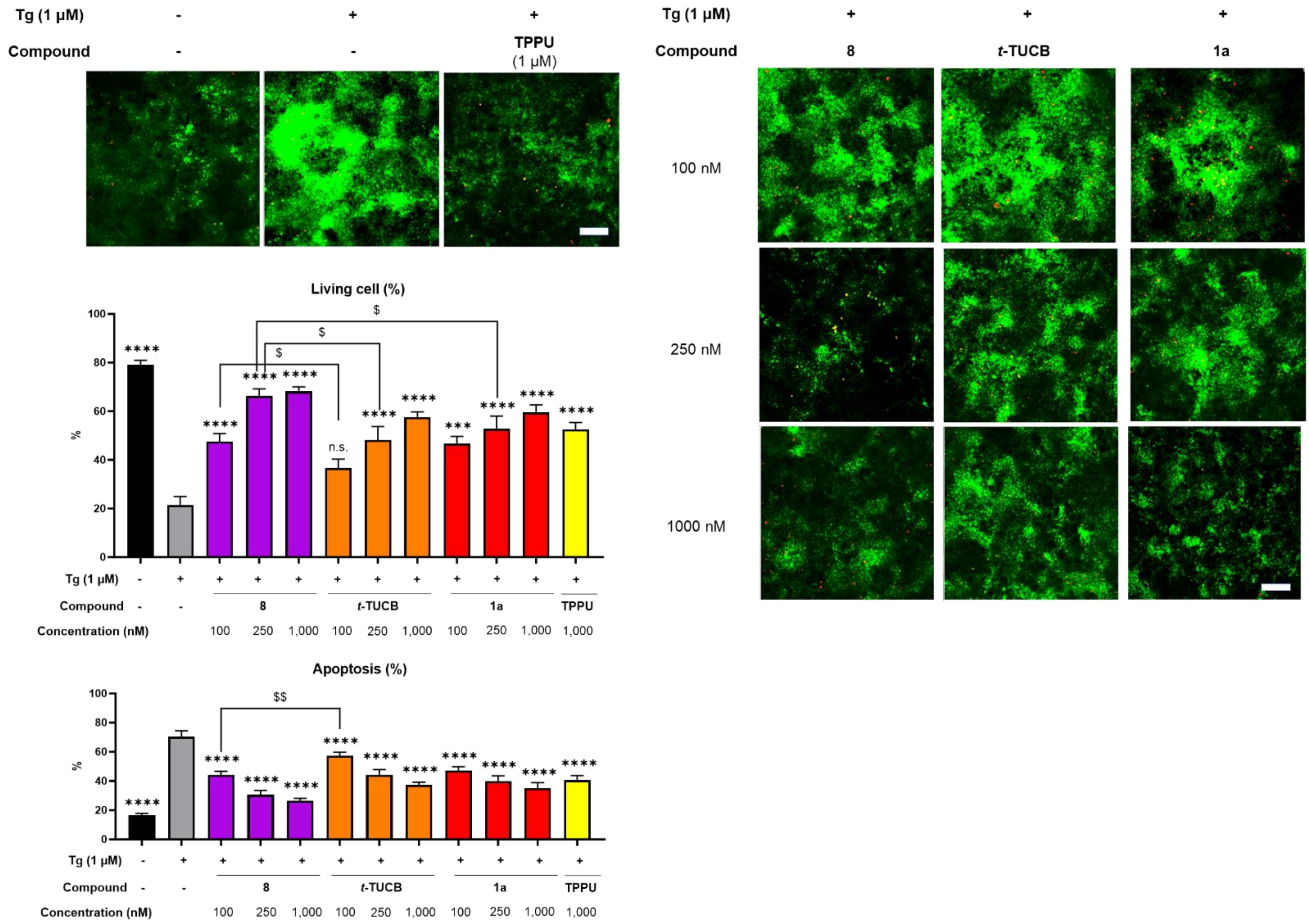


**Figure S10. Chromatin condensation assay in HEK293T cells treated with compounds.** HEK293T cells were pre-treated with the indicated compounds for 12 hours prior to exposure to 1 µM of Tg and compounds for an additional 24 hours. Cells were then stained with Hoechst 33258 (50 µg/mL in PBS; green fluorescence) and propidium iodide (10 µg/mL in PBS; Red fluorescence) and visualized by fluorescence microscopy using Leica DMI8 (Leica Microsystems Inc. Buffalo Grove, IL). Cells with increased chromatin condensation indicative of early stages of apoptosis are labelled in green fluorescence, while necrotic cells are labeled with red fluorescence. Orange cells on the other hand represent late apoptosis and are both Hoechst and propidium iodide positive. Live cells display weak green fluorescence. Images are representative of at least 4 independent experiments. For each condition, at least 500 cells were counted. Scale bar is 100 µm. Percentages of apoptotic and necrotic cells were calculated relative to total number of cells per view and are presented as Mean ± SEM ($n$ = 6) values are shown. The difference between treatment groups was analyzed by one-way ANOVA with the Holm–Sidak test (****$p < 0.0001$, ***$p < 0.001$) and by the Student's t-test ($^{\$\$}p < 0.009$, $^{\$}p < 0.04$).

**Abbreviations used**

ANOVA, analysis of variance; AUC, area under the curve; BAT, brown adipose tissue; $CL_{int}$, intrinsic clearance; $C_{max}$, maximum concentration; COPD, chronic obstructive pulmonary disease; $DC_{50}$, half-maximal degradation concentration; DIEPA, *N*,*N*-diisopropylamine; $D_{max}$, maximal level of degradation; DMEM, Dulbecco's modified Eagle medium; DMF, *N*,*N*-dimethylformamide; DMSO, dimethyl sulfoxide; eIF2α, eukaryotic translation initiation factor 2 alpha; EpFA, epoxy fatty acid; ER, Endoplasmic reticulum; HATU, 1-[bis(dimethylamino)methylene]-1H-1,2,3-triazolo[4,5-b]pyridinium 3-oxide hexafluorophosphate; i.p., intraperitoneal; $IC_{50}$, half-maximal inhibitory concentration; inf, to infinity; Int, intermediate; IRE1α, inositol-requiring enzyme 1 alpha; $k_{cat}$, turnover number; $K_M$, Michaelis–Menten constant; MRT, mean residence time; n.s., not significant; NADPH, nicotinamide adenine dinucleotide phosphate; NEDD8, neural precursor cell expressed developmentally down-regulated protein 8; PDB, protein data bank; PERK, protein kinase RNA-like ER kinase; PK, pharmacokinetics; PROTACs, proteolysis targeting chimera; RT, room temperature; SD, standard deviation; sEH, soluble epoxide hydrolase; sEHi, sEH inhibitor; SEM, standard error of the mean; $T_{1/2}$, half-life; Tg, thapsigargin; $T_{max}$, time of maximum concentration; Veh, vehicle

**Experimental procedure**

**Chemistry experiments**

***General***

NMR spectra were recorded on a Bruker biospin AVANCE III 300 (300 MHz for $^1H$, 75 MHz for $^{13}C$, 282 MHz for $^{19}F$) and a Bruker biospin AVANCE III HD 600 (600 MHz for $^1H$, 151 MHz for $^{13}C$, 565 MHz for $^{19}F$) instrument in the indicated solvent. Chemical shifts are reported in units, parts per million (ppm) relative to the signal (0.00 ppm) for internal tetramethylsilane for solutions in $CDCl_3$ (7.26 ppm for $^1H$, 77.16 ppm for $^{13}C$), $CD_3OD$ (3.31 ppm for $^1H$, 49.00 ppm for $^{13}C$), or DMSO-$d_6$ (2.50 ppm for $^1H$, 39.52 ppm for $^{13}C$).[2] $CFCl_3$ (0.65 ppm in $CDCl_3$, -1.67 ppm in $CD_3OD$, -0.24 ppm in DMSO-$d_6$) or TFA (-75.39 ppm in $CDCl_3$, -77.77 ppm in $CD_3OD$, -74.95 ppm in DMSO-$d_6$) were used as an internal standard for $^{19}F$-NMR.[3] Multiplicities are reported using the following abbreviations: s; singlet, d; doublet, t; triplet, m; multiplet, br; broad, *J*; coupling constants in Hertz. High-resolution mass spectra (HRMS) were recorded on Orbitrap Exploris 480 Mass Spectrometer (ThermoFisher Scientific). Reactions were monitored on TLC plates (silica gel 60, F254 coating, Sigma Aldrich), and spots were monitored under UV light (254 mm). The same TLC system was used to test purity, and all final products showed a single spot on TLC with both $KMnO_4$ and UV absorbance. CombiFlash NextGen 300+ (Teledyne ISCO) was used for normal-phase column chromatography. The purity of the compounds that were tested in the assay was >95% based on $^1H$ NMR and reverse phase high-performance liquid chromatography (HPLC)-UV (Agilent LC-MSD system with 1290MCT column oven, 1290 Multisampler, 1290 High Speed Pump, 1290 DAD FS, LC/MSD, reverse phase HPLC column Agilent InfinityLab Poroshell 120 EC-C18 (2.1 x 50 mm, particle size 1.9 μm)) on monitoring absorption at 254 nm. The HPLC gradient method consisted of an aqueous phase (Milli-Q water with 0.1% formic acid (FA)) and an organic phase (acetonitrile (MeCN) with 0.1% FA) with a 0.50 mL/min flow. The first step consisted of 95% aqueous and 5% organic phases followed by a 4.0 min gradient to 100% organic phase. A subsequent 30-second step of 100% organic phase was followed by 30-second gradient to 95% aqueous and 5% organic phase. Preparative HPLC was performed with Waters2545 Binary Gradient Module, Waters515 HPLC Pump, Waters SFO System Fluidics organizer, Waters2489 UV/Visible Detector, Waters2424 ELS Detector, Waters QDa Detector, and Waters3767 Sample Manager using a C18 reverse phase column (Waters, waters X Bridge BEH C18 OBD Prep Column, 19 x 250 mm, 5 μm).

***Materials***

All reagents and solvents were purchased from commercial suppliers and were used without further purification. 4-Amino-1-boc-piperidine was purchased from 1PlusChem. Pomalidomide-piperadine was purchased from AA Blocks Inc. 9-((tert-butoxycarbonyl)amino)nonanoic acid and tert-butyl (14-amino-3,6,9,12-Tetraoxatetradecyl)carbamate were purchased from AmBeed. 1-[Bis(dimethylamino)methylene] -1*H*-1,2,3-triazolo[4,5-*b*]pyridinium 3-oxid hexafluorophosphate (HATU) was purchased from Combi-Blocks. 4-(Trifluoromethoxy)phenyl isocyanate was purchased from Enamine. Formic acid (FA) , MeCN, and water were purchased from Fisher Chemical. Thalidomide-PEG3-amine hydrochloride was purchased from MedChemExpress. Dichloromethane ($CH_2Cl_2$), Hoechst 33258, *N*,*N*-diisopropylethylamine (DIEPA), *N*,*N*-dimethylformamide (DMF), *N*-acetyl-*S*-(2-amino-9-(4-fluorobenzyl)-6-oxo-6,9-dihydro-1*H*-purin-8-yl)-L-cysteine, Pomalidomide-piperazine-C1-4-piperidine hydrochloride, Propidium iodide, and Triethyl amine (TFA) were all purchased from Millipore-Sigma. Lenalidomide 4'-PEG3-amine, lenalidomide 5'-piperazine, lenalidomide 5'-piperazine-4-methylpiperidine, PD 4'-oxyacetic acid, phenyl-glutarimide 4'-oxyacetic acid, pomalidomide 4'-4-PEG4-amine, pomalidomide 4'-alkylC5-amine, pomalidomide 4'-PEG2-acid, pomalidomide 4'-PEG4-amine, pomalidomide 5'-fluoro-6'-piperazine, pomalidomide 5'-fluoro-6'-piperazine-4-methylpiperidine, pomalidomide 5'-piperazine, pomalidomide 5'-piperazine-4-methylpiperidine, tDHU acid, VH 032 amide-alkylC6-acid, and VH 032 amide-PEG4-acid were purchased from Tocris Bioscience. Compounds **1a** and *t*-TUCB (**Int 1**) were prepared as were prepared as previously reported.[4, 5]

***Synthesis procedure for PROTAC molecules***

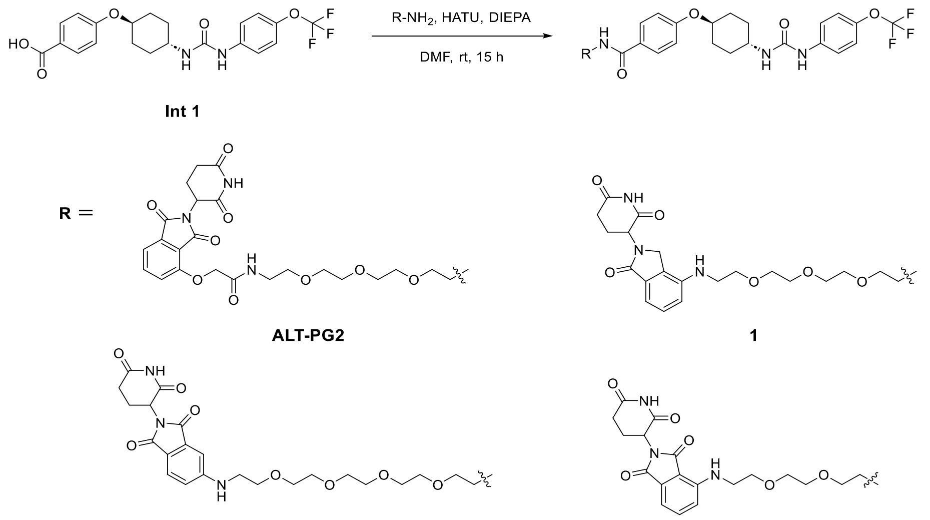


**Synthesis of ALT-PG2,[6] compounds 1, 2, and s1**

**Int 1** (10.5 mg, 24.0 μmol, 1.0 equiv.) and HTAU (18.3 mg, 48.0 mmol, 2.0 equiv.) were added to a solution of amine derivative (24.0 μmol, 1.0 equiv.) in DMF (500 μL) and DIEPA (50 μL, 287 μmol, 12.0 equiv.). The reaction mixture was stirred at room temperature for 15 hours, then filtered through a PTFE membrane and purified via preparative HPLC using $H_2O$-MeCN gradient (99: 1 to 5: 95, v/v, 0.1% FA). A fraction containing the target molecule was lyophilized to give a solid.

**ALT-PG2** (16.2 mg, 77 % yield). $^1H$ NMR (600 MHz, DMSO-$d_6$) $\delta_H$ 11.11 (br, 1H), 8.74 (br, 1H), 8.32 (t, $J$ = 5.6 Hz, 1H), 8.00 (t, $J$ = 5.6 Hz, 1H), 7.83-7.75 (m, 3H), 7.51-7.45 (m, 3H), 7.39 (d, $J$ = 8.5 Hz, 2H), 7.21 (d, $J$ = 8.6 Hz, 2H), 6.99 (d, $J$ =

8.9 Hz, 2H), 6.41 (d, $J$ = 7.4 Hz, 1H), 5.11 (dd, $J$ = 12.9 Hz, 5.4 Hz, 1H), 4.78 (s, 2H), 4.48-4.39 (m, 1H), 3.57-3.47 (m, 11H), 3.45 (t, $J$ = 5.6 Hz, 2H), 2.94-2.85 (m, 1H), 2.64-2.51 (m, 2H), 2.08-2.01 (m, 3H), 1.96-1.89 (m, 2H), 1.52-1.43 (m, 2H), 1.41-1.32 (m, 2H). $^{19}$F NMR (565 MHz, DMSO-$d_6$) $\delta_F$ -56.94. Purity (HPLC-UV 254 nm): >95%. ($^t$R= 2.716 min). White powder.

Compound **1** (18.1 mg, 88 % yield). $^1$H NMR (600 MHz, $CD_3OD$) $\delta_H$ 7.78-7.74 (m, 2H), 7.45-7.42 (m, 2H), 7.34 (t, $J$ = 7.8 Hz, 1H), 7.16 (d, $J$ = 8.4 Hz, 2H), 7.13 (d, $J$ = 7.4 Hz, 1H), 6.95-6.91 (m, 2H), 6.88 (d, $J$ = 8.0 Hz, 1H), 5.14 (dd, $J$ = 13.4 Hz, 5.2 Hz, 1H), 4.37-4.31 (m, 1H), 4.29 (d, $J$ = 4.0 Hz, 2H), 3.68-3.59 (m, 13H), 3.53 (t, $J$ = 5.5 Hz, 2H), 3.39 (t, $J$ = 5.6 Hz, 2H), 2.93-2.85 (m, 1H), 2.80-2.74 (m, 1H), 2.49-2.40 (m, 1H), 2.19-2.01 (m, 5H), 1.60-1.51 (m, 2H), 1.44-1.35 (m, 2H). $^{13}$C NMR (151 MHz, $CD_3OD$) $\delta_C$ 174.67, 172.34, 172.27, 169.82, 162.02, 157.34, 145.03, 144.58, 140.24, 133.13, 130.67, 130.23, 128.58, 127.51, 122.68, 120.89, 116.31, 114.62, 112.80, 75.90, 71.60, 71.57, 71.38, 71.28, 70.61, 70.43, [72-70 ppm region several PEG peaks overlap], 53.58, 49.57, 47.30, 44.37, 40.88, 32.38, 31.40, 31.11, 24.27. $^{19}$F NMR (565 MHz, $CD_3OD$) $\delta_F$ -60.08. HRMS (ESI, positive): calcd. for $[C_{42}H_{49}N_6O_{10}F_3+H]^+$ 855.3541, found 855.3550. Purity (HPLC-UV 254 nm): >95%. ($^t$R= 2.692 min). White powder.

Compound **2** (20.3 mg, 93 % yield). $^1$H NMR (600 MHz, $CD_3OD$) $\delta_H$ 7.79-7.76 (m, 2H), 7.54 (dd, $J$ = 8.6 Hz, 7.1 Hz, 1H), 7.45-7.41 (m, 2H), 7.16 (d, $J$ = 8.4 Hz, 2H), 7.07 (d, $J$ = 8.5 Hz, 1H), 7.05 (d, $J$ = 6.9 Hz, 1H), 6.99-6.93 (m, 2H), 5.04 (dd, $J$ = 12.6 Hz, 5.5 Hz, 2H), 4.42-4.35 (m, 1H), 3.68 (t, $J$ = 5.2 Hz, 1H), 3.66-3.59 (m, 15H), 3.54 (t, $J$ = 5.5 Hz, 1H), 3.47 (t, $J$ = 5.3 Hz, 1H), 2.88-2.80 (m, 1H), 2.77-2.66 (m, 2H), 2.18-2.02 (m, 5H), 1.62-1.53 (m, 2H), 1.47-1.37 (m, 2H). $^{13}$C NMR (151 MHz, $CD_3OD$) $\delta_C$ 174.65, 171.60, 170.67, 169.82, 169.28, 162.01, 152.33, 148.22, 145.03, 140.23, 137.21, 133.89, 130.27, 127.55, 122.67, 120.88, 118.26, 116.31, 112.02, 111.30, 75.91, 71.65, 71.60, 71.34, 71.30, 70.61, [72-70 ppm region several PEG peaks overlap], 50.19, 49.67, 43.24, 40.93, 32.20, 31.42, 31.14, 23.82. $^{19}$F NMR (565 MHz, $CD_3OD$) $\delta_F$ -60.07. HRMS (ESI, positive): calcd. for $[C_{44}H_{51}N_6O_{12}F_3+H]^+$ 913.3595, found 913.3606; calcd. for $[C_{44}H_{51}N_6O_{12}F_3+Na]^+$ 935.3415, found 935.3425. Purity (HPLC-UV 254 nm): >95%. ($^t$R= 2.884 min). Yellow powder.

Compound **s1** (16.0 mg, 81 % yield). $^1$H NMR (300 MHz, DMSO-$d_6$) $\delta_H$ 11.09 (br, 1H), 8.82 (br, 1H), 8.30 (t, $J$ = 5.6 Hz, 1H), 7.78 (d, $J$ = 8.8 Hz, 2H), 7.62-7.53 (m, 1H), 7.49 (t, $J$ = 9.1 Hz, 1H), 7.20 (d, $J$ = 8.5 Hz, 2H), 7.11 (d, $J$ = 8.6 Hz, 1H), 7.03 (d, $J$ = 7.0, 1H), 6.97 (d, $J$ = 8.9 Hz, 2H), 6.59 (t, $J$ = 5.9 Hz, 1H), 6.52-6.45 (m, 1H), 5.05 (dd, $J$ = 12.6 Hz, 5.4 Hz, 2H), 4.48-4.33 (m, 1H), 3.65-3.38 (m, 13H), 2.96-2.79 (m, 1H), 2.64-2.52 (m, 1H), 2.10-1.86 (m, 5H), 1.56-1.26 (m, 4H). $^{13}$C NMR (151 MHz, DMSO-$d_6$) $\delta_C$ 172.79, 170.07, 168.92, 167.28, 165.72, 159.65, 154.51, 146.38, 141.86, 139.98, 136.20, 132.08, 128.98, 126.39, 121.57, 118.49, 117.40, 114.81, 110.65, 109.24, 74.12, 69.66, 69.64, 69.03, 68.87, 48.55, 47.14, 41.69, 40.05, 30.97, 29.96, 29.65, 22.12. $^{19}$F NMR (565 MHz, DMSO-$d_6$) $\delta_F$ -57.00. HRMS (ESI, positive): calcd. for $[C_{40}H_{43}F_3N_6O_{10}+H]^+$ 825.3071, found 825.3092; calcd. for $[C_{40}H_{43}F_3N_6O_{10}+Na]^+$ 847.2890, found 847.2906. Purity (HPLC-UV 254 nm): >95%. ($^t$R= 2.941 min). Yellow powder.

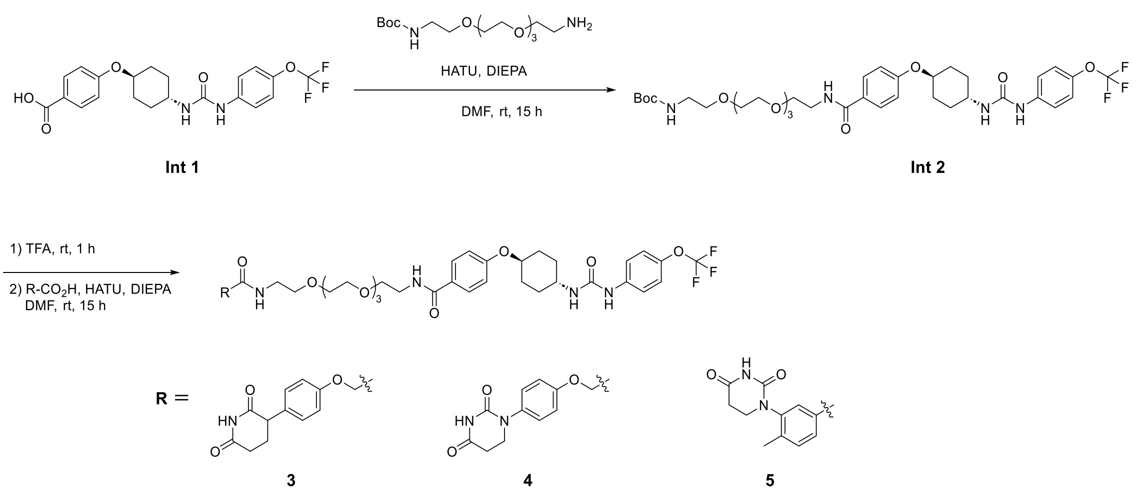


**Synthesis of Int 2**

**Int 1** (90.4 mg, 206 μmol, 1.0 equiv.) and HTAU (158.6 mg, 417 μmol, 2.0 equiv.) were added to a solution of *tert*-butyl (14-amino-3,6,9,12-tetraoxatetradecyl)carbamate (from AmBeed, 90. 2 mg, 268 μmol, 1.3 equiv.) in DMF (2.0 mL) and DIEPA (180 μL, 1.03 mmol, 5.0 equiv.). The reaction mixture was stirred at room temperature for 15 hours, then extracted with ethyl acetate three times. The combined organic layer was dried over $Na_2SO_4$, filtered, and concentrated *in vacuo*. The residue was further purified by flash chromatography (hexane: ethyl acetate).

**Int 2** (133 mg, 86 % yield). $^1$H NMR (300 MHz, $CD_3OD$) $\delta_H$ 7.79 (d, *J* = 8.9 Hz, 2H), 7.44 (d, *J* = 9.1 Hz, 2H), 7.15 (d, *J* = 8.4 Hz, 2H), 6.97 (d, *J* = 8.9 Hz, 2H), 4.46-4.34 (m, 1H), 3.70-3.53 (m, 16H), 3.48 (t, *J* = 5.6 Hz, 2H), 3.33-3.29 (m, 1H), 3.20 (t, *J* = 5.6 Hz, 2H), 2.21-2.00 (m, 4H), 1.68-1.33 (m, 13H). $^{13}$C NMR (75 MHz, $CD_3OD$) $\delta_C$ 169.82, 162.01, 157.30, 144.97, 140.23, 130.26, 127.56, 122.67, 120.85, 116.30, 80.05, 75.89, 71.53, 71.51, 71.31, 71.22, 71.05, 70.66, [72-70 ppm region several PEG peaks overlap], 41.27, 40.90, 31.42, 31.13, 28.77. $^{19}$F NMR (565 MHz, $CDCl_3$) $\delta_F$ -58.20. HRMS (ESI, positive): calcd. for $[C_{36}H_{51}N_4O_{10}F_3+Na]^+$ 779.3455, found 779.3459. White powder.

**Synthesis of compounds 3, 4, and 5**

**Int 2** (10.0 mg, 13.2 μmol, 1.0 equiv.) was dissolved in TFA (1.0 mL) and the reaction mixture was stirred at room temperature for 1 hour, then concentrated *in vacuo*. The residue and HATU (10.0 mg, 26.4 μmol, 2.0 equiv.) were added to a solution of carboxylate derivative (13.2 μmol, 1.0 equiv.) in DMF (500 mL) and DIEPA (20 μL, 115 μmol, 8.7 equiv.). The reaction mixture was stirred at room temperature for 15 hours, then filtered through a PTFE membrane and purified via preparative HPLC using a $H_2O$-MeCN gradient (99: 1 to 5: 95, v/v, 0.1% FA). A fraction containing the target molecule was lyophilized to give a solid.

Compound **3** (8.9 mg, 75 % yield). $^1$H NMR (600 MHz, $CD_3OD$) $\delta_H$ 7.80-7.76 (m, 2H), 7.45-7.41 (m, 2H), 7.21-7.18 (m, 2H), 7.17-7.14 (m, 2H), 6.99-6.95 (m, 4H), 4.51 (s, 2H), 4.44-4.36 (m, 1H), 3.83-3.78 (m, 1H), 3.68-3.52 (m, 18H), 3.48-3.42 (m, 2H), 2.73-2.65 (m, 1H), 2.64-2.58 (m, 1H), 2.25-2.11 (m, 4H), 2.10-2.03 (m, 2H), 1.63-1.54 (m, 2H), 1.48-1.36 (m, 2H). $^{13}$C NMR (151 MHz, $CD_3OD$) $\delta_C$ 176.52, 175.70, 171.14, 169.84, 162.04, 158.37, 157.34, 145.02, 140.24, 133.18, 130.81, 130.27, 127.58, 122.68, 120.89, 116.32, 116.06, 75.93, 71.55, 71.52, 71.30, 70.64, 70.43, 68.37, [72-67 ppm region several PEG peaks overlap], 49.57, 48.41, 40.91, 40.02, 32.20, 31.43, 31.15, 27.82. $^{19}$F NMR (565 MHz, $CD_3OD$) $\delta_F$ -59.83 HRMS (ESI, positive): calcd. for $[C_{44}H_{54}N_5O_{12}F_3+H]^+$ 902.3799, found 902.3812; calcd. for $[C_{44}H_{54}N_5O_{12}F_3+Na]^+$ 924.3619, found 924.3634. Purity (HPLC-UV 254 nm): >95%. ($^t$R= 2.539 min). White powder.

Compound **4** (11.3 mg, 95 % yield). $^1$H NMR (300 MHz, $CD_3OD$) $\delta_H$ 8.54 (br, 1H), 7.78 (d, $J$ = 8.8 Hz, 2H), 7.43 (d, $J$ = 9.1 Hz, 2H), 7.29 (d, $J$ = 9.0 Hz, 2H), 7.15 (d, $J$ = 8.6 Hz, 2H), 7.03 (d, $J$ = 9.0 Hz, 2H), 6.97 (d, $J$ = 8.9 Hz, 2H), 4.53 (s, 2H), 4.46-4.32 (m, 1H), 3.80 (t, $J$ = 7.1 Hz, 2H), 3.68-3.51 (m, 20H), 3.49-3.42 (m, 2H), 2.79 (t, $J$ = 6.9 Hz, 2H), 2.22-2.02 (m, 5H), 1.68-1.27 (m, 4H). $^{13}$C NMR (151 MHz, $CD_3OD$) $\delta_C$ 172.89, 170.89, 169.78, 162.00, 157.72, 157.32, 154.64, 144.95, 140.27, 136.90, 130.27, 128.34, 127.56, 122.65, 120.85, 116.38, 116.31, 75.91, 71.53, 71.50, 71.30, 71.28, 70.61, 70.39, 68.47, 55.81, 49.85, 46.78, 43.76, 40.89, 40.02, 32.12, 31.40, 31.14, 18.69, 17.26, 13.12. $^{19}$F NMR (565 MHz, $CD_3OD$) $\delta_F$ -60.71. HRMS (ESI, positive): calcd. for $[C_{43}H_{53}N_6O_{12}F_3+Na]^+$ 925.3571, found 925.3578. Purity (HPLC-UV 254 nm): >95%. ($^t$R= 2.615 min). White powder.

Compound **5** (9.3 mg, 79 % yield). $^1$H NMR (300 MHz, $CD_3OD$) $\delta_H$ 8.55 (br, 1H), 7.82-7.72 (m, 4H), 7.46-7.38 (m, 3H), 7.16 (d, $J$ = 8.4 Hz, 2H), 6.97 (d, $J$ = 8.9 Hz, 2H), 4.47-4.34 (m, 1H), 3.93-3.81 (m, 1H), 3.72-3.50 (m, 23H), 2.91-2.80 (m, 2H), 2.31 (s, 3H), 2.22-2.01 (m, 4H), 1.68-1.30 (m, 4H). $^{13}$C NMR (151 MHz, $CD_3OD$) $\delta_C$ 172.84, 169.82, 169.09, 162.03, 157.34, 154.19, 145.01, 141.77, 141.30, 140.25, 134.77, 132.29, 130.26, 127.99, 127.56, 127.44, 122.66, 120.87, 116.30, 75.91, 71.53, 71.51, 71.30, 70.62, 70.53, [72-70 ppm region several PEG peaks overlap], 55.83, 46.28, 43.78, 41.01, 40.89, 32.15, 31.41, 31.14, 18.69, 17.96, 17.26, 13.14. $^{19}$F NMR (565 MHz, $CD_3OD$) $\delta_F$ -60.70. HRMS (ESI, positive): calcd. for $[C_{43}H_{53}N_6O_{11}F_3+H]^+$ 887.3803, found 887.3825; calcd. for $[C_{43}H_{53}N_6O_{11}F_3+Na]^+$ 909.3622, found 909.3635. Purity (HPLC-UV 254 nm): >95%. ($^t$R= 2.576 min). White powder.

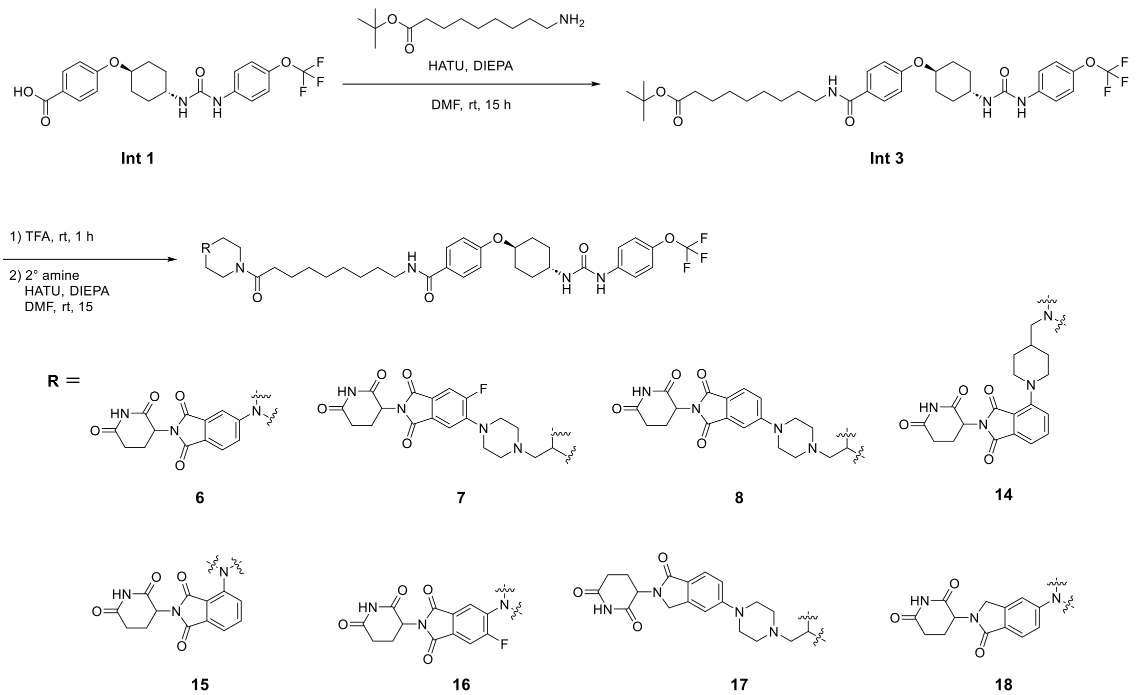


#### Synthesis of Int 3

**Int 1** (140.8 mg, 321 μmol, 1.0 equiv.) and HTAU (244.2 mg, 642 mmol, 2.0 equiv.) were added to a solution of *tert*-butyl (14-amino-3,6,9,12-tetraoxatetradecyl)carbamate (from AmBeed, 110.5 mg, 482 μmol, 1.5 equiv.) in DMF (2.0 mL) and DIEPA (300 μL, 1.72 mmol, 5.4 equiv.). The reaction mixture was stirred at room temperature for 15 hours, then extracted with ethyl acetate three times. The combined organic layer was dried over $Na_2SO_4$, filtered, and concentrated *in vacuo*. The residue was further purified by flash chromatography (hexane: ethyl acetate).

**Int 3** (159.4 mg, 76 % yield). $^1$H NMR (300 MHz, $CD_3OD$) $\delta_H$ 7.76 (d, $J$ = 8.9 Hz, 2H), 7.43 (d, $J$ = 9.1 Hz, 2H), 7.15 (d, $J$ = 8.4 Hz, 2H), 6.97 (d, $J$ = 8.9 Hz, 2H), 4.48-4.34 (m, 1H), 3.72-3.58 (m, 1H), 3.39-3.32 (m, 2H), 2.24-2.00 (m, 6H), 1.67-1.27 (m, 25H). $^{13}$C NMR (151 MHz, $CD_3Cl$) $\delta_C$ 173.50, 167.90, 160.76, 155.21, 144.09, 138.55, 128.89, 126.92, 121.89, 120.10, 115.49, 80.15, 75.65, 47.78, 40.46, 38.78, 35.68, 31.08, 30.32, 29.75, 29.31, 29.25, 29.09, 28.25, 27.10, 25.15. $^{19}$F NMR (282 MHz, $CD_3OD$) $\delta_F$ -59.93. HRMS (ESI, positive): calcd. for $[C_{34}H_{46}N_3O_6F_3+H]^+$ 650.3417, found 650.3410; calcd. for $[C_{34}H_{46}N_3O_6F_3+Na]^+$ 672.3236, found 672.3247. White powder.

**Synthesis of compounds 6 – 8, 14 – 18**

**Int 3** (10.0 mg, 15.4 µmol, 1.0 equiv.) was dissolved in TFA (1.0 mL) and the reaction mixture was stirred at room temperature for 1 hour, then concentrated *in vacuo*. The residue and HATU (11.7 mg, 30.8 µmol, 2.0 equiv.) were added to a solution of amine derivative (15.4 µmol, 1.0 equiv.) in DMF (1.0 mL) and DIEPA (30 µL, 172 µmol, 11.2 equiv.). The reaction mixture was stirred at room temperature for 15 hours, then filtered through a PTFE membrane and purified via preparative HPLC using a $H_2O$-MeCN gradient (99: 1 to 5: 95, v/v, 0.1% FA). A fraction containing the target molecule was lyophilized to give a solid.

Compound **6** (12.2 mg, 86 % yield). $^1$H NMR (600 MHz, $CD_3OD$) $\delta_H$ 7.78-7.74 (m, 2H), 7.42-7.69 (m, 1H), 7.45-7.21 (m, 2H), 7.38 (d, $J$ = 2.3 Hz, 1H), 7.24 (dd, $J$ = 8.6 Hz, 2.4 Hz 1H), 7.12 (d, $J$ = 8.3 Hz, 2H), 6.99-6.95 (m, 2H), 5.08 (dd, $J$ = 12.5 Hz, 5.5 Hz, 1H), 4.43-4.36 (m, 1H), 3.78-3.71 (m, 4H), 3.69-3.61 (m, 1H), 3.56-3.45 (m, 4H), 3.35 (t, $J$ = 7.1 Hz, 2H), 2.90-2.82 (m, 1H), 2.77-2.67 (m, 2H), 2.44 (t, $J$ = 7.5 Hz, 2H), 2.19-2.04 (m, 5H), 1.67-1.54 (m, 6H), 1.47-1.34 (m, 10H). $^{13}$C NMR (75 MHz, $CD_3OD$) $\delta_C$ 174.63, 174.44, 171.63, 169.79, 169.28, 168.91, 161.91, 157.33, 156.75, 145.06, 140.23, 135.60, 130.10, 127.81, 126.03, 122.66, 121.10, 120.87, 119.37, 116.30, 109.44, 75.90, 50.44, 46.13, 42.28, 40.92, 33.95, 32.18, 31.42, 31.14, 30.49, 30.31, 30.26, 30.21, 27.93, 26.40, 23.77. $^{19}$F NMR (565 MHz, $CD_3OD$) $\delta_F$ -60.69. HRMS (ESI, positive): calcd. for $[C_{47}H_{54}N_7O_9F_3+H]^+$ 918.4013, found 918.3995. Purity (HPLC-UV 254 nm): >95%. ($^t$R= 3.027 min). Yellow powder.

Compound **7** (13.1 mg, 82 % yield). $^1$H NMR (600 MHz, $CD_3OD$) $\delta_H$ 8.50 (br, 1H), 7.76 (d, $J$ = 8.9 Hz, 2H), 7.54 (d, $J$ = 11.3 Hz, 1H), 7.47 (d, $J$ = 7.3 Hz, 1H), 7.43 (d, $J$ = 9.1 Hz, 2H), 7.15 (d, $J$ = 8.4 Hz, 2H), 6.97 (d, $J$ = 8.9 Hz, 2H), 5.09 (d, $J$ = 12.8 Hz, 5.5 Hz, 1H), 4.54-4.49 (m, 1H), 4.44-4.36 (m, 1H), 4.01-3.92 (m, 1H), 3.70-3.61 (m, 1H), 3.35 (t, $J$ = 7.1 Hz, 2H), 3.13-3.06 (m, 1H), 2.90-2.82 (m, 1H), 2.77-2.67 (m, 2H), 2.67-2.59 (m, 5H), 2.42-2.35 (m, 2H), 2.29 (d, $J$ = 6.8 Hz, 2H), 2.19-2.04 (m, 5H), 1.93-1.80 (m, 3H), 1.65-1.54 (m, 6H), 1.48-1.28 (m, 11H), 1.18-1.01 (m, 2H). $^{13}$C NMR (151 MHz, $CD_3OD$) $\delta_C$ 174.57, 173.98, 171.47, 169.76, 168.36, 167.92, 161.92, 160.38, 157.34, 147.18, 145.01, 140.24, 130.51, 130.11, 127.82, 122.67, 120.87, 116.31, 114.55, 112.72, 112.55, 75.91, 65.31, 54.49, 50.95, 50.92, 50.69, 49.57, 47.18, 43.03, 40.95, 34.51, 34.18, 32.61, 32.17, 31.70, 31.43, 31.15, 30.53, 30.33, 30.29, 30.25, 27.98, 26.66, 23.70. $^{19}$F NMR (282 MHz, $CD_3OD$) $\delta_F$ -60.67 (s), -114.26 (dd, $J_{F\text{-}H}$ = 12.3 Hz, 9.0 Hz). HRMS (ESI, negative): calcd. for $[C_{53}H_{64}N_8O_9F_4\text{-}H]^-$ 1032.4654, found 1031.4637. Purity (HPLC-UV 254 nm): >95%. ($^t$R= 2.375 min). Yellow powder.

Compound **8** (14.9 mg, 95 % yield). $^1$H NMR (600 MHz, $CD_3OD$) $\delta_H$ 7.89-7.74 (m, 3H), 7.49 (d, $J$ = 2.2 Hz, 1H), 7.43 (d, $J$ = 9.1 Hz, 2H), 7.36 (dd, $J$ = 8.5 Hz, 2.3 Hz, 1H), 7.16 (d, $J$ = 8.4 Hz, 2H), 6.97 (d, $J$ = 8.9 Hz, 2H), 5.12-5.07 (dd, $J$ = 12.6 Hz, 5.5 Hz, 1H), 4.59 (d, $J$ = 13.4 Hz, 1H), 4.44-4.37 (m, 1H), 4.16 (br, 2H), 4.04 (d, $J$ = 13.7 Hz, 1H), 3.82-3.62 (m, 3H), 3.38-3.33 (m, 3H), 3.20-3.14 (m, 3H), 2.91-2.81 (m, 1H), 2.78-2.65 (m, 3H), 2.43-2.38 (m, 2H), 2.25-2.04 (m, 6H), 1.93-1.83 (m, 2H), 1.65-1.55 (m, 6H), 1.48-1.17 (m, 13H). $^{13}$C NMR (151 MHz, $CD_3OD$) $\delta_C$ 174.56, 174.08, 171.59, 169.81, 168.97,

168.65, 161.94, 157.35, 155.76, 140.23, 135.63, 130.11, 127.82, 126.13, 123.15, 122.68, 120.89, 120.65, 120.46, 116.33, 110.52, 75.91, 62.89, 53.07, 50.56, 49.57, 46.36, 45.96, 43.50, 42.22, 40.94, 34.07, 32.36, 32.17, 31.42, 31.14, 30.54, 30.36, 30.30, 30.24, 27.99, 26.58, 23.74. $^{19}$F NMR (565 MHz, $CD_3OD$) $\delta_F$ -60.53. HRMS (ESI, positive): calcd. for $[C_{53}H_{65}N_8O_9F_3+H]^+$ 1015.4905, found 1015.4872. Purity (HPLC-UV 254 nm): >95%. ($^t$R= 2.424 min). Yellow powder.

Compound **14** (11.3 mg, 72 % yield). $^1$H NMR (600 MHz, DMSO-$d_6$) $\delta_H$ 11.08(s, 1H), 8.62 (s, 1H), 8.28-8.22 (m, 1H), 7.78 (d, *J* = 8.8 Hz, 2H), 7.72-7.67 (m, 1H), 7.47 (d, *J* = 9.1 Hx, 2H), 7.38-7.36 (d, *J* = 7.1 Hz, 1H), 7.33 (d, *J* = 8.5 Hx, 1H), 7.21 (d, *J* = 8.5 Hz, 2H), 6.99 (d, *J* = 8.9 Hz, 2H), 6.32-6.26 (m, 1H), 5.09 (dd, *J* = 12.9 Hz, 5.5 Hz, 1H), 4.46-4.39 (m, 1H), 4.39-4.31 (m, 1H), 3.86-3.80 (m, 1H), 3.57-3.48 (m, 1H), 3.24-3.19 (m, 2H), 2.96 (t, *J* = 12.2 Hz, 1H), 2.91-2.83 (m, 1H), 2.63-2.56 (m, 1H), 2.55-2.51 (m, 4H), 2.29-2.23 (m, 2H), 2.19 (d, *J* = 7.1 Hz, 2H), 2.08-1.99 (m, 3H), 1.96-1.90 (m, 2H), 1.83-1.67 (m, 3H), 1.53-1.43 (m, 6H), 1.41-1.32 (m, 2H), 1.32-1.22 (m, 8H), 1.05-0.953 (m, 1H), 0.953-0.843 (m, 1H). $^{13}$C NMR (151 MHz, DMSO-$d_6$) $\delta_C$ 170.25, 167.05, 166.29, 166.00, 165.51, 159.54, 154.72, 149.71, 141.70, 140.31, 135.85, 133.67, 128.92, 126.71, 123.69, 121.48, 121.06, 119.37, 118.40, 116.51, 114.82, 74.15, 63.69, 53.04, 50.53, 48.80, 47.09, 44.99, 32.75, 32.41, 31.09, 30.97, 30.22, 29.92, 29.65, 29.18, 28.81, 28.75, 28.69, 27.59, 26.44, 25.96, 24.92, 22.06. $^{19}$F NMR (282 MHz, DMSO-$d_6$) $\delta_F$ -56.92. HRMS (ESI, positive): calcd. for $[C_{53}H_{65}N_8O_9F_3+H]^+$ 1014.4905, found 1015.4881. Purity (HPLC-UV 254 nm): >95%. ($^t$R= 2.503 min). Yellow powder.

Compound **15** (13.2 mg, 93 % yield). $^1$H NMR (600 MHz, $CD_3OD$) $\delta_H$ 7.76 (d, *J* = 8.8 Hz, 2H), 7.72-7.66 (m, 1H), 7.45-7.40 (m, 3H), 7.33 (d, *J* = 8.3 Hz, 2H), 7.16 (d, *J* = 8.5 Hz, 2H), 6.96 (d, *J* = 8.9 Hz, 2H), 5.11 (dd, *J* = 12.5 Hz, 5.5 Hz, 1H), 4.42-4.36 (m, 1H), 3.82-3.73 (m, 4H), 3.68-3.61 (m, 1H), 3.38-3.33 (m, 4H), 2.92-2.82 (m, 1H), 2.78-2.68 (m, 1H), 2.44 (t, *J* = 7.7 Hz, 2H), 2.18-2.11 (m, 3H), 2.09-2.04 (m, 2H), 1.67-1.53 (m, 6H), 1.47-1.31 (m, 10H). $^{13}$C NMR (75 MHz, $CD_3OD$) $\delta_C$ 174.61, 174.41, 171.61, 169.76, 168.88, 168.09, 161.91, 157.33, 151.22, 145.00, 140.24, 136.97, 135.47, 130.11, 127.81, 124.82, 122.67, 120.87, 119.30, 116.83, 116.30, 75.90, 52.56, 51.80, 50.49, 47.06, 42.87, 40.94, 34.03, 32.17, 31.43, 31.14, 30.49, 30.32, 30.28, 30.22, 27.94, 26.52, 23.69. $^{19}$F NMR (282 MHz, DMSO-$d_6$) $\delta_F$ -58.65. HRMS (ESI, positive): calcd. for $[C_{47}H_{54}N_7O_9F_3+H]^+$ 918.4013, found 918.4032. Purity (HPLC-UV 254 nm): >95%. ($^t$R= 3.087 min). Yellow powder.

Compound **16** (12.6 mg, 86 % yield). $^1$H NMR (600 MHz, $CD_3OD$) $\delta_H$ 8.55 (s, 1H), 7.76 (d, *J* = 8.9 Hz, 2H), 7.58 (d, *J* = 11.0 Hz, 1H), 7.51 (d, *J* = 7.3 Hz, 1H), 7.43 (d, *J* = 9.1 Hz, 2H), 7.16 (d, *J* = 8.5 Hz, 2H), 6.97 (d, *J* = 8.9 Hz, 2H), 5.12-5.07 (m, 1H), 4.43-4.36 (m, 1H), 3.79-3.70 (m, 4H), 3.68-3.61 (m, 1H), 3.37-3.34 (m, 3H), 3.28-3.24 (m, 2H), 2.92-2.81 (m, 1H), 2.79-2.67 (m, 2H), 2.44 (t, *J* = 7.5 Hz, 2H), 2.18-2.04 (m, 5H), 1.67-1.54 (m, 6H), 1.45-1.31 (m, 11H). $^{13}$C NMR (151 MHz, DMSO-$d_6$) $\delta_C$ 172.78, 170.86, 169.90, 166.62, 166.17, 165.57, 165.09, 159.57, 154.50, 141.94, 139.92, 128.95, 128.72, 126.75, 121.61, 121.10, 119.38, 118.54, 114.85, 114.00, 111.98, 74.13, 54.92, 49.10, 47.18, 44.67, 32.25, 30.95, 29.98, 29.67, 29.21, 28.85, 28.77, 28.73, 26.48, 24.76, 22.07. $^{19}$F NMR (282 MHz, $CD_3OD$) $\delta_F$ -60.69 (s), -114.35 (dd, $J_{F\text{-}H}$ = 12.6 Hz, 9.0 Hz). HRMS (ESI, positive): calcd. for $[C_{47}H_{53}N_7O_9F_4+H]^+$ 936.3920, found 936.3911; calcd. for $[C_{47}H_{53}N_7O_9F_4+Na]^+$ 958.3739, found 958.3727. Purity (HPLC-UV 254 nm): >95%. ($^t$R= 3.132 min). Yellow powder.

Compound **17** (11.4 mg, 74 % yield). $^1$H NMR (600 MHz, DMSO-$d_6$) $\delta_H$ 10.94 (s, 1H), 8.91 (s, 1H), 8.37 (s, 1H), 8.25 (t, *J* = 5.6 Hz, 1H), 7.78 (d, *J* = 8.8 Hz, 2H), 7.52-7.48 (m, 3H), 7.20 (d, *J* = 8.6 Hz, 2H), 7.06-7.04 (m, 2H), 6.99 (d, *J* = 8.9 Hz, 2H), 6.57 (d, *J* = 7.5 Hz, 1H), 5.04 (dd, *J* = 13.3 Hz, 5.1 Hz, 1H), 4.44-4.19 (m, 4H), 3.82 (d, *J* = 13.1 Hz, 1H), 3.58-3.50 (m, 3H), 3.24-3.18 (m, 4H), 2.99-2.87 (m, 3H), 2.62-2.57 (m, 1H), 2.40-2.32 (m, 1H), 2.27-2.23 (m, 2H), 2.17 (d, *J* = 7.1 Hz, 2H),

2.09-2.00 (m, 2H), 1.99-1.91 (m, 3H), 1.80-1.69 (m, 3H), 1.51-1.45 (m, 6H), 1.40-1.33 (m, 2H), 1.32-1.22 (m, 9H), 1.03-0.867 (m, 2H). $^{13}$C NMR (151 MHz, $CD_3OD$) $\delta_C$ 174.73, 173.98, 172.44, 171.97, 169.75, 161.93, 157.33, 156.00, 145.87, 145.01, 140.25, 130.12, 127.83, 125.31, 122.74, 122.68, 120.87, 116.42, 116.31, 109.50, 75.91, 65.23, 54.48, 53.51, 49.57, 47.13, 42.98, 40.96, 34.40, 34.18, 32.57, 32.41, 31.67, 31.43, 31.15, 30.53, 30.33, 30.29, 30.25, 27.98, 26.66, 24.20. $^{19}$F NMR (565 MHz, $CD_3OD$) $\delta_F$ -60.69. HRMS (ESI, positive): calcd. for $[C_{53}H_{67}N_8O_8F_3+H]^+$ 1001.5112, found 1001.5086. Purity (HPLC-UV 254 nm): >95%. ($^t$R= 2.399 min). White powder.

Compound **18** (13.0 mg, 93 % yield). $^1$H NMR (600 MHz, DMSO-$d_6$) $\delta_H$ 10.94 (s, 1H), 8.98 (s, 1H), 8.50 (s, 1H), 8.25 (t, *J* = 5.6 Hz, 1H), 7.78 (d, *J* = 8.8 Hz, 2H), 7.54 (d, *J* = 8.4 Hz, 1H), 7.50 (d, *J* = 7.9 Hz, 2H), 7.20 (d, *J* = 8.6 Hz, 2H), 7.11- 7.05 (m, 2H), 6.99 (d, *J* = 8.9 Hz, 2H), 6.88 (br, 3H), 6.64 (d, *J* = 7.6 Hz, 1H), 6.28 (s, 1H), 5.05 (dd, *J* = 13.3 Hz, 5.1 Hz, 1H), 4.46-4.38 (m, 1H), 4.36-4.31 (m, 2H), 3.65-3.57 (m, 4H), 3.56-3.49 (m, 1H), 3.28-3.18 (m, 3H), 2.95-2.85 (m, 1H), 2.62-2.56 (m, 1H), 2.42-2.30 (m, 4H), 2.09-2.01(m, 2H), 1.99-1.89 (m, 3H), 1.55-1.43 (m, 6H), 1.41-1.22 (m, 11H). $^{13}$C NMR (151 MHz, $CD_3OD$) $\delta_C$ 174.72, 174.35, 172.42, 171.88, 169.77, 161.92, 157.34, 155.75, 145.89, 145.02, 140.25, 130.11, 127.82, 125.38, 123.16, 122.67, 120.88, 116.75, 116.30, 109.86, 75.91, 53.53, 49.60, 49.57, 46.52, 42.53, 40.92, 33.97, 32.40, 31.42, 31.14, 30.50, 30.31, 30.26, 30.21, 27.94, 26.48, 24.18. $^{19}$F NMR (565 MHz, $CD_3OD$) $\delta_F$ -59.83. HRMS (ESI, positive): calcd. for $[C_{47}H_{56}N_7O_8F_3+H]^+$ 904.4221, found 904.4211. Purity (HPLC-UV 254 nm): >95%. ($^t$R= 2.873 min). White powder.

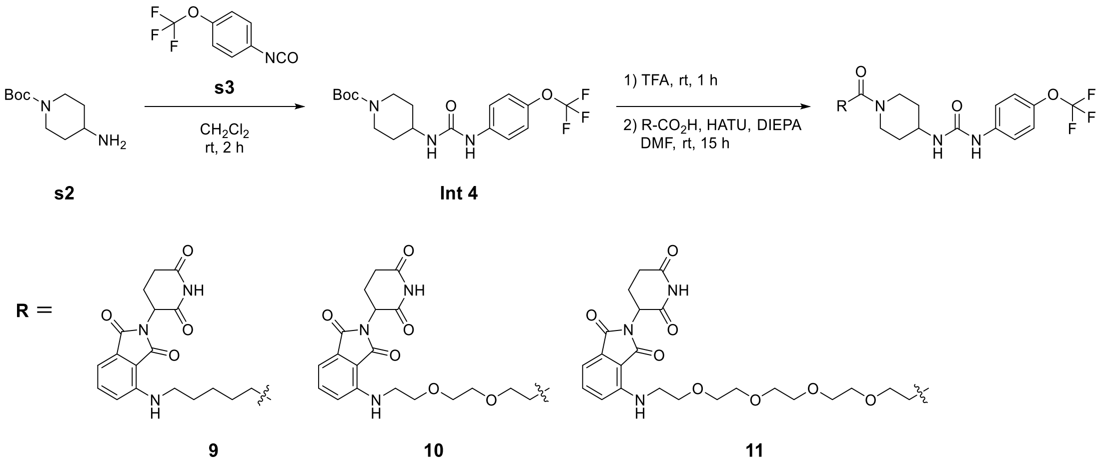


**Synthesis of Int 4**

**Int 4** was prepared by the similar procedure as that described in literature.[7] 4-(Trifluoromethoxy)phenyl isocyanate (150 µL, 994 µmol, 1.0 equiv.) and 4-amino-1-Boc-piperidine (239.5 mg, 1.20 mmol, 1.2 equiv.) was dissolved in $CH_2Cl_2$ (10 mL) and stirred for 2 hours at room temperature. The reaction was quenched by addition of water (5 mL) and then the solution was extracted with ethyl acetate three times. The combined organic layer was dried over $Na_2SO_4$, filtered, and concentrated *in vacuo*. The residue was further purified by flash chromatography (hexane: ethyl acetate).

**Int 4** (250. 5 mg, 63 % yield). $^1$H NMR (600 MHz, $CDCl_3$) $\delta_H$ 7.78 (s, 1H), 7.29 (d, *J* = 8.9 Hz, 2H), 7.07 (d, *J* = 8.6 Hz, 2H), 5.62 (d, *J* = 7.7 Hz, 1H), 3.91 (br, 2H), 3.81-3.71 (m, 1H), 2.82 (t, *J* = 11.8 Hz, 2H), 1.86 (d, *J* = 10.9 Hz, 1H), 1.44 (s, 9H), 1.22-1.08 (m, 2H). $^{13}$C NMR (151 MHz, $CDCl_3$) $\delta_C$ 155.38, 155.14, 144.28, 138.02, 121.93, 120.19, 80.38, 46.96, 46.85, 32.64, 29.78, 28.50. $^{19}$F NMR (565 MHz, $CD_3Cl$) $\delta_F$ -58.15. White powder.

**Synthesis of compounds 9, 10, and 11**

**Int 4** (10.0 mg, 24.8 µmol, 1.0 equiv.) was dissolved in TFA (1.0 mL) and the reaction mixture was stirred at room temperature for 1 hour, then concentrated *in vacuo*. The residue and HATU (18.9 mg, 49.6 μmol, 2.0 equiv.) were added to a solution of carboxylate derivative (24.8 μmol, 1.0 equiv.) in DMF (500 μL) and DIEPA (40 μL mg, 230 μmol, 9.3 equiv.). The reaction mixture was stirred at room temperature for 15 hours, then filtered through a PTFE membrane and purified via preparative HPLC using a $H_2O$-MeCN gradient (99: 1 to 5: 95, v/v, 0.1% FA). A fraction containing the target molecule was lyophilized to give a solid.

Compound **9** (15.9 mg, 95 % yield). $^1$H NMR (600 MHz, $CD_3OD$) $\delta_H$ 7.55 (dd *J* = 8.6 Hz, 7.1 Hz, 1H), 7.43 (d, *J* = 9.1 Hz, 2H), 7.15 (d, *J* = 8.4 Hz, 2H), 7.07-7.03 (m, 2H), 5.05 (dd, *J* = 12.2 Hz, 5.7 Hz, 1H), 4.40-4.32 (m, 1H), 3.95-3.88 (m, 1H), 3.85-3.78 (m, 1H), 3.35 (t, *J* = 6.8 Hz, 2H), 3.27-3.20 (m, 1H), 2.92-2.81 (m, 2H), 2.77-2.67 (m, 2H), 2.48-2.38 (m, 2H), 2.14-2.08 (m, 1H), 2.04-1.98 (m, 1H), 1.97-1.91 (m, 1H), 1.74-1.64 (m, 4H), 1.52-1.25 (m, 5H). $^{13}$C NMR (151 MHz, $CD_3OD$) $\delta_C$ 174.65, 173.86, 171.69, 171.68, 170.81, 169.32, 157.18, 148.29, 145.10, 140.13, 137.26, 133.94, 122.68, 120.97, 118.01, 111.02, 50.19, 48.14, 45.75, 43.20, 43.19, 41.78, 34.08, 33.89, 33.11, 32.20, 30.04, 27.62, 26.32, 23.81. $^{19}$F NMR (565 MHz, $CD_3OD$) $\delta_F$ -60.58. HRMS (ESI, positive): calcd. for $[C_{32}H_{35}N_6O_7F_3+H]^+$ 673.2598, found 673.2606; calcd. for $[C_{32}H_{35}N_6O_7F_3+Na]^+$ 695.2417, found 695.2425. Purity (HPLC-UV 254 nm): >95%. ($^t$R= 2.762 min). Yellow powder.

Compound **10** (16.1 mg, 90 % yield). $^1$H NMR (300 MHz, $CD_3OD$) $\delta_H$ 7.58-7.50 (m, 1H), 7.42 (d, *J* = 9.0 Hz, 2H), 7.15 (d, *J* = 8.5 Hz, 2H), 7.09 (d, *J* = 8.5 Hz, 1H), 7.04 (d, *J* = 7.1 Hz, 1H), 5.05 (dd, *J* = 12.2 Hz, 5.0 Hz, 1H), 4.40-4.29 (m, 1H), 4.00-3.88 (m, 1H), 3.87-3.59 (m, 9H), 3.50 (t, *J* = 5.1 Hz, 2H), 3.28-3.15 (m, 1H), 2.94-2.58 (m, 6H), 2.17-2.06 (m, 1H), 2.01-1.86 (m, 2H), 1.50-1.27 (m, 2H). $^{13}$C NMR (75 MHz, $CD_3OD$) $\delta_C$ 174.66, 172.10, 171.65, 170.71, 169.29, 157.14, 148.23, 145.08, 140.13, 137.22, 133.89, 122.67, 120.93, 118.28, 112.04, 111.30, 71.57, 70.63, 68.69, 50.21, 48.10, 45.95, 43.26, 41.80, 34.50, 33.96, 33.04, 32.20, 23.82. $^{19}$F NMR (565 MHz, $CD_3OD$) $\delta_F$ -60.68. HRMS (ESI, positive): calcd. for $[C_{33}H_{37}N_6O_9F_3+H]^+$ 719.2652, found 719.2660; calcd. for $[C_{33}H_{37}N_6O_9F_3+Na]^+$ 741.2472, found 741.2482. Purity (HPLC-UV 254 nm): >95%. ($^t$R= 2.590 min). Yellow powder.

Compound **11** (10.5 mg, 53 % yield). $^1$H NMR (300 MHz, $CD_3OD$) $\delta_H$ 7.57-7.50 (m, 1H), 7.42 (d, *J* = 9.0 Hz, 2H), 7.14 (d, *J* = 8.6 Hz, 2H), 7.10-7.01 (m, 2H), 7.04 (d, *J* = 7.1 Hz, 1H), 5.10-5.01 (m, 1H), 4.43-4.32 (m, 1H), 4.03-3.92 (m, 1H), 3.86-3.54 (m, 17H), 3.49 (t, *J* = 5.2 Hz, 2H), 3.28-3.16 (m, 1H), 2.94-2.54 (m, 6H), 2.17-2.06 (m, 1H), 2.05-1.87 (m, 2H), 1.55-1.27 (m, 2H). $^{13}$C NMR (75 MHz, $CD_3OD$) $\delta_C$ 174.66, 172.06, 171.60, 170.71, 169.34, 164.35, 157.12, 148.20, 140.12, 137.20, 133.88, 122.66, 120.91, 118.25, 112.02, 111.33, 71.68, 71.62, 71.58, 71.53, 71.48, 70.62, 68.60, 50.19, 46.00, 43.23, 41.87, 34.42, 33.95, 33.08, 32.20, 23.82. $^{19}$F NMR (565 MHz, $CD_3OD$) $\delta_F$ -60.51. HRMS (ESI, positive): calcd. for $[C_{37}H_{45}N_6O_{11}F_3+H]^+$ 807.3177, found 807.3187; calcd. for $[C_{37}H_{45}N_6O_{11}F_3+Na]^+$ 829.2996, found 829.3004. Purity (HPLC-UV 254 nm): >95%. ($^t$R= 2.592 min). Yellow powder.

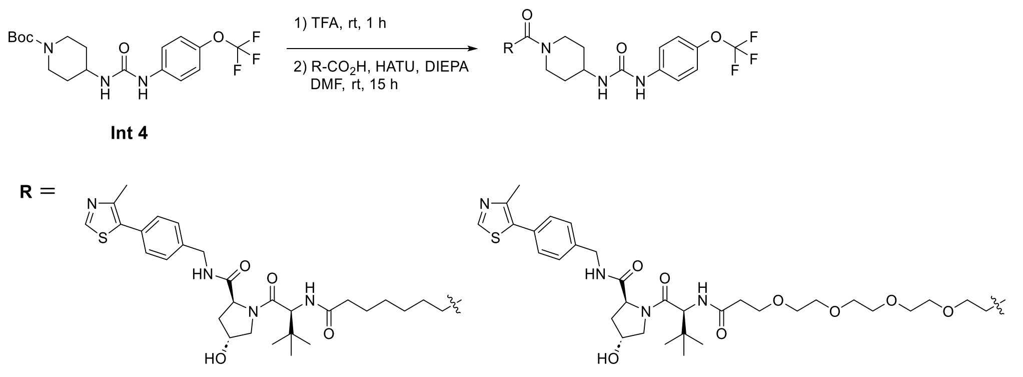


**12** **13**

**Synthesis of compounds 12 and 13**

**Int 4** (10.0 mg, 24.8 μmol, 1.0 equiv.) was dissolved in TFA (1.0 mL) and the reaction mixture was stirred at room temperature for 1 hour, then concentrated *in vacuo*. The residue and HATU (18.9 mg, 49.6 μmol, 2.0 equiv.) were added to a solution of carboxylate derivative (24.8 μmol, 1.0 equiv.) in DMF (500 μL) and DIEPA (40 μL mg, 230 μmol, 9.3 equiv.). The reaction mixture was stirred at room temperature for 15 hours, then filtered through a PTFE membrane and purified via preparative HPLC using a $H_2O$-MeCN gradient (99: 1 to 5: 95, v/v, 0.1% FA). A fraction containing the target molecule was lyophilized to give a solid.

Compound **12** (14.4 mg, 85 % yield). $^1$H NMR (300 MHz, $CD_3OD$) $\delta_H$ 8.98 (s, 1H), 7.52-7.40 (m, 6H), 7.17 (d, $J$ = 8.5 Hz, 2H), 4.69-4.33 (m, 6H), 4.00-3.77 (m, 4H), 3.31-3.18 (m, 1H), 2.97-2.82 (m, 1H), 2.50 (s, 3H), 2.42 (t, $J$ = 7.6 Hz, 2H), 2.37-2.18 (m, 3H), 2.16-1.90 (m, 3H), 1.72-1.55 (m, 4H), 1.51-1.28 (m, 6H), 1.06 (2, 9H). $^{13}$C NMR (75 MHz, $CD_3OD$) $\delta_C$ 175.99, 174.50, 174.02, 172.34, 157.17, 153.12, 148.52, 145.09, 140.45, 140.13, 133.73, 131.24, 130.37, 129.00, 122.67, 120.94, 71.06, 60.82, 58.97, 58.03, 45.77, 43.69, 41.77, 38.94, 36.53, 36.46, 34.11, 34.01, 33.13, 30.02, 29.92, 27.03, 26.81, 26.49, 15.58. $^{19}$F NMR (565 MHz, $CD_3OD$) $\delta_F$ -60.14. HRMS (ESI, positive): calcd. for $[C_{43}H_{56}N_7O_7F_3S+H]^+$ 872.3992, found 872.4007; calcd. for $[C_{43}H_{56}N_7O_7F_3S+Na]^+$ 894.3812, found 894.3825. Purity (HPLC-UV 254 nm): >95%. ($^t$R= 2.660 min). White powder.

Compound **13** (12.6 mg, 81 % yield). $^1$H NMR (300 MHz, $CD_3OD$) $\delta_H$ 8.99 (s, 1H), 7.51-7.38 (m, 6H), 7.15 (d, $J$ = 8.6 Hz, 2H), 4.69-4.46 (m, 4H), 4.44-4.32 (m, 2H), 4.03-3.54 (m, 20H), 3.30-3.17 (m, 1H), 2.97-2.83 (m, 1H), 2.77-2.42 (m, 7H), 2.29-1.88 (m, 4H), 1.56-1.24 (m, 2H), 1.04 (s, 9H). $^{13}$C NMR (75 MHz, $CD_3OD$) $\delta_C$ 174.51, 173.72, 172.12, 171.98, 157.16, 153.25, 148.34, 145.10, 140.54, 140.14, 133.85, 131.13, 129.03, 122.69, 120.94, 71.61, 71.54, 71.48, 71.40, 71.07, [72-71 ppm region several PEG peaks overlap], 68.58, 68.29, 60.82, 58.94, 58.01, 45.96, 43.69, 41.86, 38.95, 37.33, 36.78, 34.41, 33.98, 33.09, 31.54, 29.73, 27.04, 15.52. $^{19}$F NMR (565 MHz, $CD_3OD$) $\delta_F$ -60.31. HRMS (ESI, positive): calcd. for $[C_{47}H_{64}N_7O_{11}F_3S+H]^+$ 992.4415, found 992.4424. Purity (HPLC-UV 254 nm): >95%. ($^t$R= 2.486 min). White powder.

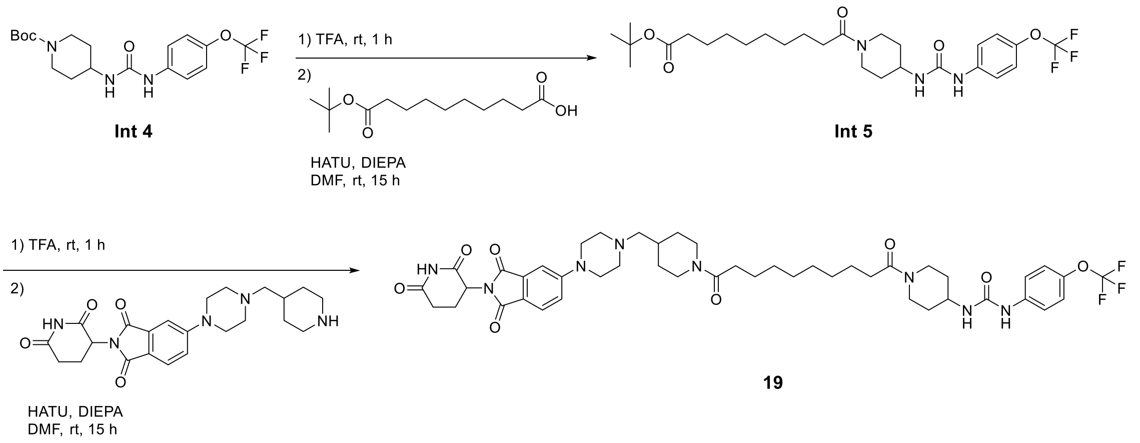


**Synthesis of Int 5**

**Int 4** (115.7 mg, 286.8 µmol, 1.0 equiv.) was dissolved in TFA (200 µL) and $CH_2Cl_2$ (2 mL), and the reaction mixture was stirred at room temperature for 1 hour, then concentrated *in vacuo*. The residue and HATU (172.26 mg, 453.0 µmol, 1.6 equiv.) were added to a solution of 10-(tert-butoxy)-10-oxodecanoic acid (from AmBeed, 125.3 mg, 484.8 µmol, 1.7 equiv.) in DMF (3.0 mL) and DIEPA (200 µL, 1.15 mmol, 4.0 equiv.). The reaction mixture was stirred at room temperature for 15 hours, then extracted with ethyl acetate three times. The combined organic layer was dried over $Na_2SO_4$, filtered, and concentrated *in vacuo*. The residue was further purified by flash chromatography (hexane: ethyl acetate).

**Int 5** (113.1 mg, 73 % yield). $^1$H NMR (300 MHz, $CDCl_3$) $\delta_H$ 7.57 (br, 1H), 7.39 (d, *J* = 9.0 Hz, 2H), 7.11 (d, *J* = 8.6 Hz, 2H), 5.42 (d, *J* = 8.0 Hz, 1H), 4.58-4.45 (m, 1H), 4.02-3.76 (m, 2H), 3.14 (t, *J* = 12.0 Hz, 1H), 2.75 (t, *J* = 11.2 Hz, 1H), 2.34 (t, *J* = 7.7 Hz, 2H), 2.23-2.09 (m, 3H), 2.00-1.87 (m, 1H), 1.67-1.47 (m, 4H), 1.44 (s, 9H), 1.37-1.11 (m, 10H). $^{13}$C NMR (75 MHz, $CD_3Cl$) $\delta_C$ 173.83, 172.49, 154.93, 144.15, 138.32, 121.95, 119.92, 80.37, 46.72, 45.01, 41.20, 35.80, 34.17, 33.46, 32.68, 29.31, 29.19, 29.02, 28.26, 25.62, 25.19. $^{19}$F NMR (282 MHz, $CD_3Cl$) $\delta_F$ -58.11. HRMS (ESI, positive): calcd. for $[C_{27}H_{40}N_3O_5F_3+Na]^+$ 566.2818, found 566.2828. White powder.

**Synthesis of compound 19**

**Int 5** (9.6 mg, 17.6 µmol, 1.0 equiv.) was dissolved in TFA (800 µL) and the reaction mixture was stirred at room temperature for 1 hour, then concentrated *in vacuo*. The residue and HATU (13.4 mg, 35.2 mmol, 2.0 equiv.) were added to a solution of pomalidomide 5'-piperazine-4-methylpiperidine (from Tocris, 7.7 mg, 17.6 µmol, 1.0 equiv.) in DMF (500 µL) and DIEPA (20 µL, 110 µmol, 6.5 equiv.). The reaction mixture was stirred at room temperature for 15 hours, then filtered through a PTFE membrane and purified via preparative HPLC using a $H_2O$-MeCN gradient (99:1 to 5:95, v/v, 0.1% FA). A fraction containing the target molecule was lyophilized to give a solid.

Compound **19** (8.7 mg, 55 % yield). $^1$H NMR (300 MHz, $CD_3OD$) $\delta_H$ 7.69 (d, *J* = 8.5 Hz, 1H), 7.48-7.36 (m, 3H), 7.25 (dd, *J* = 8.6 Hz, 2.2 Hz, 1H), 7.16 (d, *J* = 8.5 Hz, 1H), 5.07 (dd, *J* = 12.4 Hz, 5.7 Hz, 1H), 4.60-4.48 (m, 1H), 4.46-4.32 (m, 1H), 4.04-3.74 (m, 3H), 3.58-3.47 (m, 4H), 3.27-3.05 (m, 2H), 2.95-2.59 (m, 9H), 2.49-2.33 (m, 6H), 2.17-1.78 (m, 6H), 1.68-1.52 (m, 4H), 1.50-1.27 (m, 10H), 1.26-1.00 (m, 2H). $^{13}$C NMR (75 MHz, DMSO-$d_6$) $\delta_C$ 173.26, 170.85, 170.73, 170.54, 168.02, 167.44, 163.83, 155.70, 154.80, 140.21, 134.31, 125.34, 122.06, 119.11, 118.77, 118.21, 108.33, 64.05, 53.16, 49.23, 47.37, 46.73, 45.45, 44.13, 41.44, 33.20, 33.10, 32.91, 32.77, 32.28, 31.54, 31.44, 30.68, 29.27, 25.41, 25.37, 22.64. $^{19}$F NMR (282 MHz, $CD_3OD$) $\delta_F$ -60.69. HRMS (ESI, positive): calcd. for $[C_{46}H_{59}F_3N_8O_8+H]^+$ 909.4486, found 909.4498. Purity (HPLC-UV 254 nm): >95%. ($^t$R= 2.204 min). Yellow powder.

**Biology experiments**

***Materials***

Precision Plus Protein Dual Color Standard as protein ladder, SDS-PAGE (4–12%) gel, and Clarity western ECL substrate for immunoblotting were purchased from Bio-Rad. Trifluoroacetic acid (TFA), 0.25% Trypsin-EDTA, RIPA buffer, protease inhibitor, anti-GAPDH antibody from mouse, BCA protein assay kit-reducing agent compatible, sulfic acid, phenylmethylsulfonyl fluoride (PMSF), microplate for ELISA, and 3,3′,5,5′-tetramethylbenzidine (TMB) substrate for ELISA were purchased from ThermoFisher Scientific. Bullet Blocking One for Western Blotting as blocking reagent was purchased from Nacalai USA Inc. Dimethyl sulfoxide (DMSO), MG132, bovine serum albumin (BSA), Ammonium bicarbonate ($NH_4HCO_3$), dithiothreitol, 2-iodoacetamide, and hydroxypropyl-β-cyclodextrin were purchased from Sigma Aldrich. Anti-sEH antibody-HRP conjugate and anti-β-actin-HRP conjugate were purchased from Santa Cruz Biotechnology. Anti-catalase monoclonal antibody from rabbit was purchased from Cell Signaling Technology. Streptavidin-poly (Horseradish peroxidase) was purchased from Fitzgerald. Trypsin was purchased from Promega. Methanol ($CH_3OH$), water, and MeCN were purchased from Fisher. and S-trap was purchased from Protifi. Oasis HLB resin (30 μm) was purchased from Waters. 96-well Orochem filter plate was purchased from Orochem. PEG400 was purchased from MedChemExpress. Dulbecco's phosphate buffered saline (DPBS) was purchased from Cytiva. Fetal bovine serum (FBS) was purchased from Phenix. Scientific. High-glucose (4.5 g/L) Dulbecco’s modified Eagle’s medium (DMEM) was purchased from Corning. DMEM/F12 was purchased from Gibco. The media was supplemented with 10% FBS (Phenix Scientific), 100 units/mL of penicillin, and 100 mg/mL of streptomycin (Cytiva). Saline was purchased from Pfizer. Anti-IRE1 monoclonal antibody from rabbit was purchased from Cell Signaling Technology. Anti-phospho-IRE1$^{S724}$ polyclonal antibody from rabbit was purchased from Abcam. Anti-eIF2α, anti-phospho- eIF2α$^{S51}$, anti-PERK, and anti-phospho-PERK$^{T980}$ were purchased from Santa Cruz Biotechnology.

***Cells and Animals***

Human hepatocyte carcinoma cells (HepG2) and kidney cells (HEK293T) were purchased from American Type Culture Collection (ATCC). Human breast adenocarcinoma cells (MDA-MB-231) were provided from Dr, R. J. Eddy (Department of Pathology, Albert Einstein College of Medicine). Human bone marrow neuroblast cells (SH-SY5Y) were provided from Dr. Q. Zhang. The cell lines (HepG2, HEK293T, and MDA-MB-231) were cultured in medium (DMEM). SH-SY5Y cells were cultured in DMEM/F12. The cells were cultured at 37 °C in a 5% $CO_2$ incubator. Cells were used in biological assays at 3 - 20 passages after thawing. 5-week-old male C57BL/6J mice (Strain #:**000664**) were purchased from the Jackson Laboratory.

***Compound treatment and immunoblotting analysis***

Cells were treated with sEH PROTAC or DMSO vehicle in complete medium. After 2–48 hours of incubation, the medium was decanted, and the cells were washed with cold DPBS (Cytiva) and lysed using RIPA buffer with protease inhibitor cocktail (ThermoFisher Scientific). The cell lysates were then resolved using SDS-PAGE and transferred onto a nitrocellulose membrane using the iBlot2 Gel Transfer System (ThermoFisher Scientific). The membranes were blocked in Bullet Blocking One for Western Blotting (Nacalai USA Inc.) for 5 minutes at room temperature and probed with the primary antibody in Can Get Signal Solution 1 (TOYOBO) overnight. The protein ladder used in this experiment was Precision Plus Protein Dual Color Standard (Bio-Rad). The membranes were then probed with goat anti-rabbit and goat anti-mouse secondary antibodies (Cell Signaling) in Can Get Signal Solution 2 (TOYOBO), and blots were incubated with the Clarity Western ECL Substrate (Bio-Rad) and imaged using the iBright (ThermoFisher Scientific). ImageJ was used to quantify the amount of protein by immunoblotting. Each experiment has three independent repeats.

***Evaluation of cell viability***

HepG2 cells were seeded in 96-well plate and incubated for 24 hours. After removing the cell culture medium, the cells were treated with DMSO or each compound (final concentration 1 μM) and incubated for 24 hours. Then, the cell death was measured using MTT assay. After removing the medium, 100 μL of 0.2–0.5 mg/mL MTT dye (3-(4,5-dimethylthiazol-2-yl)-2,5-diphenyltetrazolium bromide) solution in DMEM was added to the well and then incubated at 37°C for 2–3 hours. After the medium containing MTT, 200 μL of DMSO was added to the well to dissolve the product and the absorbance of 590 nm was measured.

***Evaluation of chemical stability***

The 10 mM stock solution of compound in DMSO was diluted into the cell culture medium (DMEM) at a final compound concentration 100 μM. The solution was incubated at 37 °C or 4 °C for 24 hours. After incubation, the solution was analyzed with LC-MS using a $H_2O$-MeCN gradient (100:0 to 5:95, v/v, 0.1% FA).

***Evaluation of microsomal stability***

Metabolic stability of compounds **1a**, **6**, and **8** were evaluated at ChemPartner (China). Ketanserin was used as a reference compound. Pooled mouse liver microsomes were purchased from a commercial supplier (BD Gentest). The degradation kinetics of each 0.5 μM compound in 0.25 mg/mL of mouse liver microsomes were incubated in 0.1 M potassium phosphate buffer (pH 7.4), 1.0 mM ethylenediaminetetraacetic acid (EDTA), and 1 mM nicotinamide adenine dinucleotide phosphate (NADPH) at 37 °C. At 5, 10, 15, 30, and 45 min, 135 μL of MeCN containing internal standard (osalmid or imipramine) were added to the wells of corresponding plates, respectively, to stop the reaction. After quenching the reaction, the solutions in the plate were shaken for 10 minutes (600 rpm/min) by the vibrator (IKA, MTS 2/4) and then centrifuged at 5,594 g for 15 minutes (Thermo Multifuge × 3R) to removing the precipitates. 50 μL of the supernatant in each well were transferred into a 96-well sample plate containing 50 μL of ultra-pure water (Millipore, ZMQS50F01) for LC/MS (UPLC-MS/MS-048 (Triple Quad 6500+), UPLC-MSMS-32 (Triple Quad 6500+)) analysis. The experiment was performed in duplicate.

***Pharmacokinetic studies***

Pharmacokinetic analysis of compounds **8**, **17**, and **19** was performed at ChemPartner (China). Eanch compound concentration in mouse plasma were measured by quantitative LC-MS/MS (UPLC/MS-MS-51 (Triple Quad 7500)) using each standard curve. Calibration and quality control samples were prepared using blank plasma from untreated mice. Compounds **8**, **17**, and **19** were dissolved in mixed solvents (10% DMSO, 40% PEG400, 50% (50% hydroxypropyl-β-cyclodextrin) for intraperitoneal injection (i.p.), with the final concentration of 2.0 mg/mL. For calculating the administration dose, all the CD1 mice (30–31 g, 6–8 weeks, male, $n$ = 3) were weighed before injection. A compound solution was i.p. injected into mice at a dose of 10 mg/kg. At 0.25, 0.5, 1, 2, 4, 6, 8, and 24 hours, approximately 40 μL whole blood was collected in $K_2$EDTA tube via facial vein. Blood samples were put on ice and centrifuged at 2,000 g for 5 minutes to obtain plasma sample within 15 minutes. An aliquot of the 10 μL plasma sample was added to 100 μL of internal standard (Propranolol, 50 ng/mL) in MeCN and the mixture was shaken for 5 minutes and centrifuged at 14,000 rpm for 5 minutes at 4 °C. 50 μL of supernatant was transferred to a new plate and sealed, and then 3 μL of solution was injected to LC-MS/MS. PK parameters were estimated by non-compartmental model using WinNonlin 8.2.

***In vivo sEH degradation***

All animal experiments were performed following protocols approved by Albert Einstein College of Medicine Institutional Animal Care and Use Committee (Protocol #00001439). Compounds **8** was dissolved in mixed solvents (10% DMSO, 40% PEG400, 50% (50% hydroxypropyl-β-cyclodextrin) with the final concentration of 3.0 mg/mL. All the C57BL/6J mice (22–28 g, 5 weeks, male, $n$ = 3) were weighed before injection. A compound solution was injected intraperitoneally (i.p.) at a dose of 12 mg/kg. After 24 hours, the mice ($n$ = 3 per group) were sacrificed by carbon dioxide asphyxia. The liver and brown adipose tissue (BAT) were rapidly removed and homogenized in 1xRIPA buffer with protease inhibitor cocktail and EDTA (final concentration 5 mM) by using TissueLyser LT (QIAGEN) for 2–5 minutes. The homogenate was centrifuged at 15,000 g for 5 minutes at 4 ℃ and the lower layer was collected. Centrifugation and recovery of the lower layer were repeated two more times.

***Measurement of Biochemical Inhibitory Potency of PROTAC Compounds against Human and Mouse sEH Hydrolase***

The inhibitory potency of compounds was measured using a fluorescent assay as previously described.[8] The $IC_{50}$ values are defined as inhibitor concentrations that reduced the enzymatic activity by 50% and were calculated in the linear range of the dose–response curve for log inhibitor concentration vs % inhibition. These values are the averages of three replicates, obtained from the experiments with various concentrations of the inhibitors (1–10,000 nM) including at least two datum points above and two datum points below the $IC_{50}$. High Z′-factor values (0.75–0.80) were obtained for the assay. The standard error for the $IC_{50}$ values is between 10% and 20% for the assays performed here, suggesting that the differences of 2-fold or greater would be significant.

***Quantitation of sEH Protein Level Using a PolyHRP ELISA***

The sEH protein level of cells treated with compound was measured using an ELISA method.[9,10] The high-binding microplate (Thermo Fisher Scientific) was coated with an anti-human sEH rabbit serum (1:2000 dilution) in 50 mM carbonate–bicarbonate buffer (pH 9.6) overnight at 4 °C. Then, the plate was blocked with 3% (w/v) of skim milk in PBS for 1 hour at room temperature. Human sEH standards and samples with different dilutions in PBS containing 0.1 mg/mL of bovine serum albumin (BSA) were then applied in the wells. The biotinylated sEH nanobody (VHH, variable heavy-chain-only fragment antibody) (1 μg/mL) in PBS was added together to each well immediately to proceed with the immunoreaction for 1 hour at room temperature. Then, the SA-polyHRP in PBS (25 ng/mL) was applied to continue the reaction for another 30 minutes after washing. The 3,3′,5,5′-tetramethylbenzidine (TMB) substrate was added and incubated for 10–15 minutes at room temperature. The reaction was stopped using the color development with 2 M sulfuric acid, and the optical density at 450 nm was quantified using a SpectraMax M2 microplate reader (Molecular Devices) within 10 minutes. Results are average of 3-wells.

***Subcellular Fraction Separation***[11]

HepG2 cells ($8 \times 10^5$ cells) were seeded in 6 wells plate and incubated for 24 hours. After removing the medium, the cells were treated with DMSO or compound **8** (final concentration 100 μM) and incubated for 48 h. Then, the cells were homogenized in 20 mM sodium phosphate buffer (pH 7.4) containing 5 mM EDTA, 1 mM DTT, and 1 mM PMSF with ultrasonic homogenizer. The homogenates were centrifuged at 55,000 rpm (optima TLX ultracentrifuge, TAL-100 rotor, Beckman Coulter Life Sciences) for 15 minutes at 4 °C. The supernatants were collected as S10 fraction, which contains cytosol and microsome. The pellets were resuspended with the same buffer and further sonicated and then centrifuged at

60,000 rpm for 15 minutes at 4 °C. The supernatants were collected as P10 fraction, which contains nuclear, mitochondria, and peroxisome.

***Immunoblotting Assay for Detection of Reducing ER Stress***

HEK293T cells were grown to confluency 24 hours before the experiment, the cell culture media was replaced with fresh media. The cells were pretreated with sEH modulators for 12 hours before being exposed to 1 μM of thapsigargin (Tg) and sEH modulators for an additional 24 hours. The cells were washed with ice-cold PBS and lysed in radio-immunoprecipitation assay (RIPA) buffer. Lysates were clarified by centrifugation at 15,000 g for 10 minutes, and protein concentrations were determined using a bicinchoninic acid assay kit (Pierce Chemical, Dallas, TX). Thirty micrograms of total protein extracts were resolved by sodium dodecyl sulfate polyacrylamide gel electrophoresis (SDS-PAGE) and transferred to polyvinylidene fluoride (PVDF) membranes. Immunoblotting of lysates was performed with primary antibodies (**Table S6**), and after incubation with secondary antibodies, proteins were visualized using Luminata™ Forte Western Chemiluminescent HRP Substrate (Millipore-Sigma). Pixel intensities of immunoblots were quantified using FluorChem Q Imaging software (Alpha Innotech Corp., San Leandro, CA). Data for phosphorylated proteins are presented as the intensity of phosphorylation normalized to its total protein expression.

***Chromatin Condensation Assay***

Cells were grown to confluency. 24 hours before the experiment, the cell culture media was replaced with fresh media. The cells were then pretreated with sEH modulators for 12 hours before being exposed to 1 μM Tg and sEH modulators for an additional 24 hours. The cells were then washed with fresh warm media containing 50 μg/mL Hoechst 33258 and 10 μg/mL propidium iodide to stain apoptotic and necrotic cells, respectively. Using a Leica DMI8 fluorescence microscope (Leica Microsystems Inc., Buffalo Grove, IL), the cells were visualized to identify live cells (low green fluorescence), early apoptotic cells (high green fluorescence), late apoptotic cells (orange fluorescence), and necrotic cells (red fluorescence). For each condition, at least 500 cells were counted. Percentages of apoptotic and necrotic cells were calculated relative to the total number of cells.

***Proteomics Analysis***

HepG2 cells were treated with sEH PROTAC **8** or DMSO vehicle in triplicate. After 24 hours of incubation, the medium was decanted, and the cells were washed with cold phosphate-buffered saline (PBS) and lysed using RIPA buffer with protease inhibitor cocktail (ThermoFisher Scientific). The protein concentration of the lysate was measured using BCA Protein Assay Kit (Thermo Scientific). 500 mM dithiothreitol solution (final concentration 5 mM) was added to the lysate (100 ug protein) and the mixture was incubated at 56 °C for 30 minutes for reduction of cysteine residues. After reduction the reaction, 200 mM 2-iodoacetamide solution (final concentration 20 mM) was added to the mixture, and it was incubated at room temperature for 30 minutes under dark for alkylation of cysteine residues. 12% w/v phosphoric acid ($H_3PO_4$) aqueous solution (final concentration 1.2% w/v) was added to the mixture and the sample was diluted six times with S-trap binding buffer (90% v/v $CH_3OH$ in 10 mM $NH_4HCO_3$ aqueous solution (pH 8.0)). After gentle mixing, the protein solution was loaded onto an S-trap column (Protifi) and centrifuged at 500xg for 30 seconds. The sample was washed three times with the S-trap binding buffer. Finally, 1 μg of sequencing grade trypsin (Promega) in 50 mM $NH_4HCO_3$ aqueous solution (pH 8.0), was added onto the S-trap column and samples were digested at 37 °C for 18 hours. Peptides were eluted in three steps: (i) 80 μL of 50 mM $NH_4HCO_3$ aqueous solution (pH 8.0), (ii) 80 μL of 0.1% v/v TFA aqueous solution and (iii) 80 μL of 60% v/v MeCN and

0.1% v/v TFA aqueous solution. The peptide solution was pooled, centrifuged at 1,000 g for 30 sec and dried in a vacuum centrifuge.

Prior to mass spectrometry analysis, samples were desalted using a 96-well plate filter (Orochem) packed with 1 mg of Oasis HLB C-18 resin (Waters). Briefly, the samples were resuspended in 100 µL of 0.1% TFA and loaded onto the HLB resin, which was previously equilibrated using 100 µL of the same buffer. After washing with 100 µL of 0.1% TFA, the samples were washed with 0.1% TFA, and then eluted with a buffer containing 70 µL of 60% MeCN and 0.1% TFA and then dried in a vacuum centrifuge.

Samples were resuspended in 10 µL of 0.1% TFA and loaded onto a Dionex RSLC Ultimate 3000 (Thermo Scientific), coupled online with an Orbitrap Fusion Lumos (Thermo Scientific). Chromatographic separation was performed with a two-column system, consisting of a C-18 trap cartridge (300 µm ID, 5 mm length) and a picofrit analytical column (75 µm ID, 25 cm length) packed in-house with reversed-phase Repro-Sil Pur C18-AQ 3 µm resin. Peptides were separated using a 90 min gradient from 4-30% buffer B (buffer A: 0.1% formic acid, buffer B: 80% MeCN + 0.1% formic acid) at a flow rate of 300 nl/min. The mass spectrometer was set to acquire spectra in a data-dependent acquisition (DDA) mode. Briefly, the full MS scan was set to 300-1200 m/z in the orbitrap with a resolution of 120,000 (at 200 m/z) and an AGC target of 5x10e5. MS/MS was performed in the ion trap using the top speed mode (2 secs), an AGC target of 1x10e4 and an HCD collision energy of 35.

Proteome raw files were searched using Proteome Discoverer software (v2.5, Thermo Scientific) using SEQUEST search engine and the SwissProt human database (updated April 2023). The search for total proteome included variable modification of N-terminal acetylation, and fixed modification of carbamidomethyl cysteine. Trypsin was specified as the digestive enzyme with up to 2 missed cleavages allowed. Mass tolerance was set to 10 ppm for precursor ions and 0.2 Da for product ions. Peptide and protein false discovery rate was set to 1%. Following the search, data was processed as described by Aguilan, J.T. *et al. Mol. Omics* **2020**.[12] Briefly, proteins were $\log_2$ transformed, normalized by the average value of each sample and missing values were imputed using a normal distribution 2 standard deviations lower than the mean. Statistical regulation was assessed using heteroscedastic t-test (if $p$-value < 0.05). Data distribution was assumed to be normal but this was not formally tested.

***Data Analysis***

Descriptive statistics were obtained and include the mean and standard error of mean for continuous variables and count and proportion for categorical variables. Pairwise comparisons in changes in protein levels was performed and include Student's t-test for continuous variables. Comparisons across all groups were made via an ANOVA. The details of statistical analyses are given in the figure legend.

NMR spectra

ALT-PG2

**[1]H NMR (600 MHz, DMSO-$d_6$)**

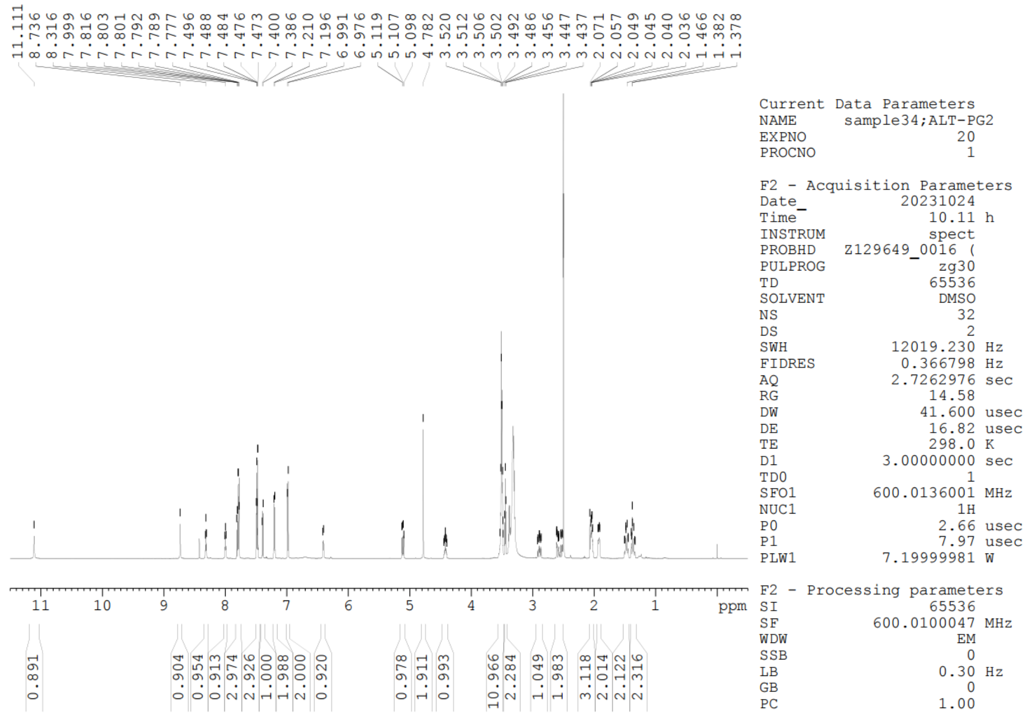

**[19]F NMR (565 MHz, DMSO-$d_6$)** $CFCl_3$ was used as an internal standard.

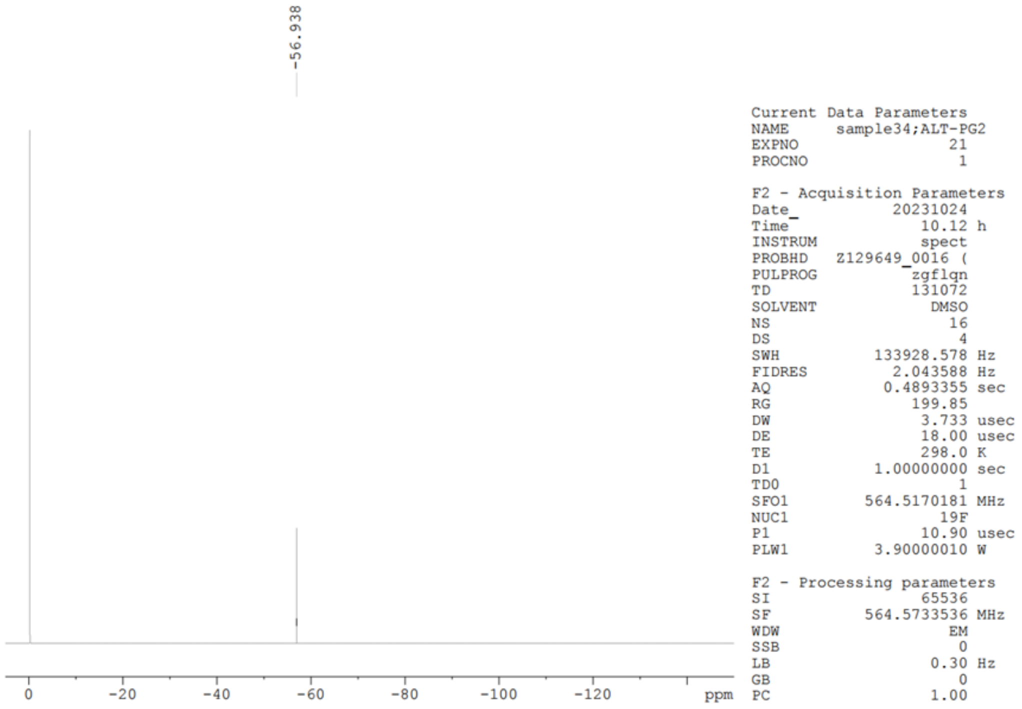

**Compound 1**

**$^{1}$H NMR (600 MHz, $CD_3OD$)**

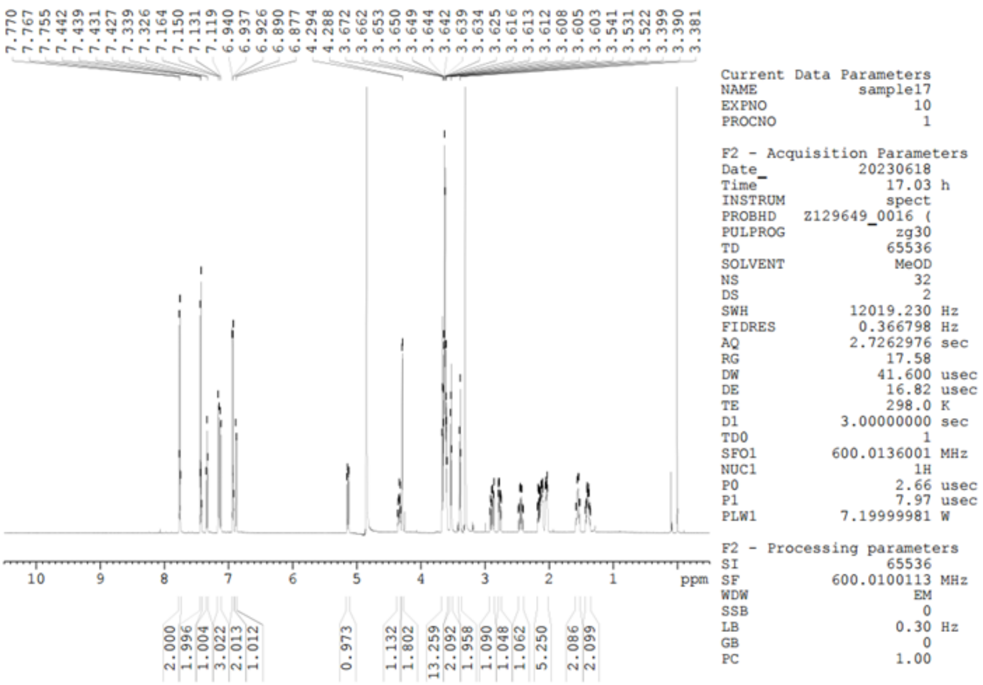


**$^{13}$C NMR (151 MHz, $CD_3OD$)**

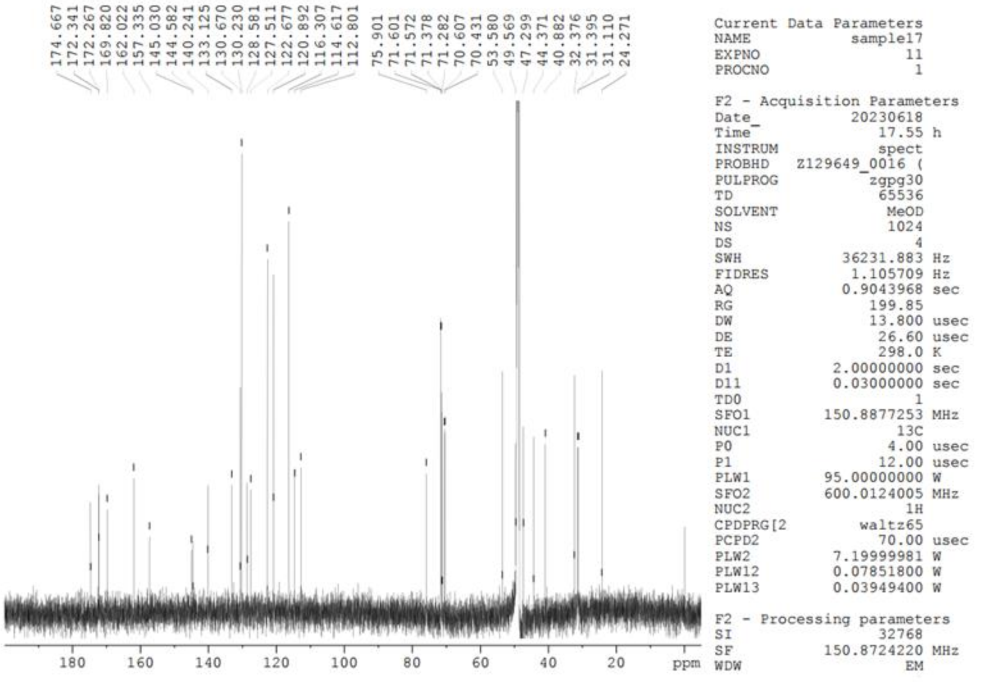

**Compound 1**

**$^{19}$F NMR (565 MHz, $CD_3OD$).** $CFCl_3$ was used as an internal standard.

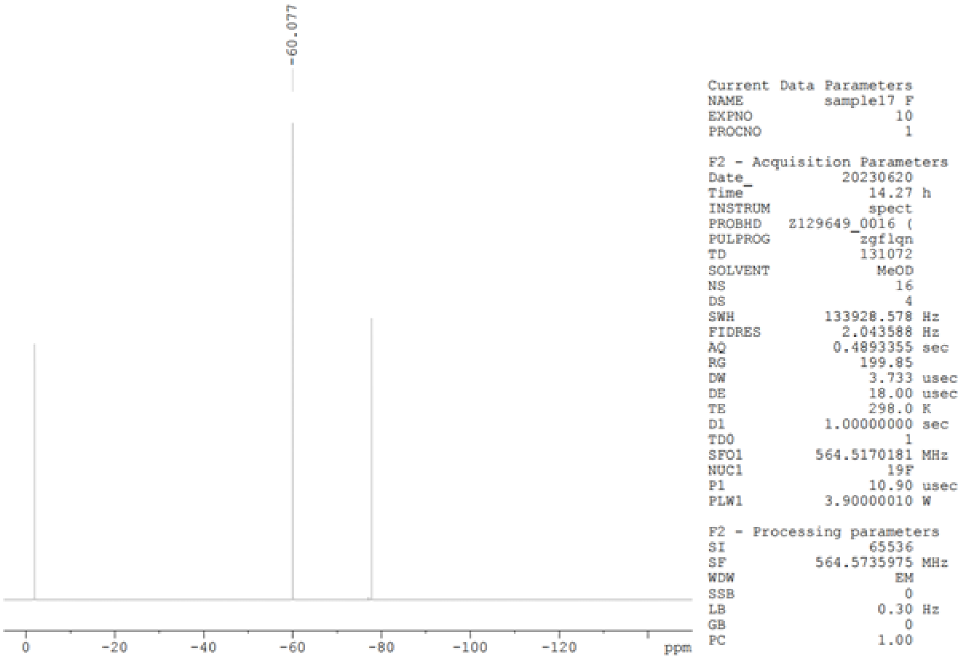

**Compound 2**

**$^{1}H$ NMR (600 MHz, $CD_3OD$)**

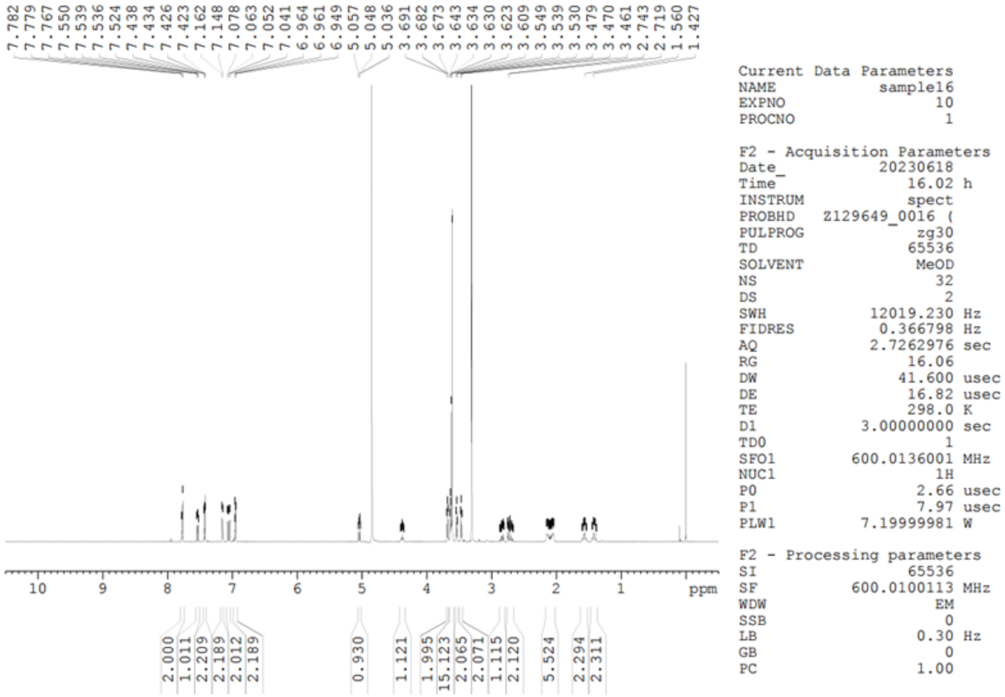


**$^{13}C$ NMR (151 MHz, $CD_3OD$)**

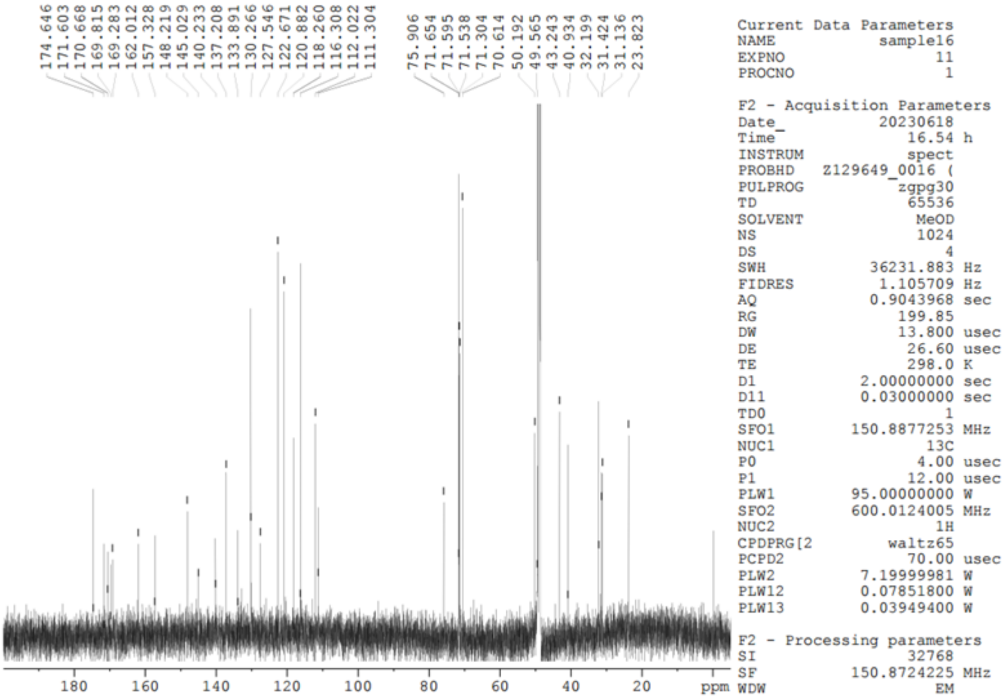

**Compound 2**

**$^{19}$F NMR (565 MHz, $CD_3OD$)** TFA was used as an internal standard.

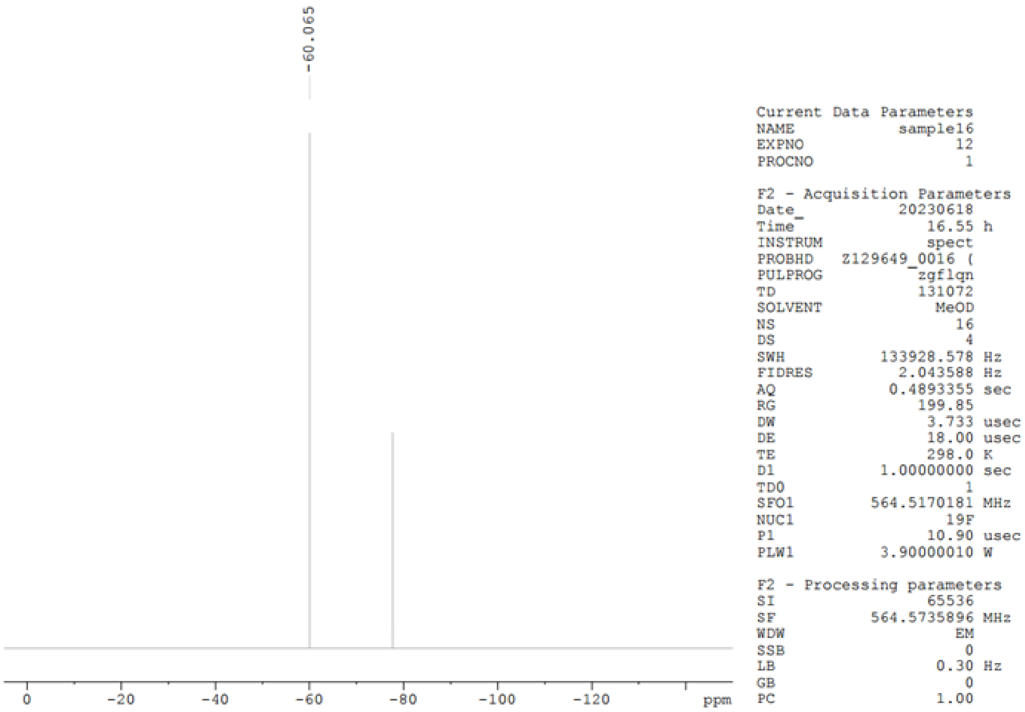

**Compound 3**

**$^{1}$H NMR (600 MHz, $CD_3OD$)**

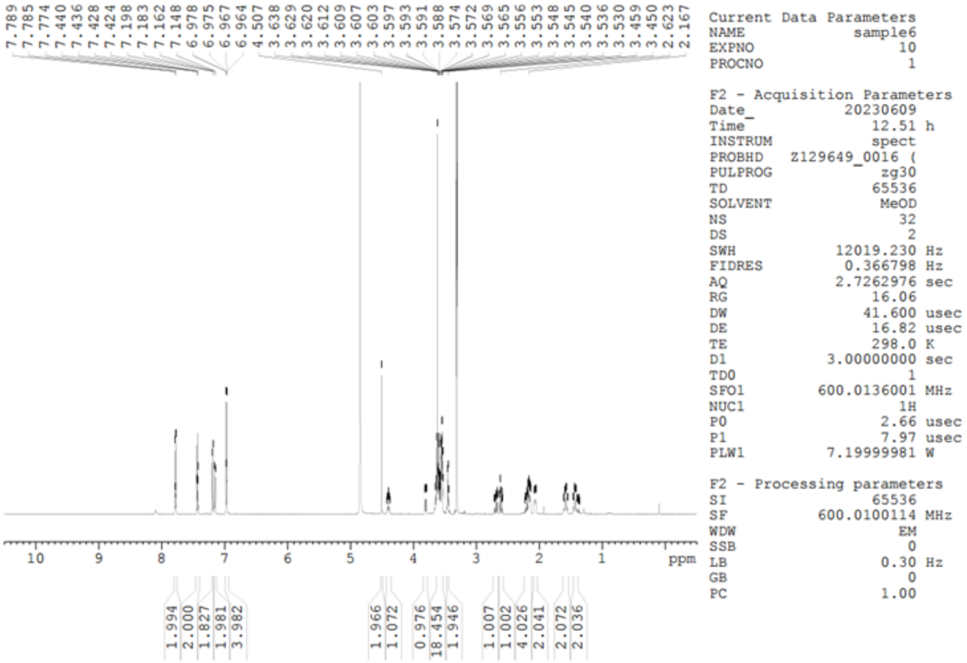


**$^{13}$C NMR (151 MHz, $CD_3OD$)**

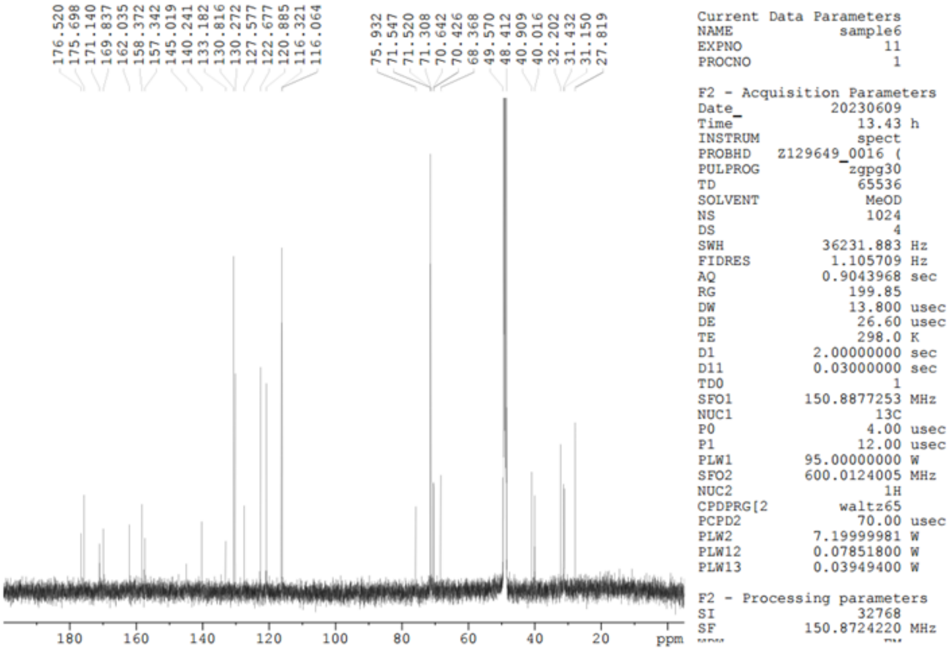

**Compound 3**

**$^{19}$F NMR (565 MHz, $CD_3OD$)** $CFCl_3$ was used as an internal standard.

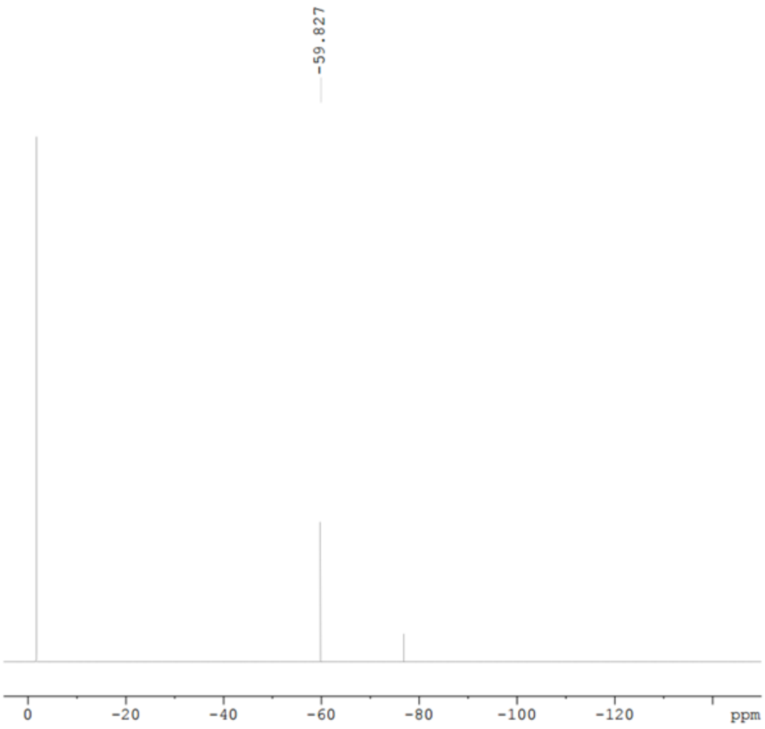


```
Current Data Parameters
NAME          sample6 F
EXPNO                11
PROCNO                1

F2 - Acquisition Parameters
Date_          20230620
Time              20.30 h
INSTRUM           spect
PROBHD   Z129649_0016 (
PULPROG          zgflqn
TD               131072
SOLVENT            MeOD
NS                   16
DS                    4
SWH          133928.578 Hz
FIDRES         2.043588 Hz
AQ            0.4893355 sec
RG               199.85
DW                3.733 usec
DE                18.00 usec
TE                298.0 K
D1           1.00000000 sec
TD0                   1
SFO1        564.5170181 MHz
NUC1                19F
P1                10.90 usec
PLW1         3.90000010 W

F2 - Processing parameters
SI                65536
SF          564.5734571 MHz
WDW                  EM
SSB                   0
LB                 0.30 Hz
GB                    0
PC                 1.00
```

**Compound 4**

**$^{1}$H NMR (300 MHz, $CD_3OD$)**

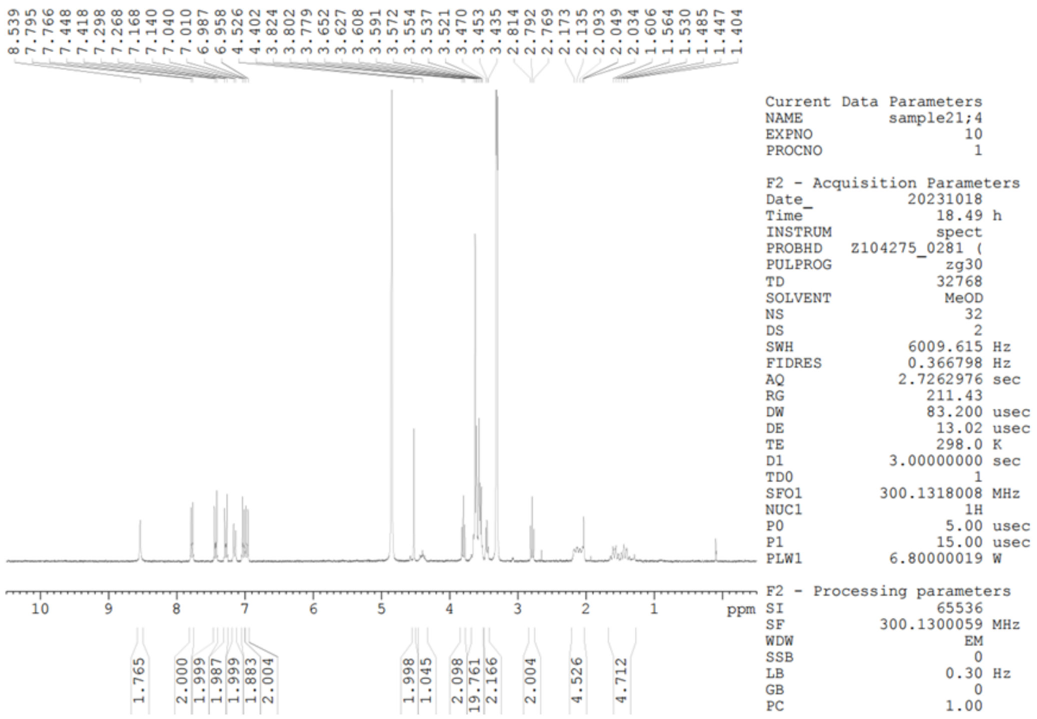

**$^{13}$C NMR (151 MHz, $CD_3OD$)**

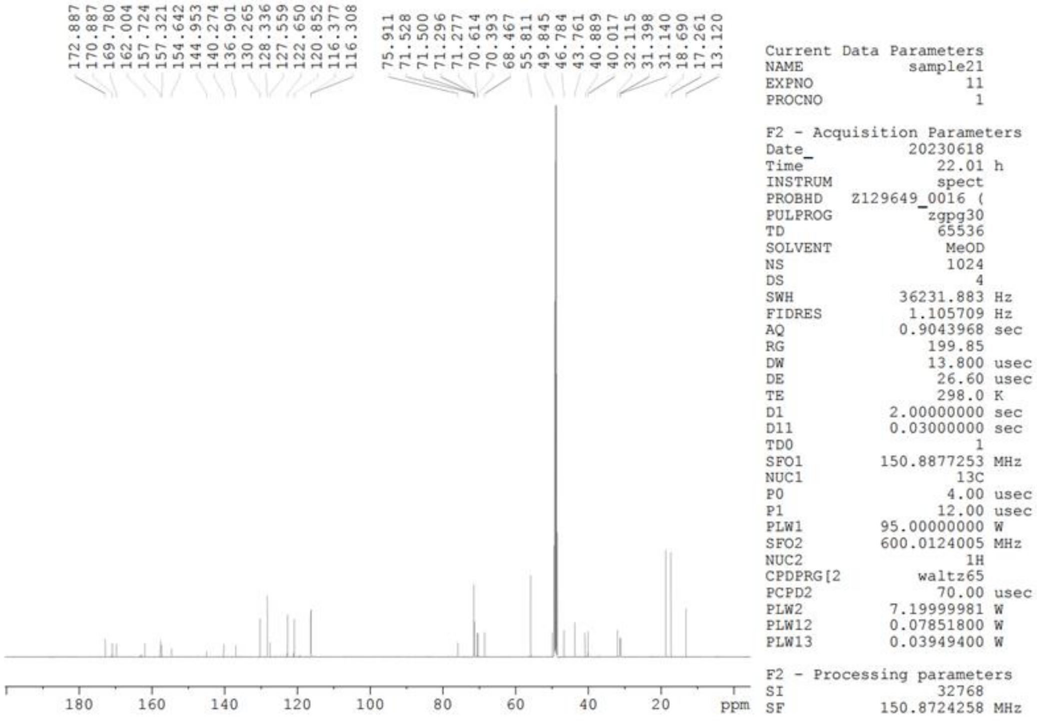

**Compound 4**

**$^{19}$F NMR (565 MHz, $CD_3OD$)** TFA was used as an internal standard.

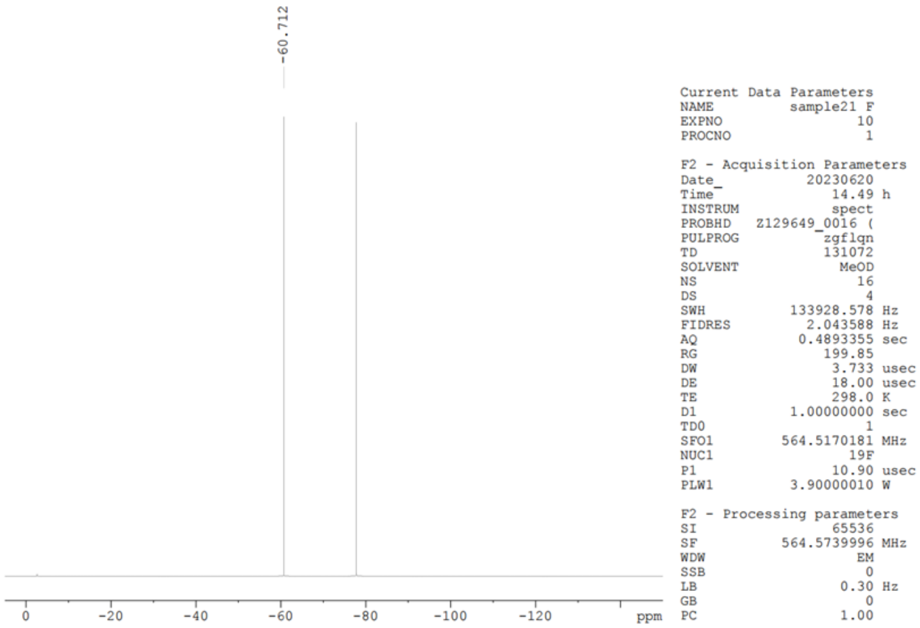

**Compound 5**

**$^1$H NMR (300 MHz, $CD_3OD$)**

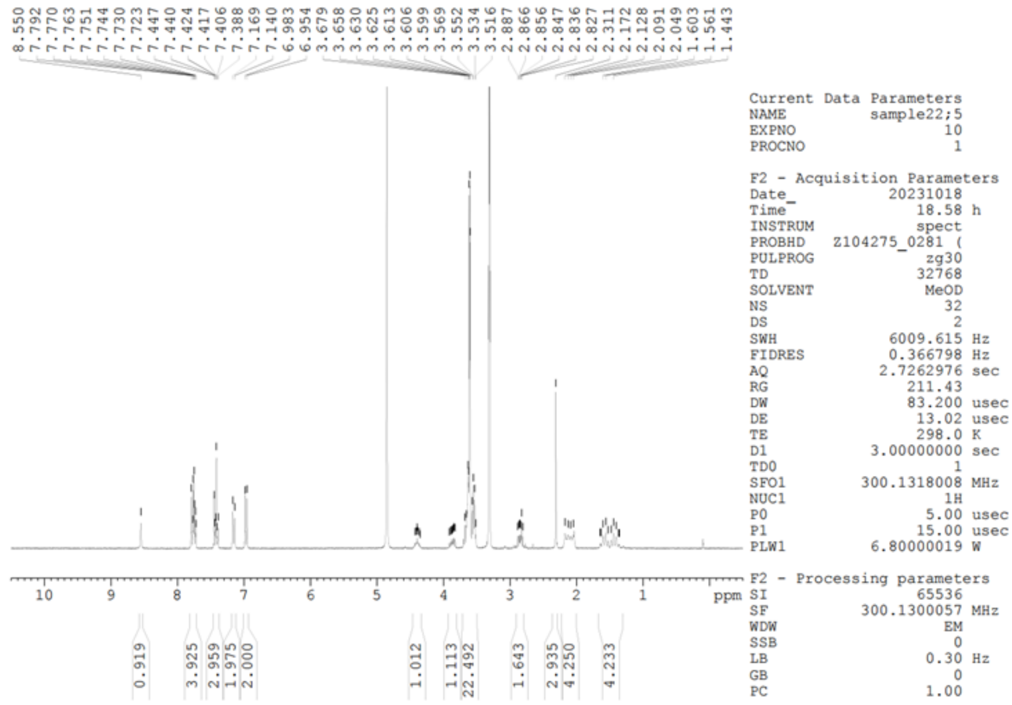


**$^{13}$C NMR (151 MHz, $CD_3OD$)**

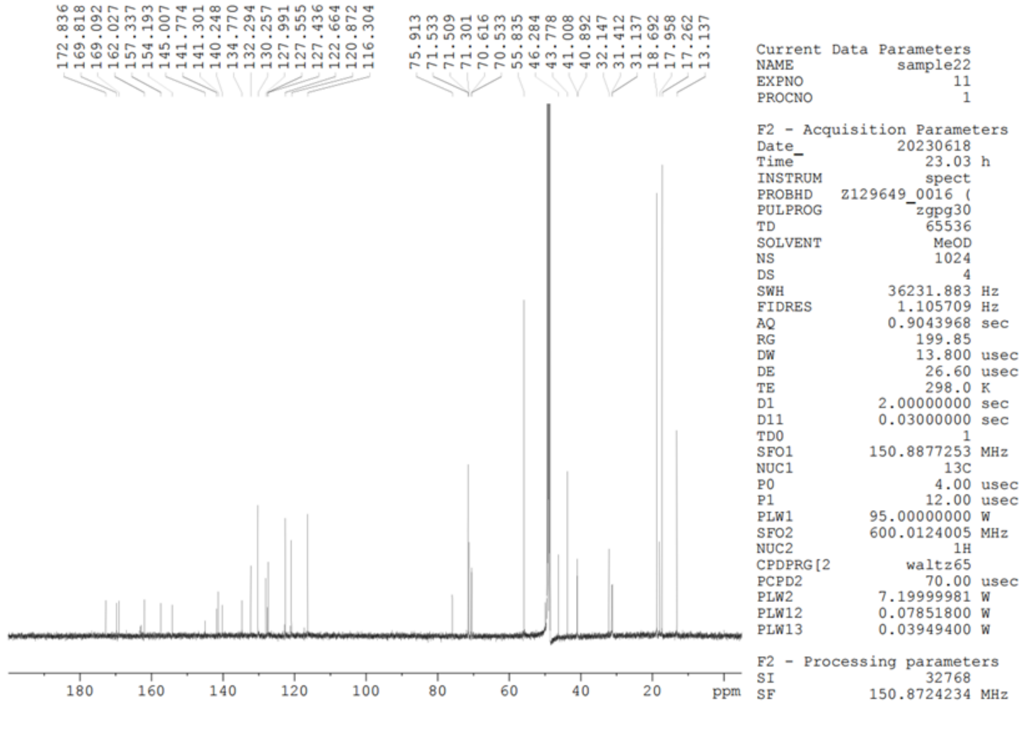

**Compound 5**

**$^{19}$F NMR (565 MHz, $CD_3OD$)** $CFCl_3$ was used as an internal standard.

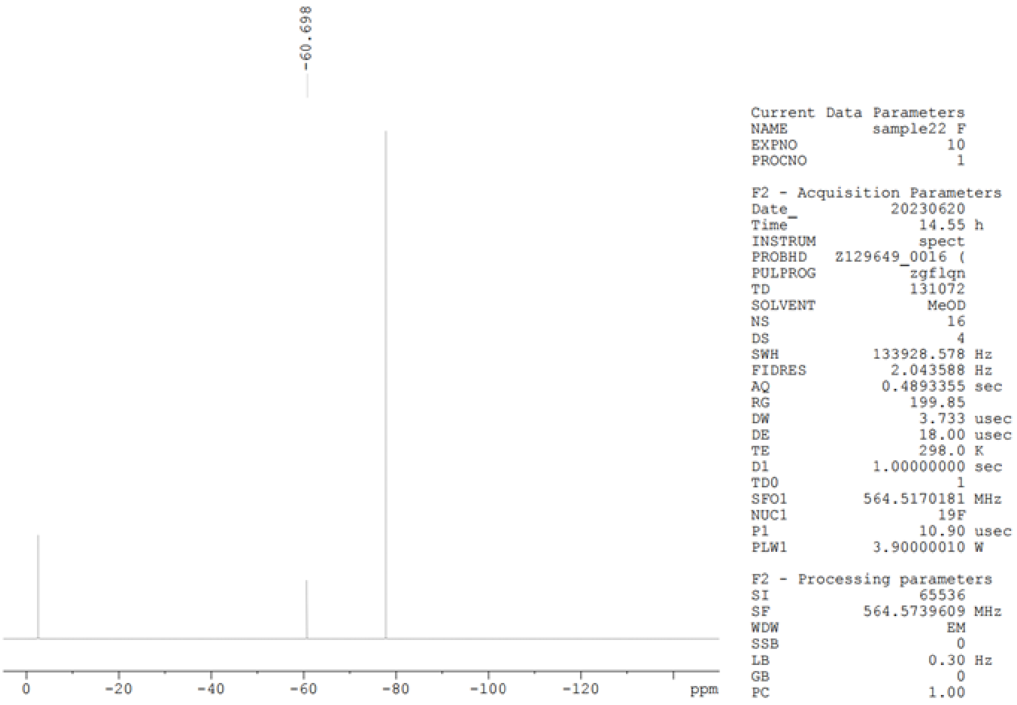

**Compound 6**

**$^{1}$H NMR (600 MHz, $CD_3OD$)**

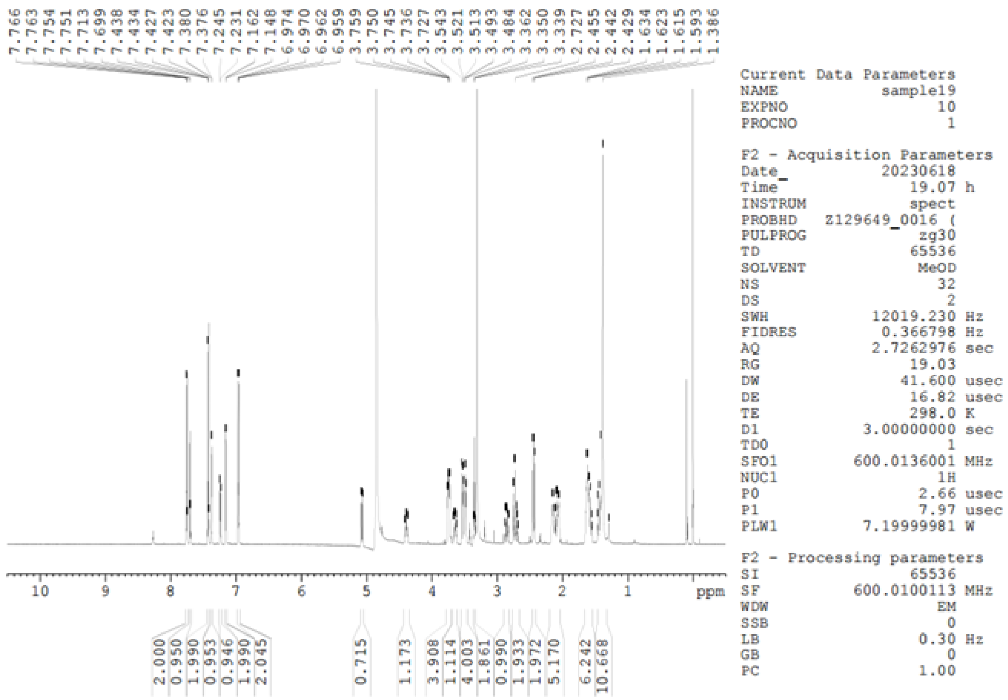


**$^{13}$C NMR (75 MHz, $CD_3OD$)**

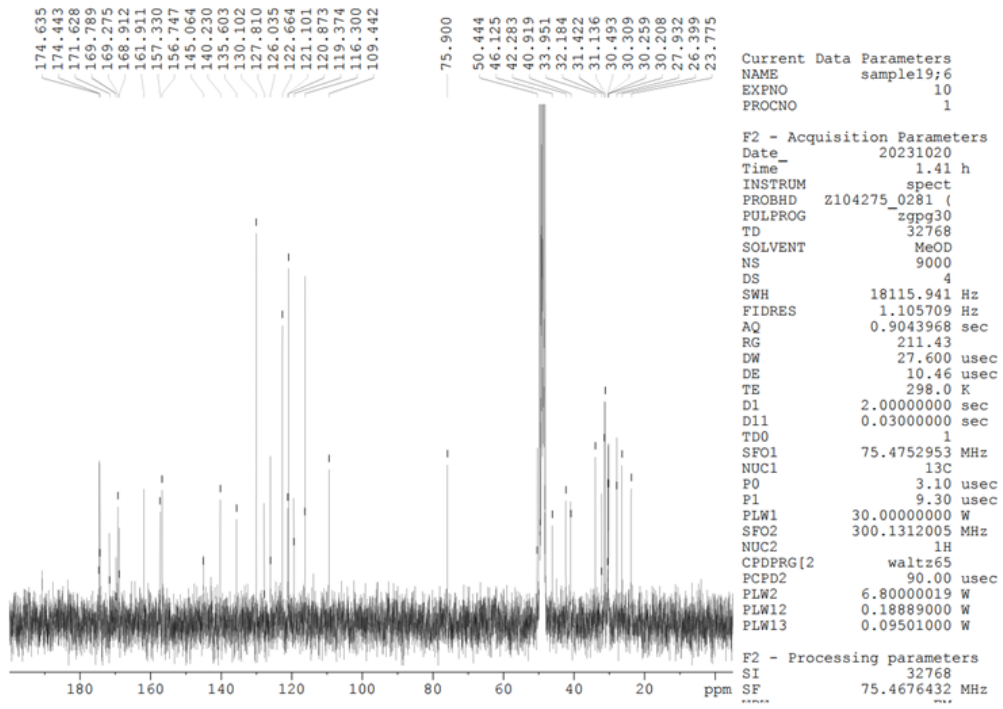

**Compound 6**

**$^{19}$F NMR (565 MHz, $CD_3OD$)** $CFCl_3$ was used as an internal standard.

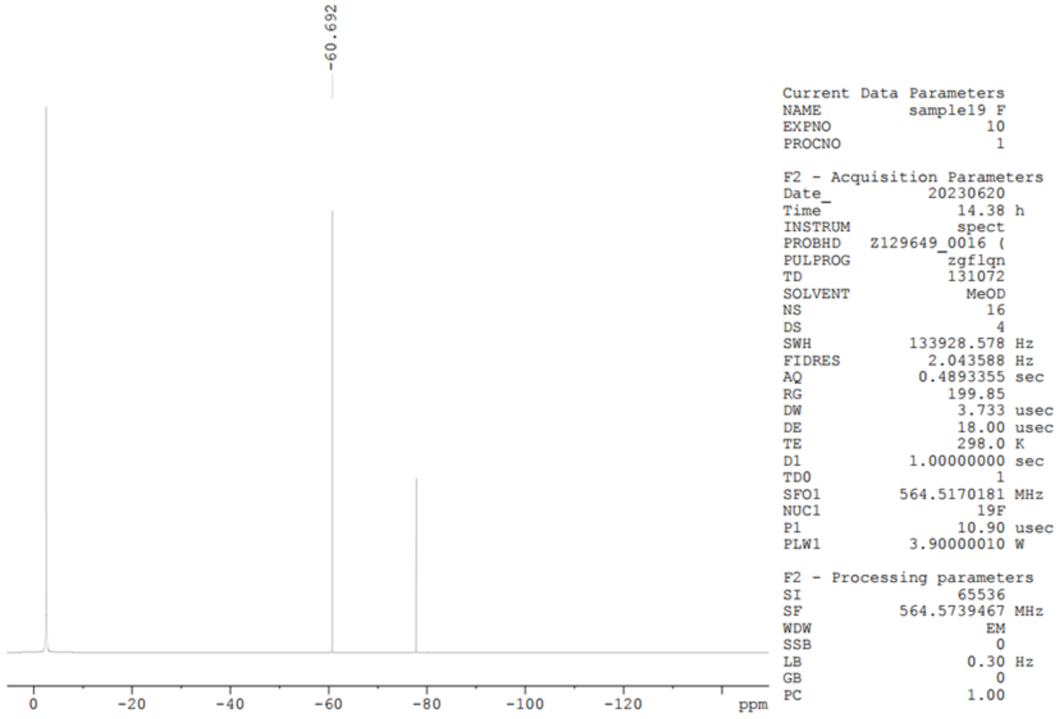

**Compound 7**

**$^{1}$H NMR (600 MHz, $CD_3OD$)**

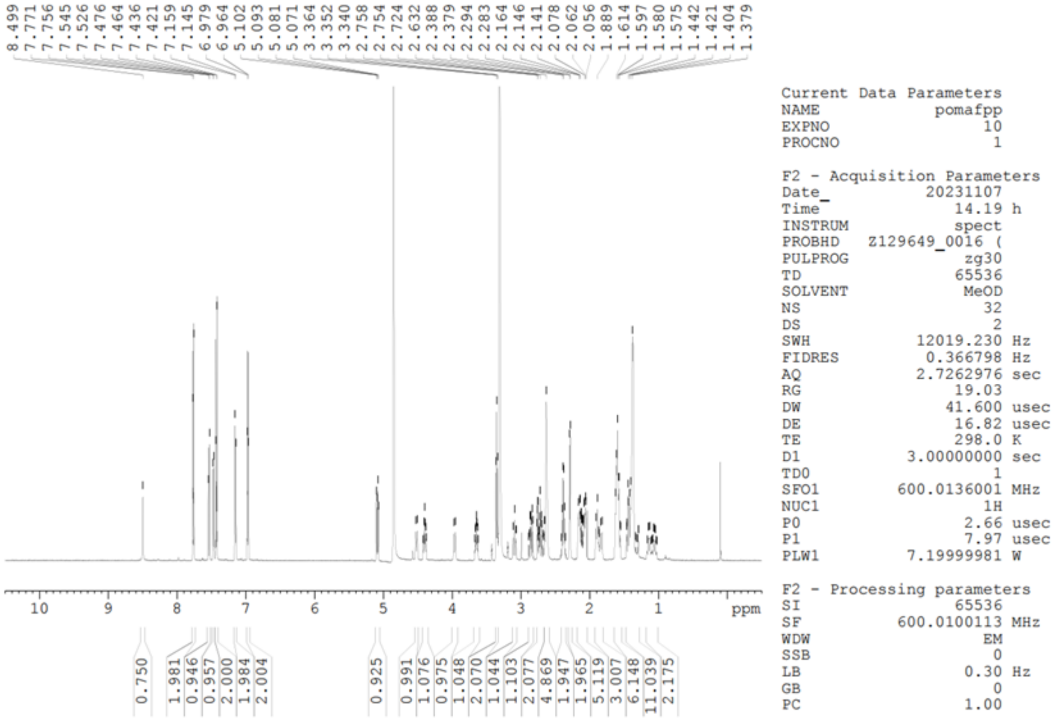


**$^{13}$C NMR (151 MHz, $CD_3OD$)**

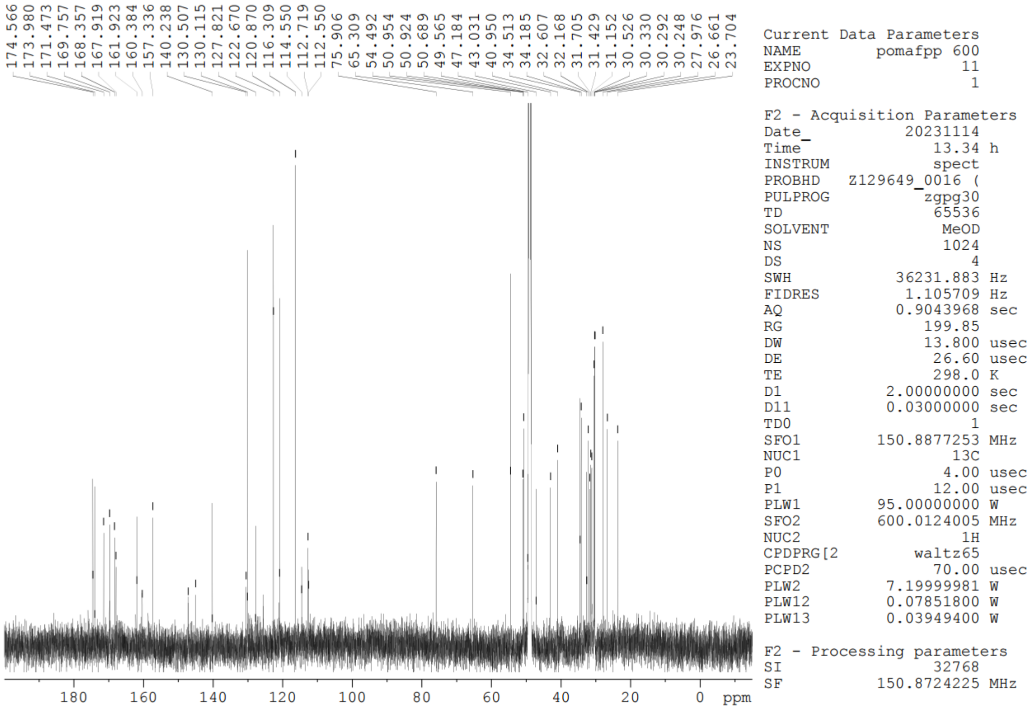

**Compound 7**

**$^{19}$F NMR (565 MHz, $CD_3OD$)** $CFCl_3$ was used as an internal standard.

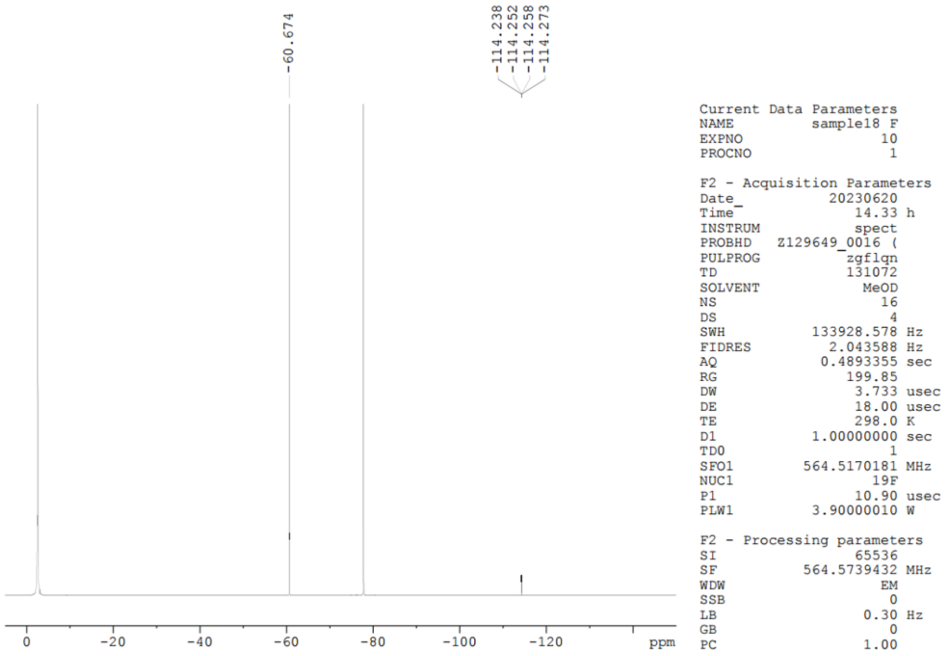


**Compound 7**

**$^{19}$F NMR (565 MHz, $CD_3OD$)** $CFCl_3$ was used as an internal standard.

**(-113.50 - -115.00 ppm)**

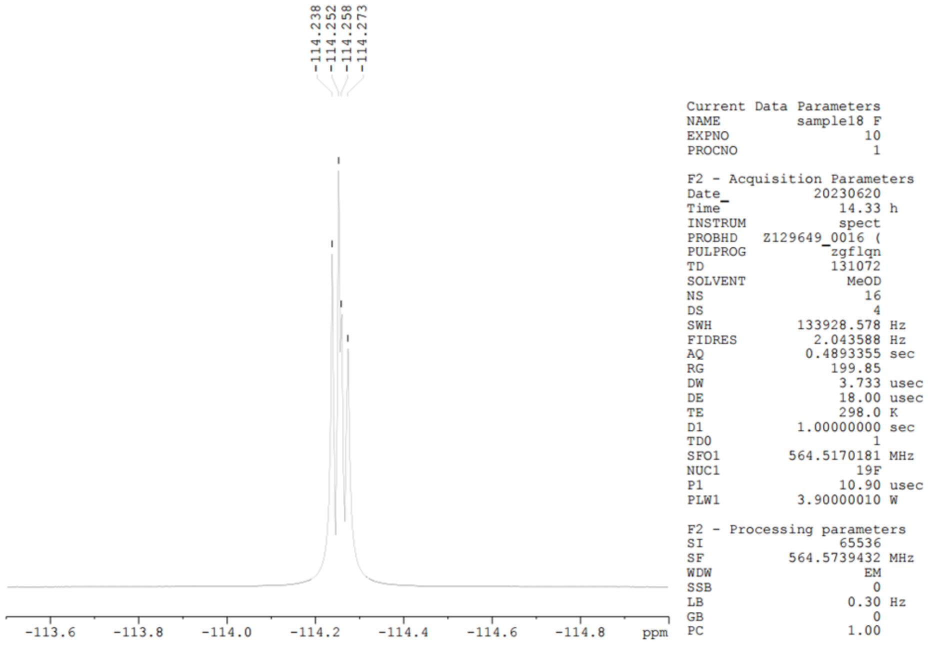

**Compound 8**

**$^{1}$H NMR (600 MHz, $CD_3OD$)**

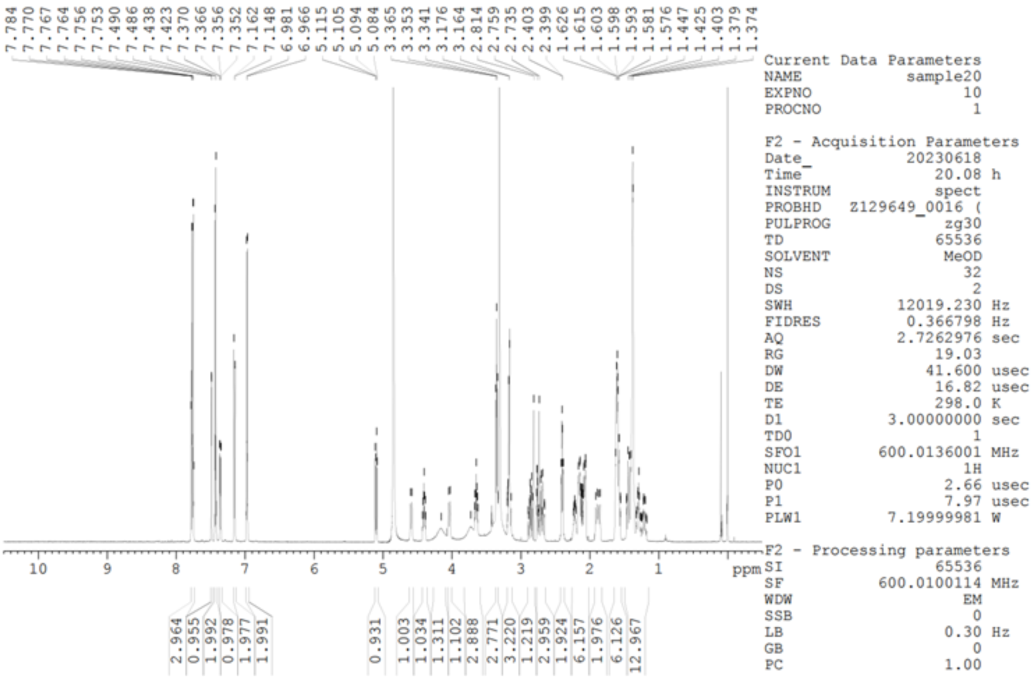


**$^{13}$C NMR (151 MHz, $CD_3OD$)**

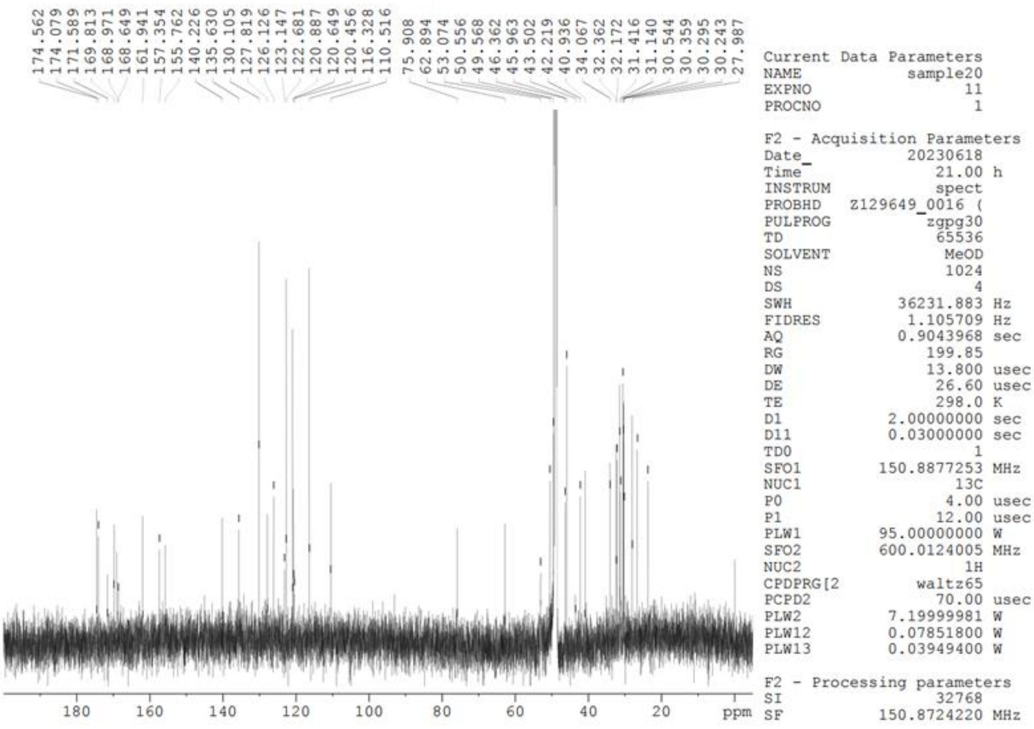

**Compound 8**

**$^{19}$F NMR (565 MHz, $CD_3OD$)** $CFCl_3$ was used as an internal standard.

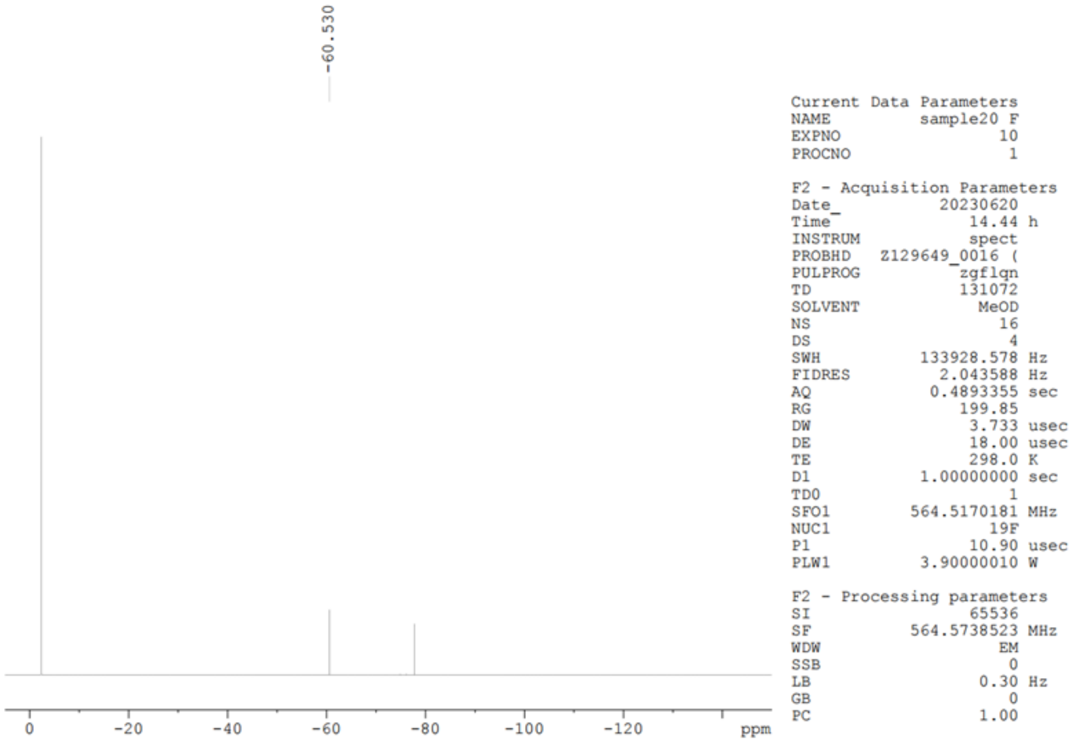

**Compound 14**

**$^1$H NMR (600 MHz, DMSO-$d_6$)**

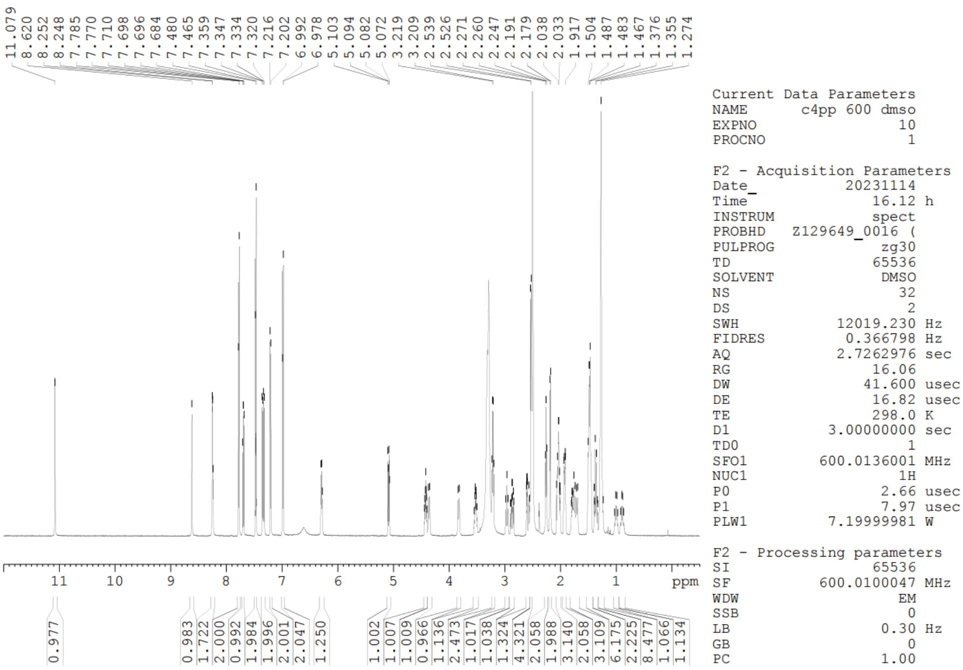

**$^{13}$C NMR (151 MHz, DMSO-$d_6$)**

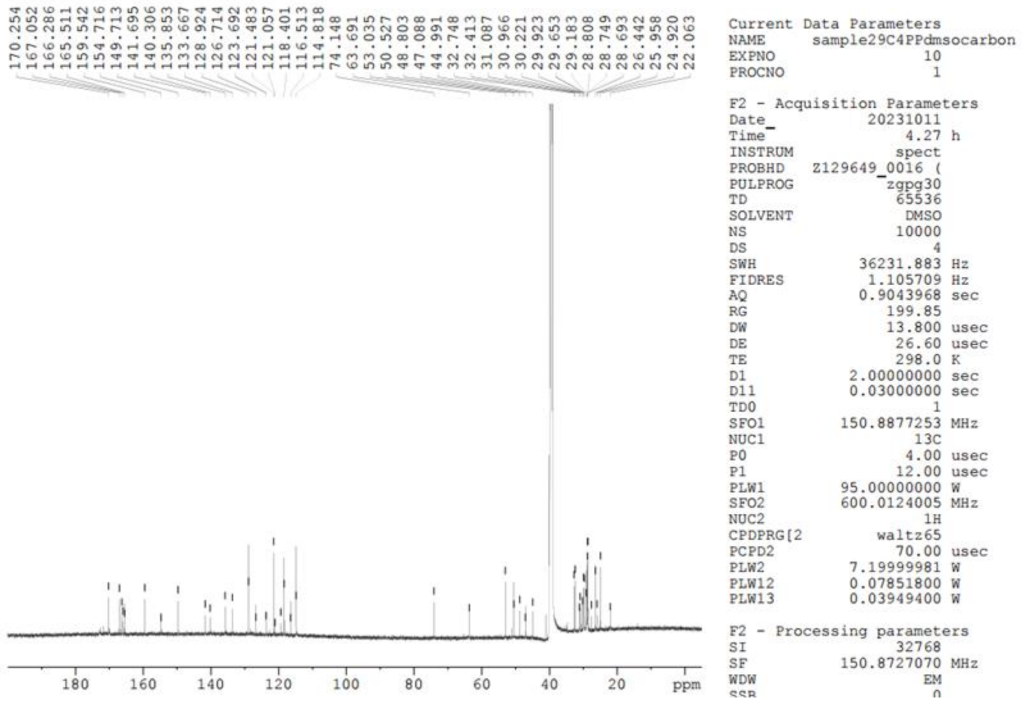

**Compound 14**

**$^{19}$F NMR (282 MHz, DMSO-$d_6$)** $CFCl_3$ was used as an internal standard.

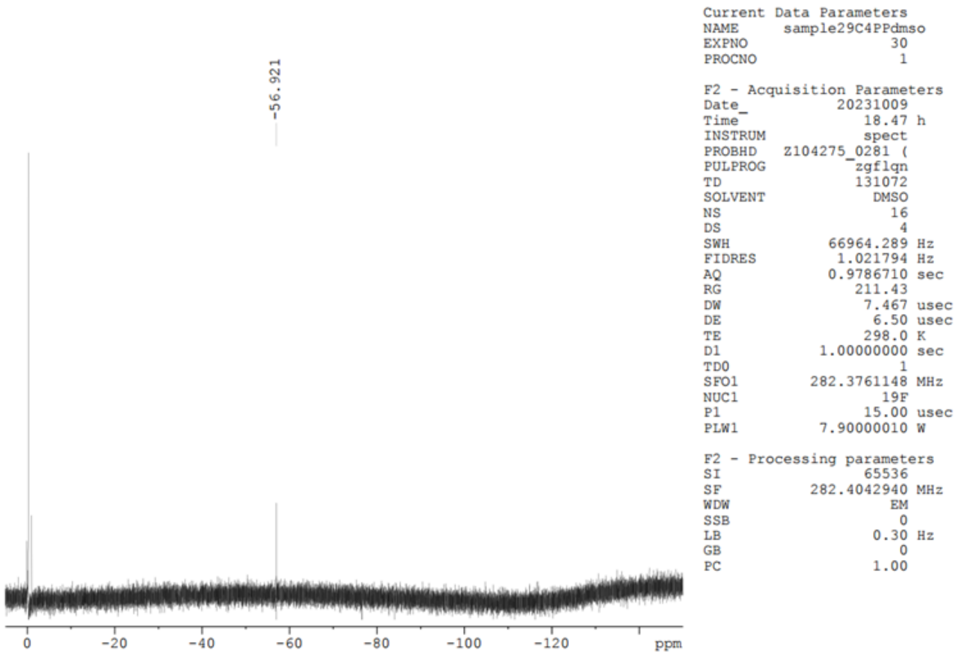

**Compound 15**

**$^{1}$H NMR (600 MHz, $CD_3OD$)**

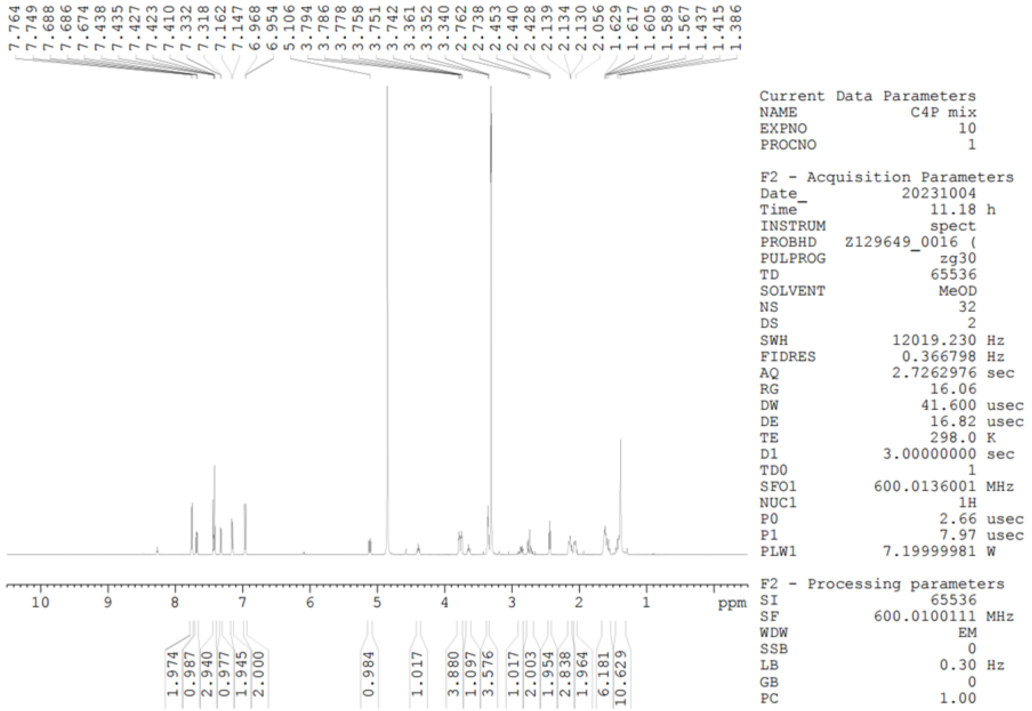


**$^{13}$C NMR (75 MHz, $CD_3OD$)**

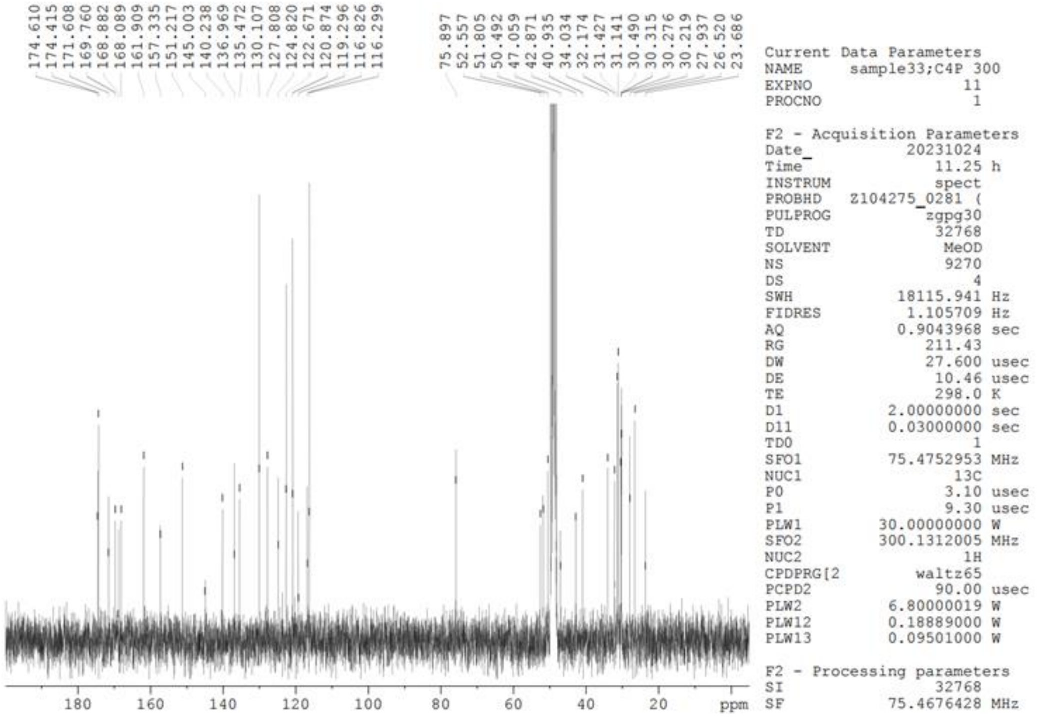

**Compound 15**

**$^{19}$F NMR (282 MHz, DMSO-$d_6$)** TFA was used as an internal standard.

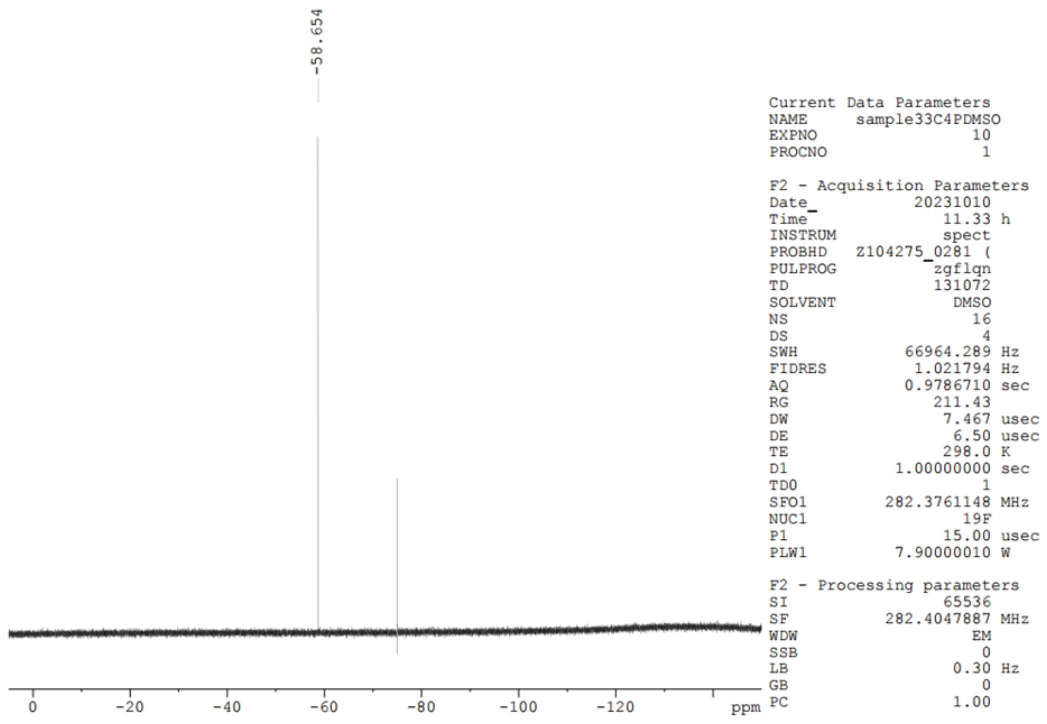

**Compound 16**

**$^{1}$H NMR (600 MHz, $CD_3OD$)**

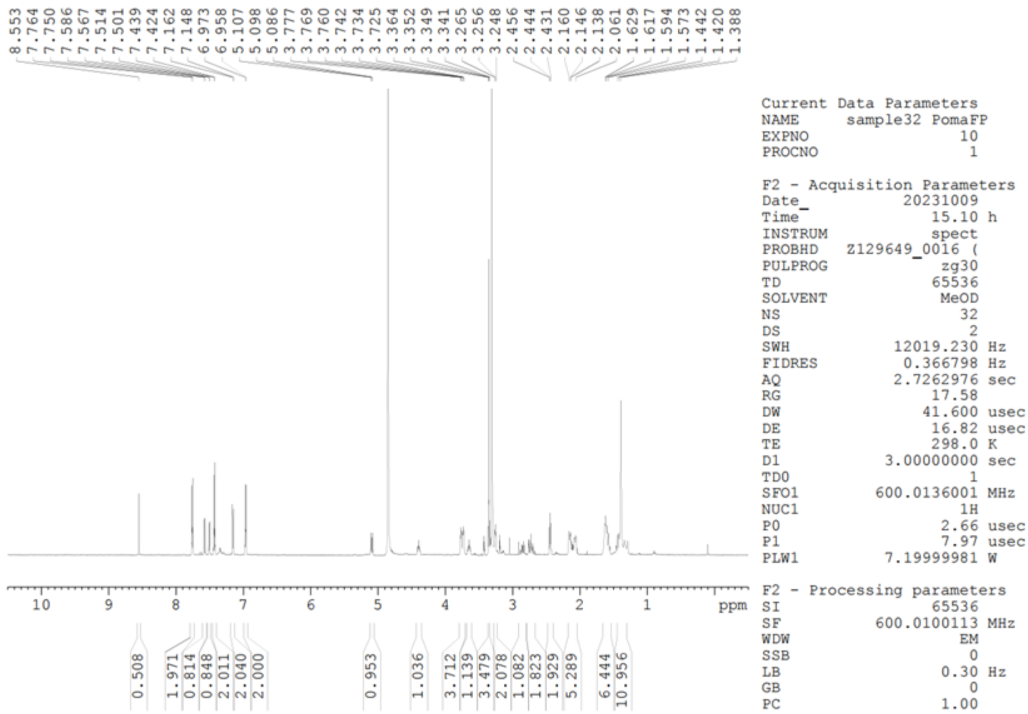

**$^{13}$C NMR (151 MHz, DMSO-$d_6$)**

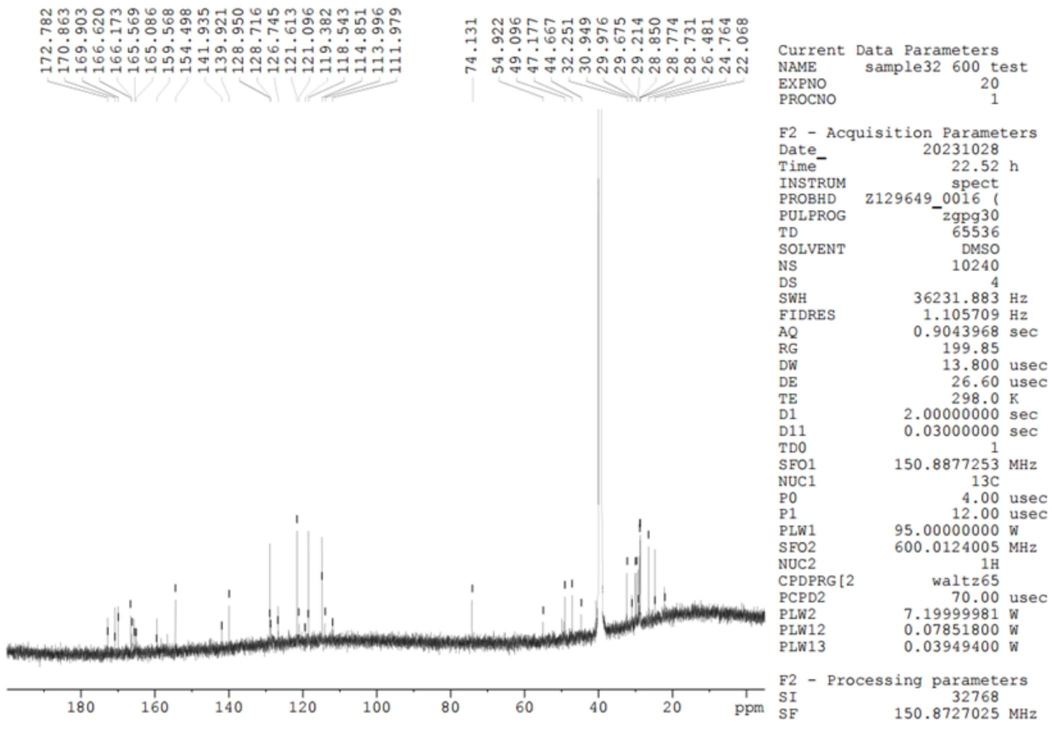

**Compound 16**

**$^{19}$F NMR (565 MHz, $CD_3OD$)** TFA was used as an internal standard.

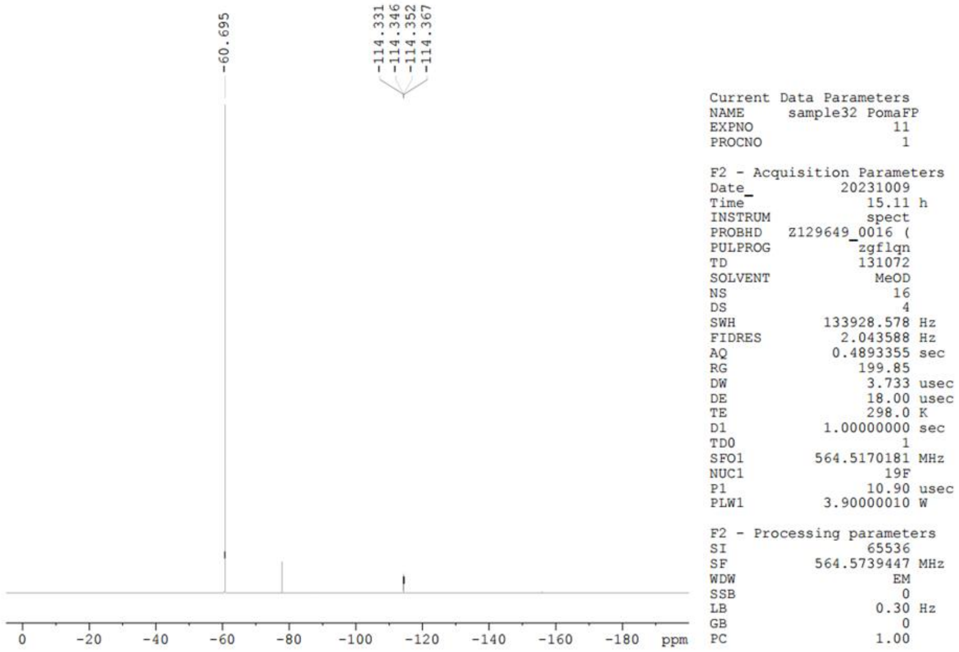


**Compound 16**

**$^{19}$F NMR (565 MHz, $CD_3OD$)**

**(-113.50 - -115.00 ppm)**

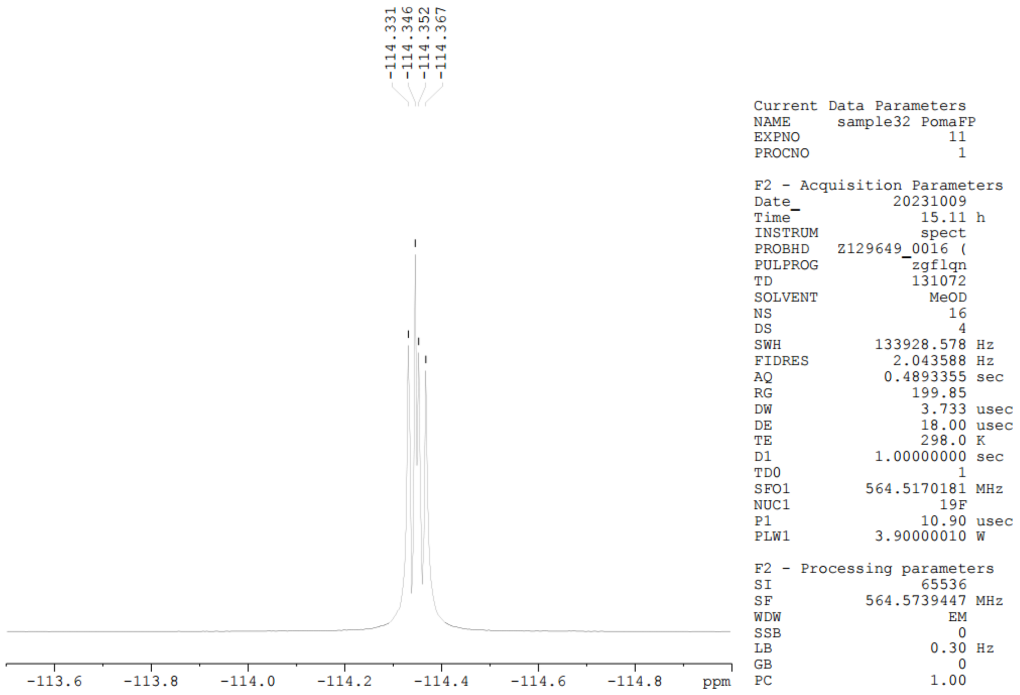

**Compound 17**

**$^{1}H$ NMR (600 MHz, $CD_3OD$)**

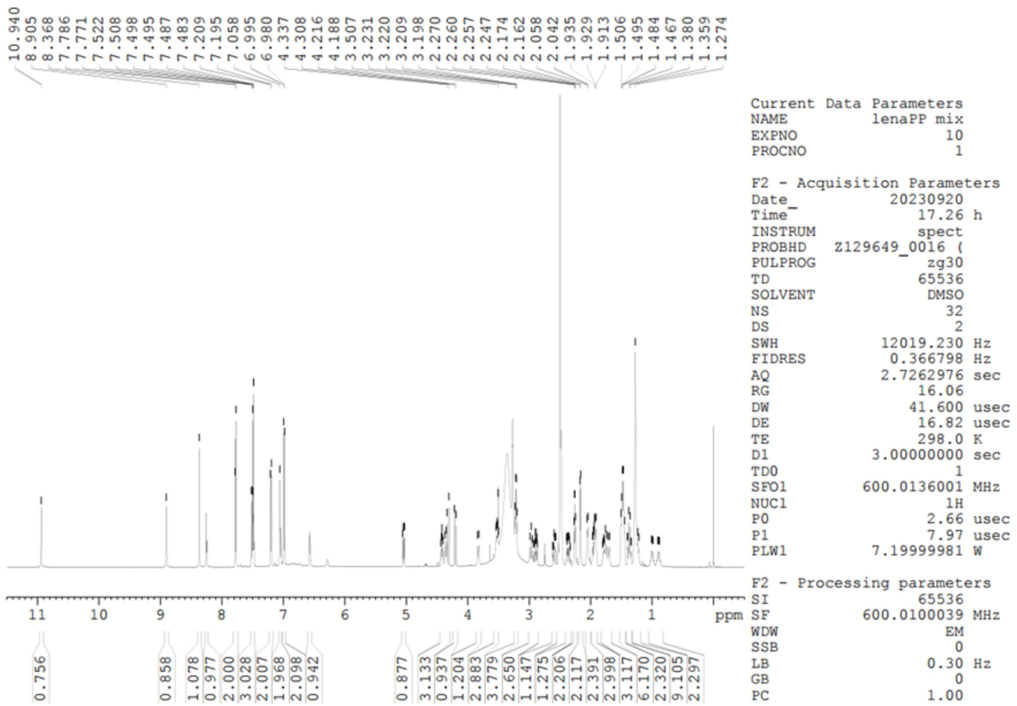


**$^{13}C$ NMR (151 MHz, $CD_3OD$)**

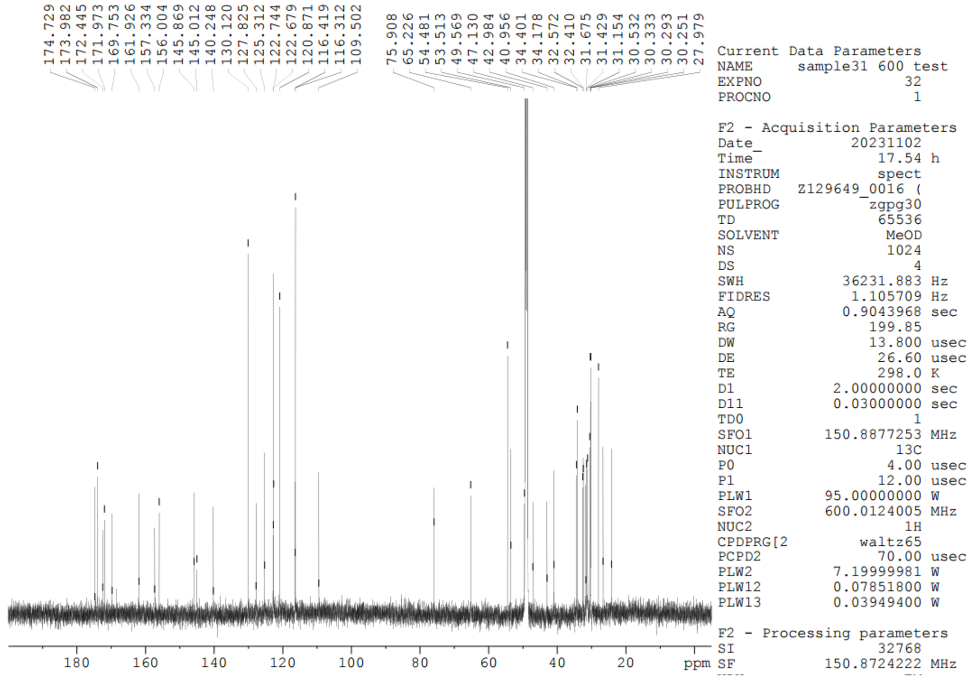

**Compound 17**

**[19]F NMR (565 MHz, $CD_3OD$)** TFA was used as an internal standard.

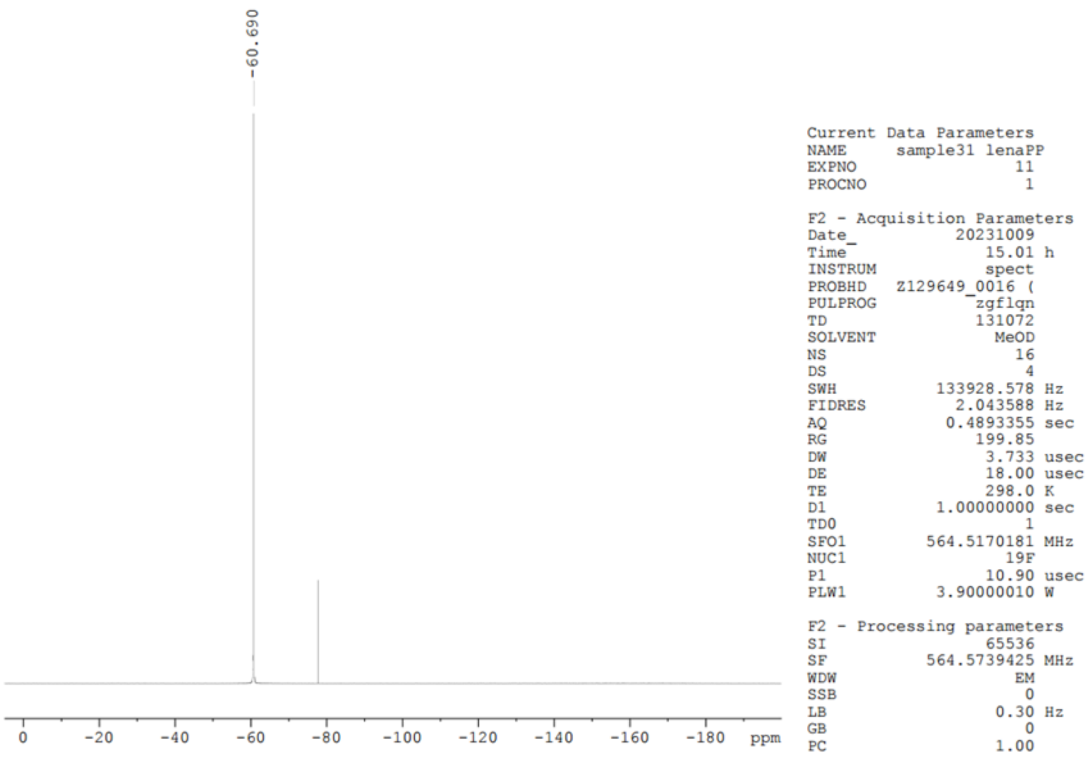

**Compound 18**

**$^{1}$H NMR (600 MHz, DMSO-$d_6$)**

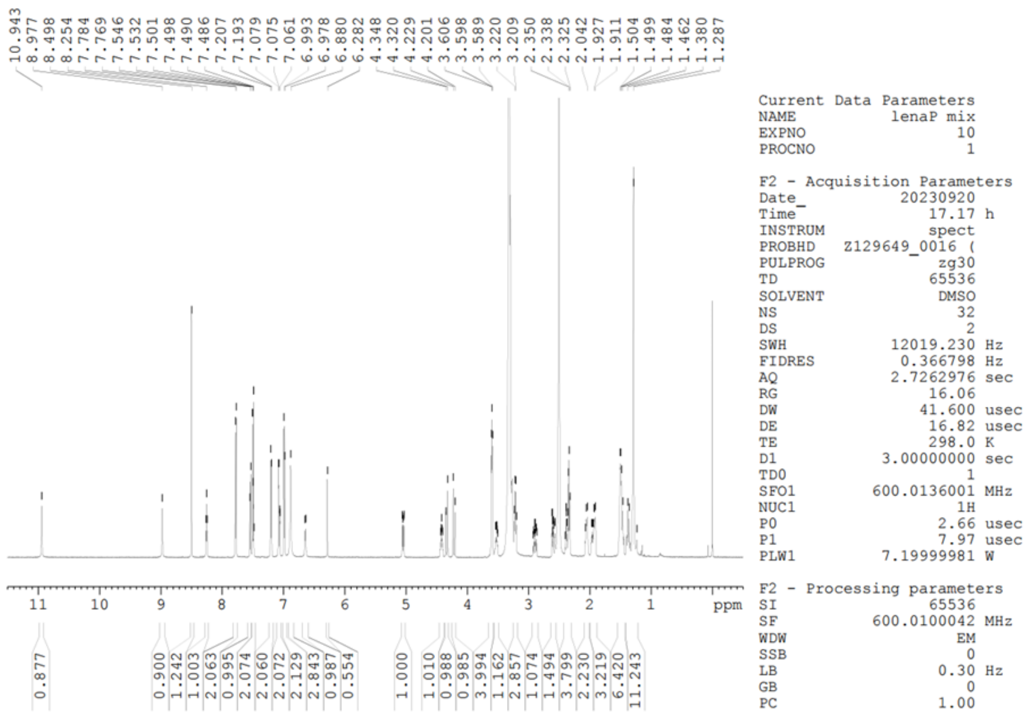


**$^{13}$C NMR (151 MHz, $CD_3OD$)**

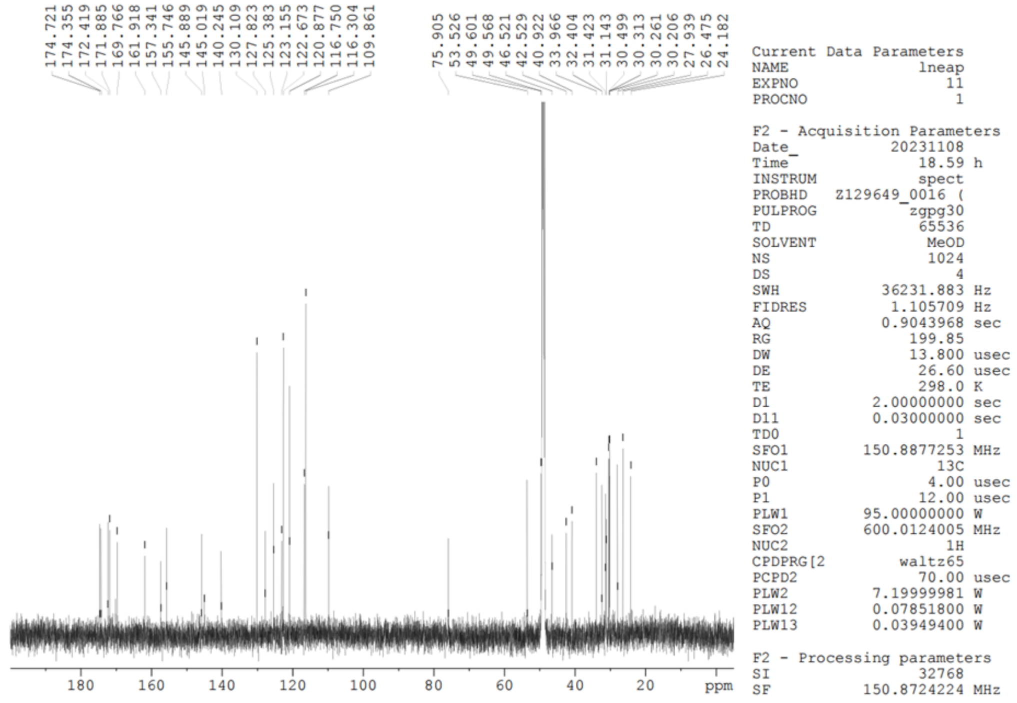

**Compound 18**

**$^{19}$F NMR (565 MHz, $CD_3OD$)** $CFCl_3$ was used as an internal standard.

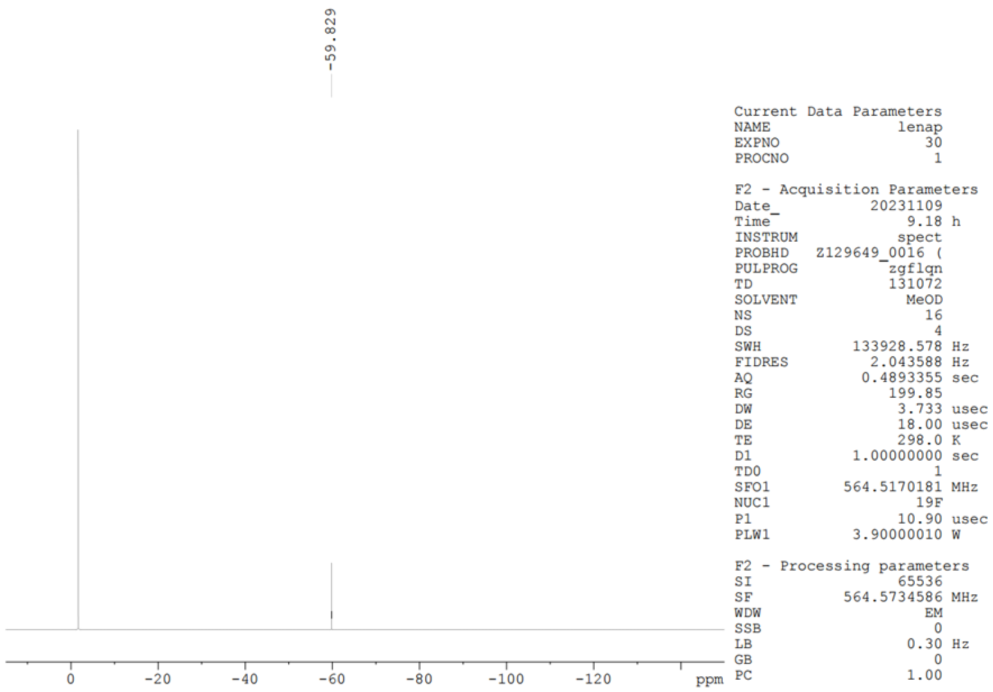

**Compound 9**

**$^{1}H$ NMR (600 MHz, $CD_3OD$)**

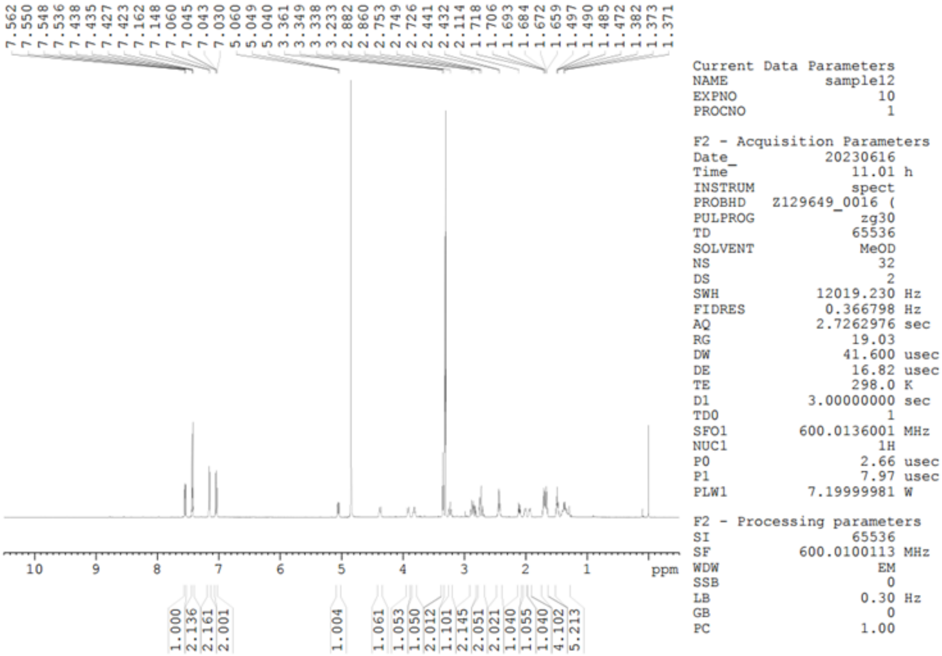


**$^{13}C$ NMR (151 MHz, $CD_3OD$)**

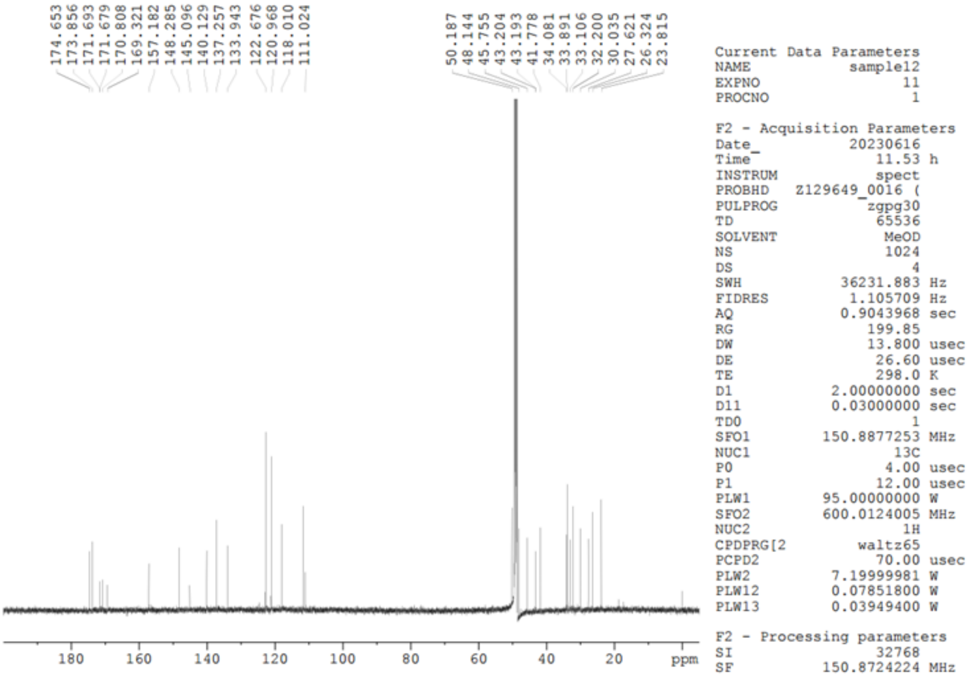

**Compound 9**

**$^{19}$F NMR (565 MHz, $CD_3OD$)** TFA was used as an internal standard.

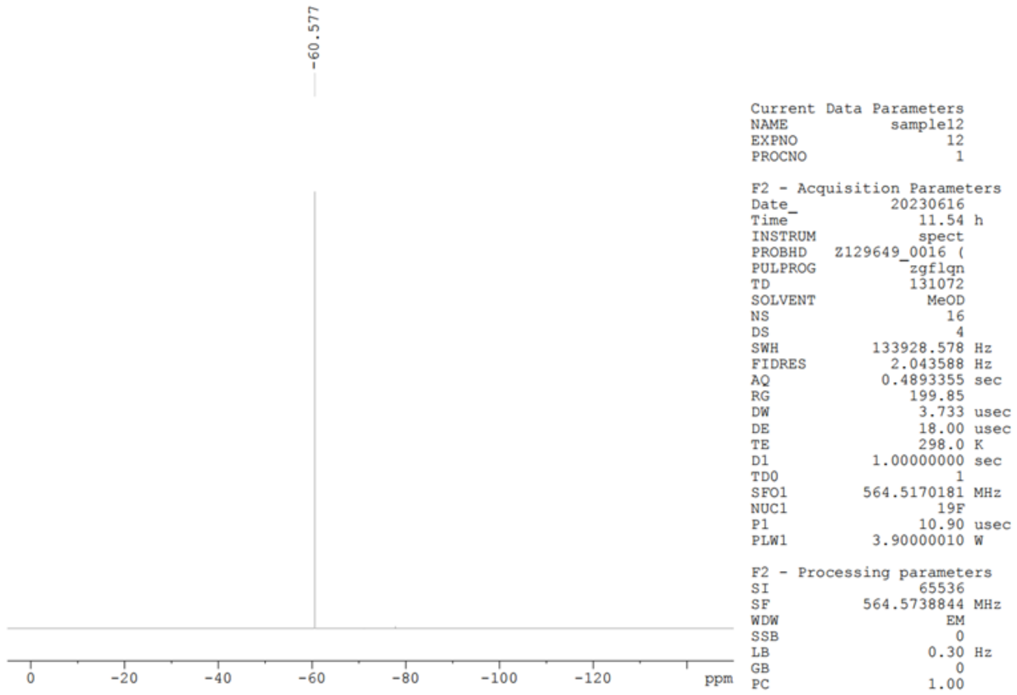

**Compound 10**

**$^1$H NMR (300 MHz, $CD_3OD$)**

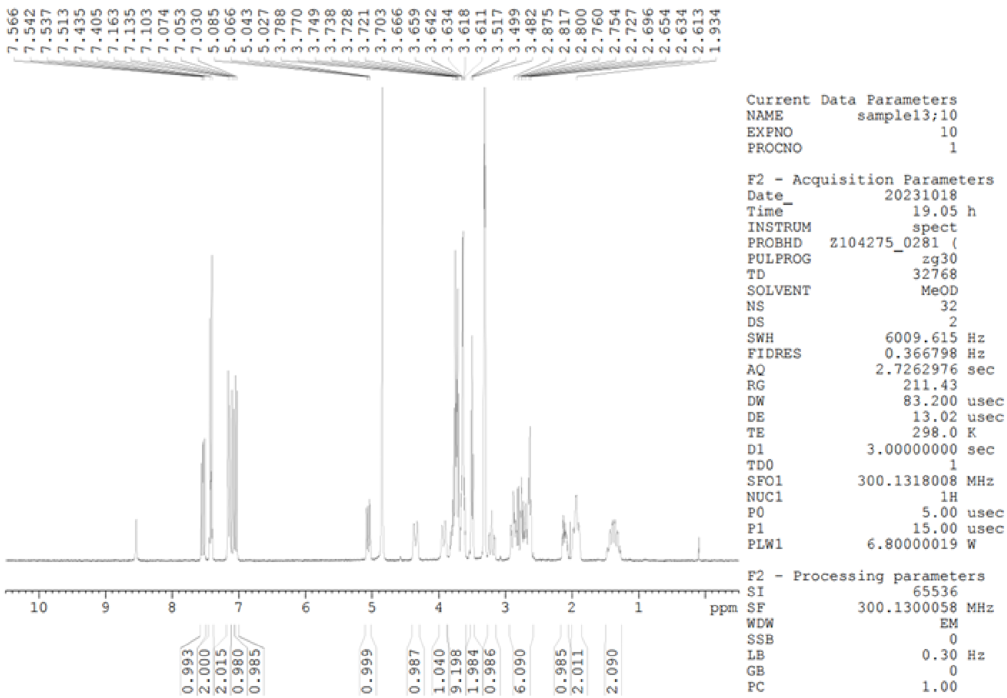


**$^{13}$C NMR (75 MHz, $CD_3OD$)**

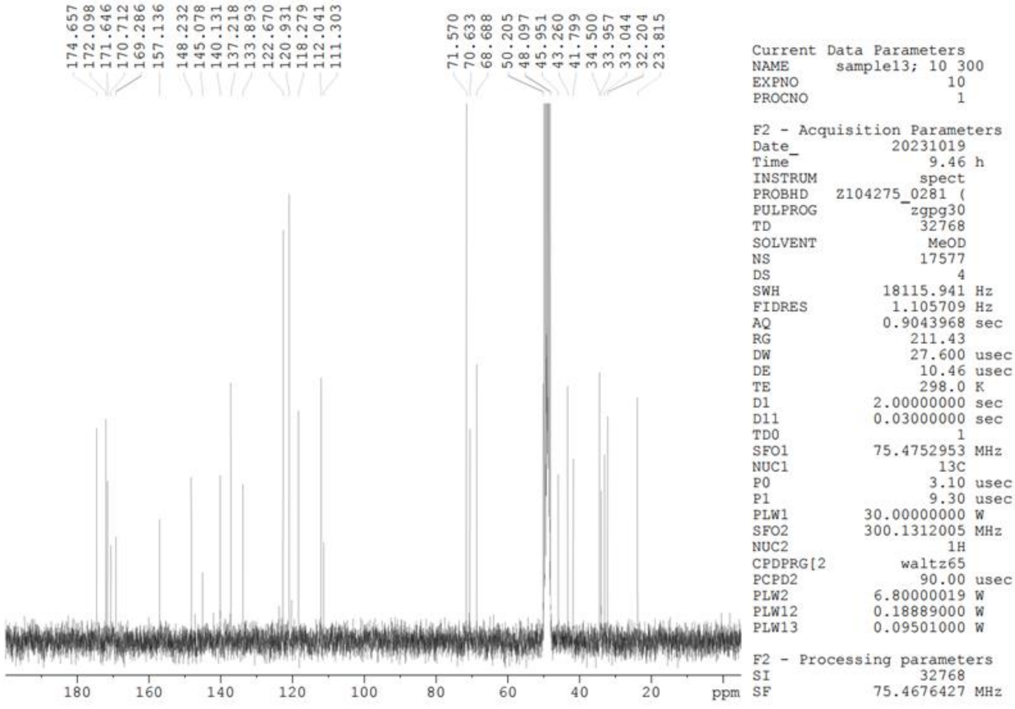

**Compound 10**

**[19]F NMR (565 MHz, $CD_3OD$)** TFA was used as an internal standard.

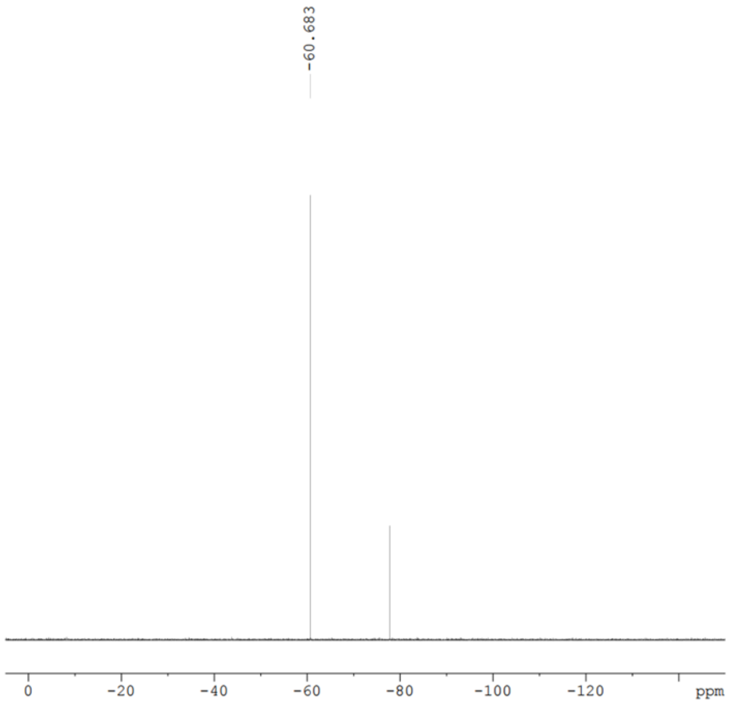


```
Current Data Parameters
NAME            sample13
EXPNO                 12
PROCNO                 1

F2 - Acquisition Parameters
Date_           20230616
Time               12.56 h
INSTRUM            spect
PROBHD   Z129649_0016 (
PULPROG      zgfhigqn.2
TD                131072
SOLVENT             MeOD
NS                    16
DS                     4
SWH           133928.578 Hz
FIDRES          2.043588 Hz
AQ             0.4893355 sec
RG                199.85
DW                 3.733 usec
DE                 18.00 usec
TE                 298.0 K
D1            1.00000000 sec
D11           0.03000000 sec
D12           0.00002000 sec
TD0                    1
SFO1         564.5170181 MHz
NUC1                 19F
P1                 10.90 usec
PLW1          3.90000010 W
SFO2         600.0124000 MHz
NUC2                  1H
CPDPRG[2         waltz16
PCPD2              70.00 usec
PLW2          7.19999981 W
PLW12         0.07851800 W

F2 - Processing parameters
SI                 65536
SF           564.5739447 MHz
```

**Compound 11**

**$^{1}$H NMR (300 MHz, $CD_3OD$)**

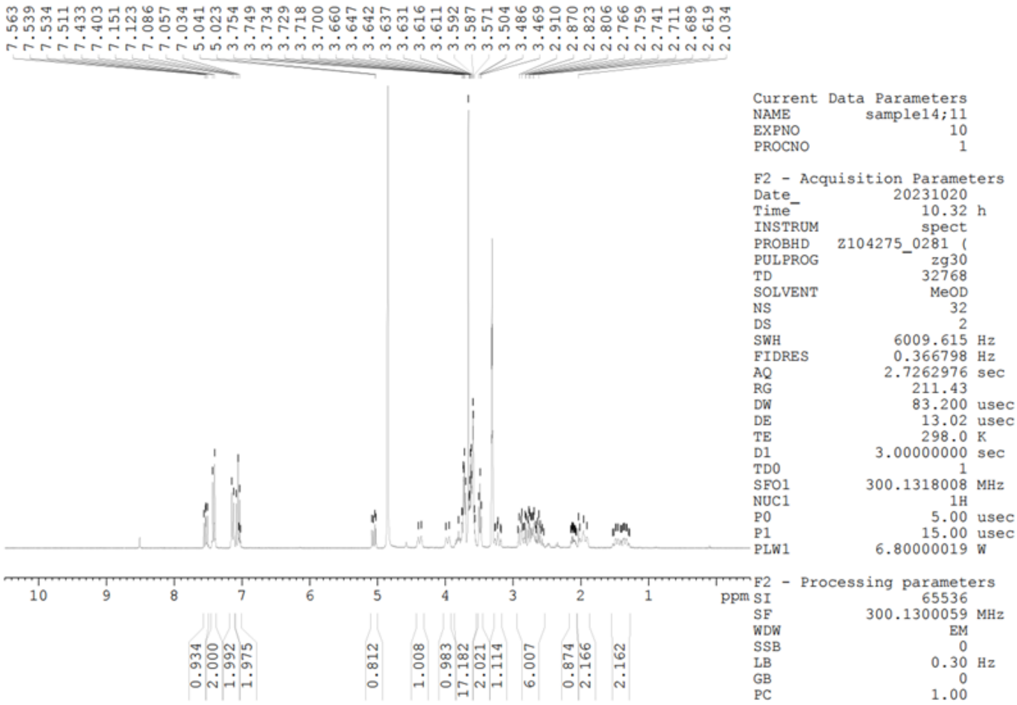


**$^{13}$C NMR (75 MHz, $CD_3OD$)**

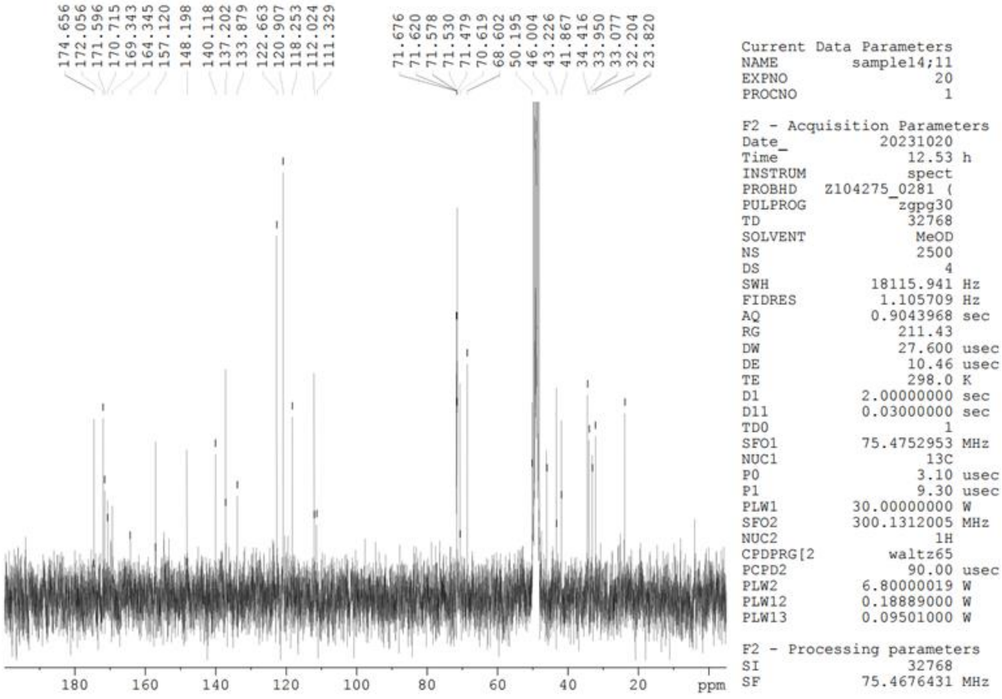

**Compound 11**

**$^{19}$F NMR (565 MHz, $CD_3OD$)** TFA was used as an internal standard.

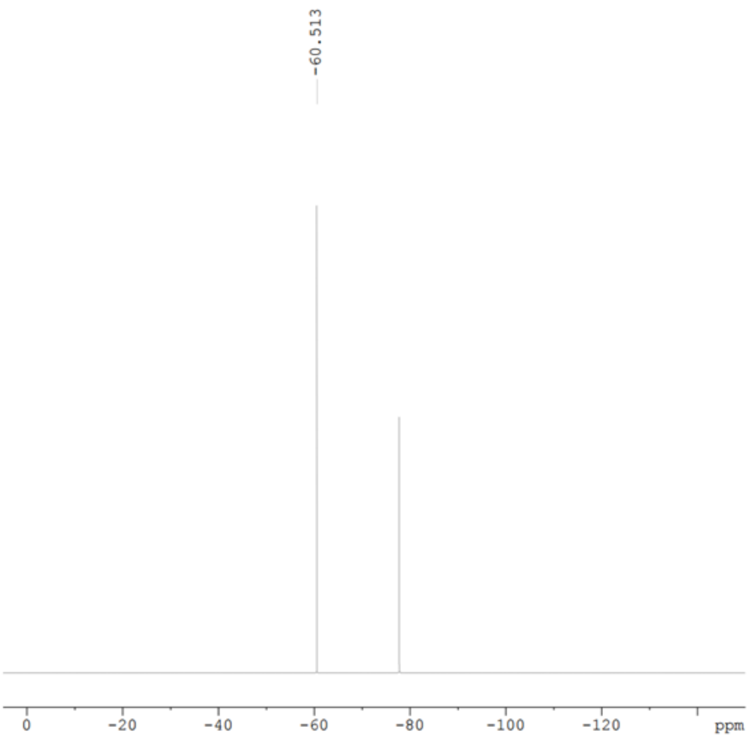


| Current Data Parameters | |
|---|---|
| NAME | sample14 |
| EXPNO | 12 |
| PROCNO | 1 |

| F2 - Acquisition Parameters | |
|---|---|
| Date_ | 20230616 |
| Time | 13.58 h |
| INSTRUM | spect |
| PROBHD | Z129649_0016 ( |
| PULPROG | zgflqn |
| TD | 131072 |
| SOLVENT | MeOD |
| NS | 16 |
| DS | 4 |
| SWH | 133928.578 Hz |
| FIDRES | 2.043588 Hz |
| AQ | 0.4893355 sec |
| RG | 199.85 |
| DW | 3.733 usec |
| DE | 18.00 usec |
| TE | 298.0 K |
| D1 | 1.00000000 sec |
| TD0 | 1 |
| SFO1 | 564.5170181 MHz |
| NUC1 | 19F |
| P1 | 10.90 usec |
| PLW1 | 3.90000010 W |

| F2 - Processing parameters | |
|---|---|
| SI | 65536 |
| SF | 564.5738527 MHz |
| WDW | EM |
| SSB | 0 |
| LB | 0.30 Hz |
| GB | 0 |
| PC | 1.00 |

**Compound 12**

**$^1$H NMR (300 MHz, $CD_3OD$)**

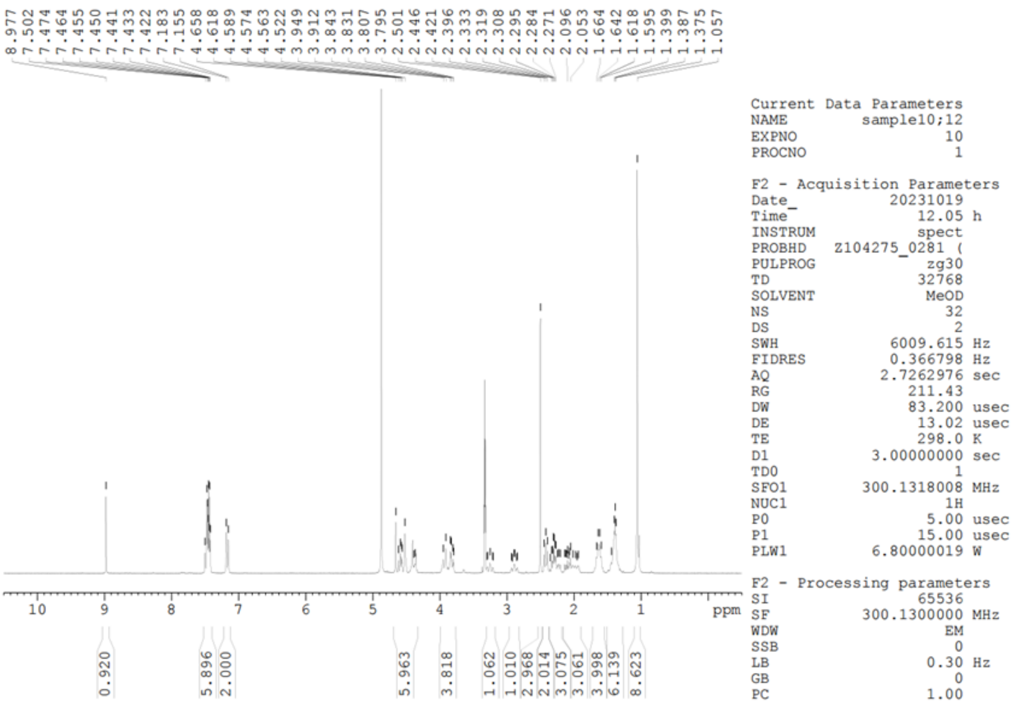


**$^{13}$C NMR (75 MHz, $CD_3OD$)**

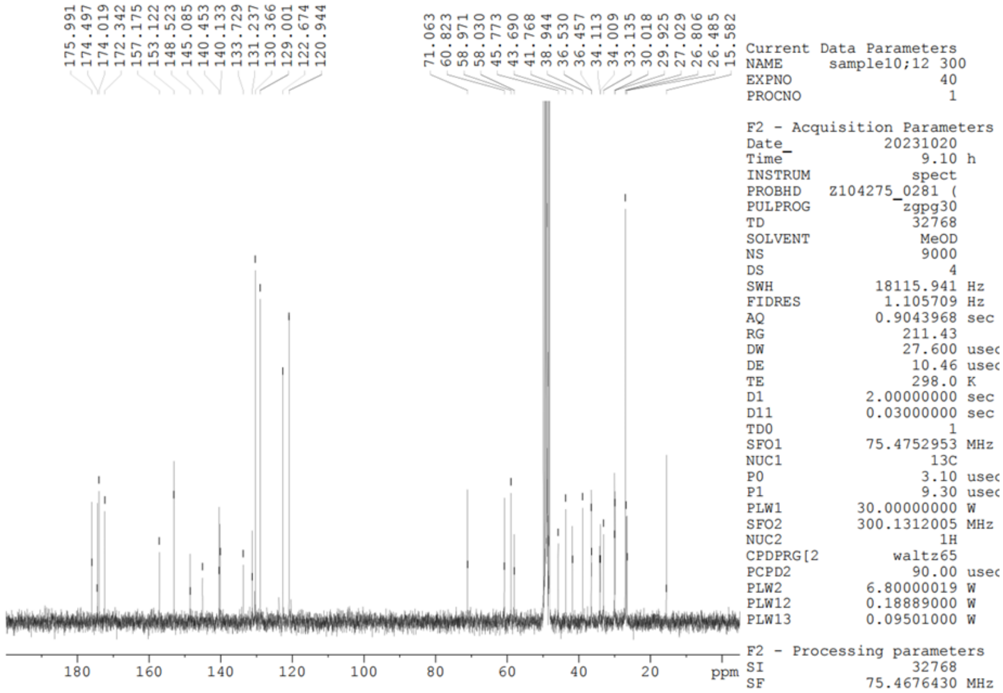

**Compound 12**

**$^{19}$F NMR (565 MHz, $CD_3OD$)** TFA was used as an internal standard.

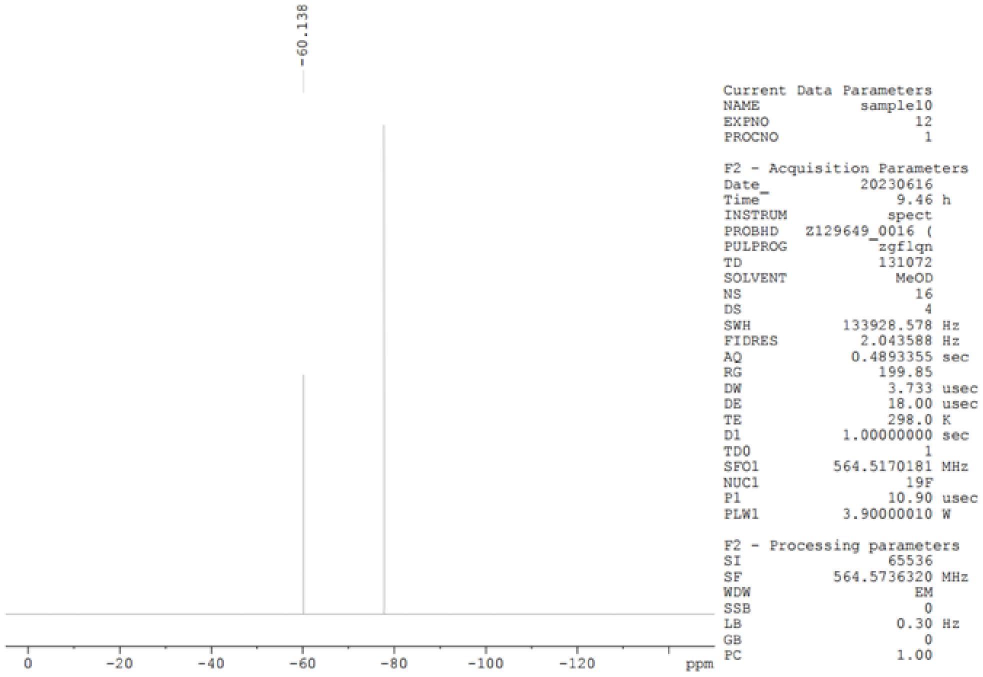

**Compound 13**

**$^1$H NMR (300 MHz, $CD_3OD$)**

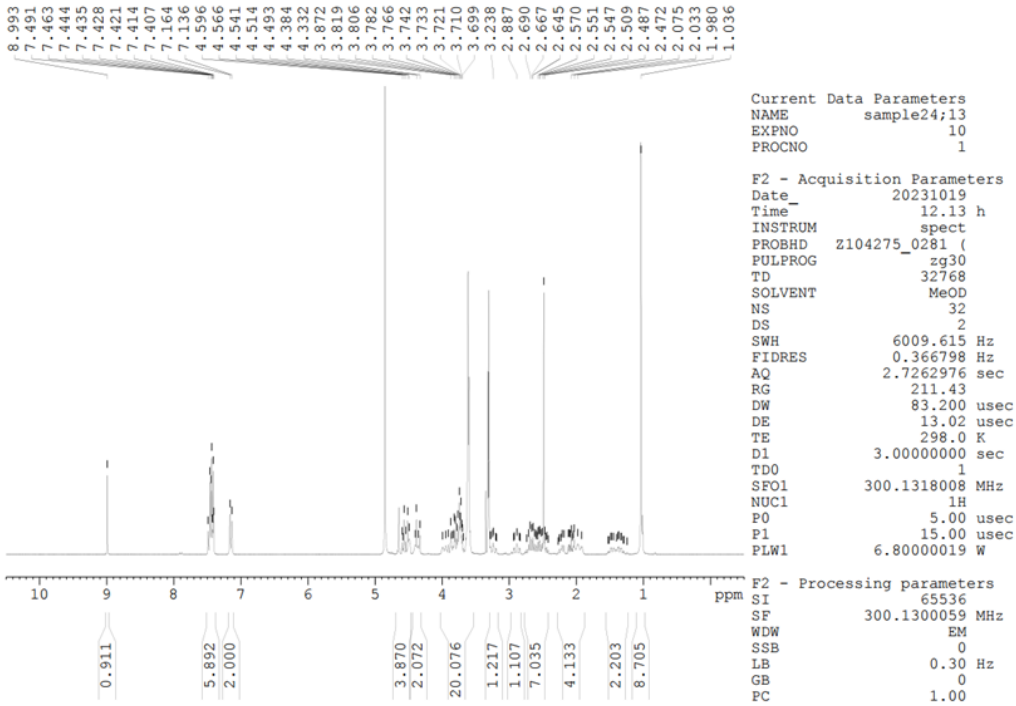


**$^{13}$C NMR (75 MHz, $CD_3OD$)**

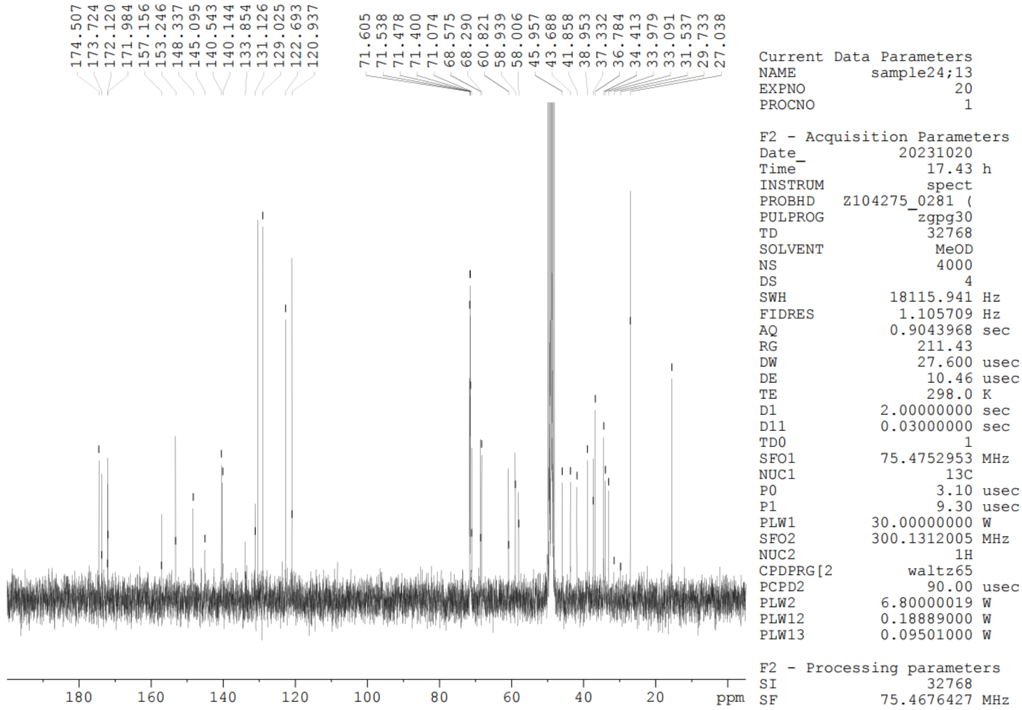

**Compound 13**

**$^{19}$F NMR (565 MHz, $CD_3OD$)** $CFCl_3$ was used as an internal standard.

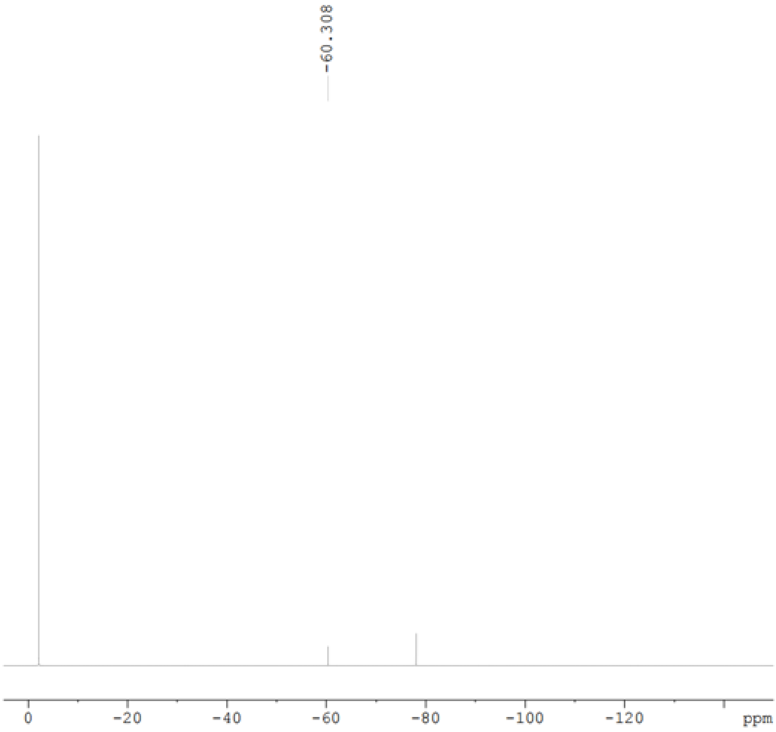


```
Current Data Parameters
NAME         sample24 F
EXPNO                11
PROCNO                1

F2 - Acquisition Parameters
Date_          20230621
Time               7.18 h
INSTRUM           spect
PROBHD   Z129649_0016 (
PULPROG          zgflqn
TD               131072
SOLVENT            MeOD
NS                   16
DS                    4
SWH          133928.578 Hz
FIDRES         2.043588 Hz
AQ            0.4893355 sec
RG               199.85
DW                3.733 usec
DE                18.00 usec
TE                298.0 K
D1           1.00000000 sec
TD0                   1
SFO1        564.5170181 MHz
NUC1                19F
P1                10.90 usec
PLW1         3.90000010 W

F2 - Processing parameters
SI                65536
SF          564.5737427 MHz
WDW                  EM
SSB                   0
LB                 0.30 Hz
GB                    0
PC                 1.00
```

**Compound 19**

**$^{1}$H NMR (300 MHz, $CD_3OD$)**

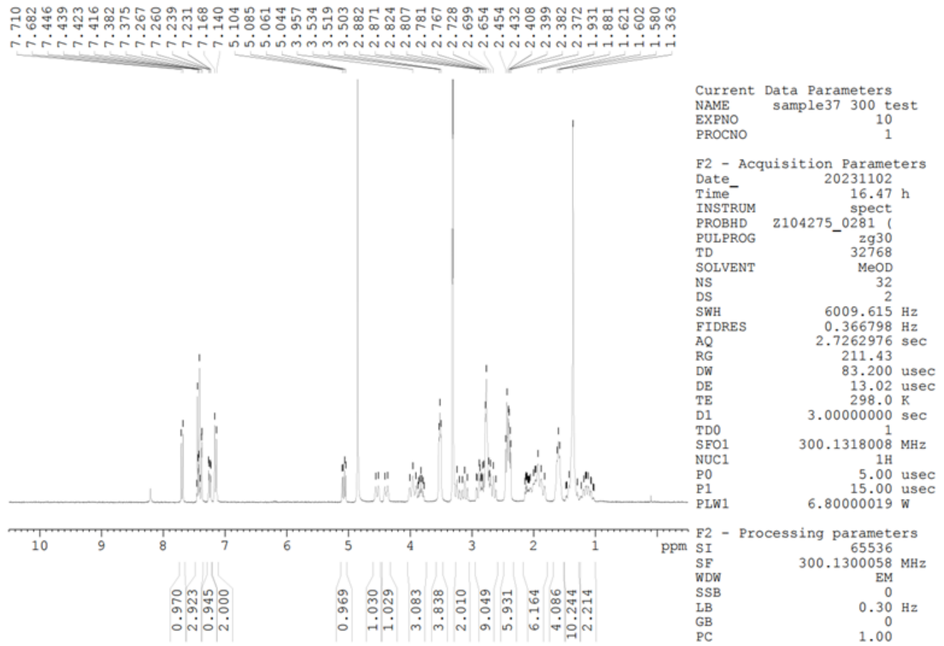


**$^{13}$C NMR (75 MHz, DMSO-$d_6$)**

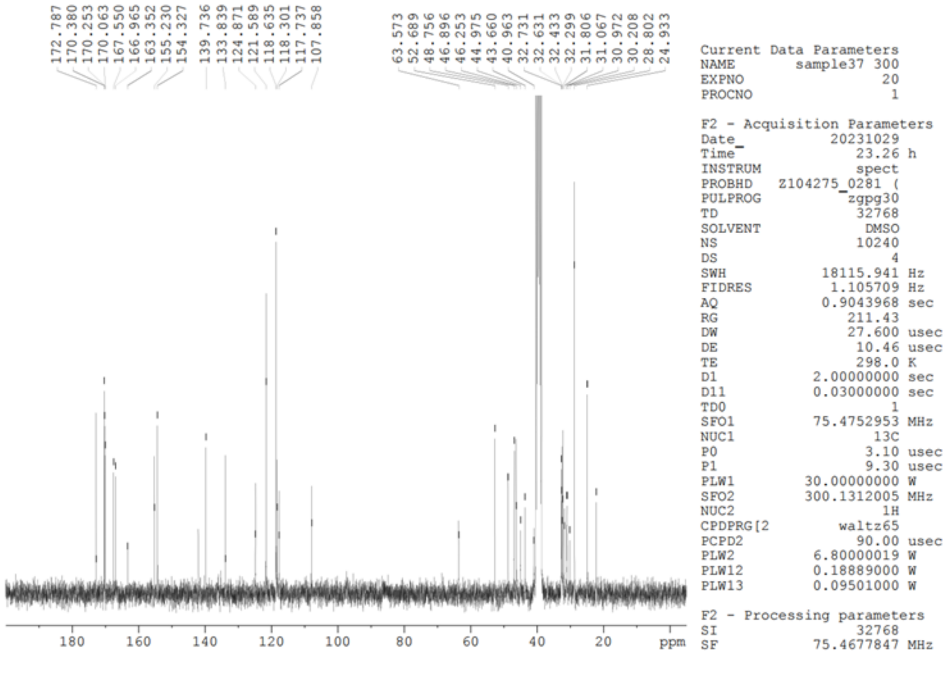

**Compound 19**

**$^{19}$F NMR (282 MHz, $CD_3OD$)** TFA was used as an internal standard.

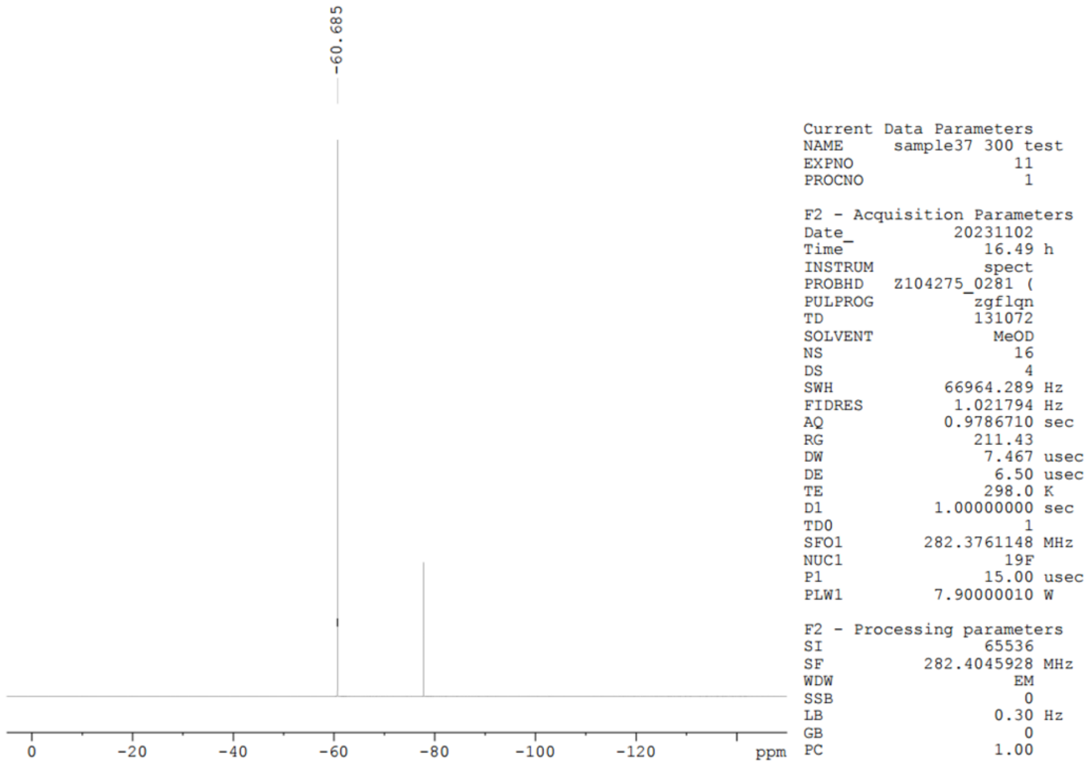

**Compound s1**

**$^{1}$H NMR (300 MHz, DMSO-$d_6$)**

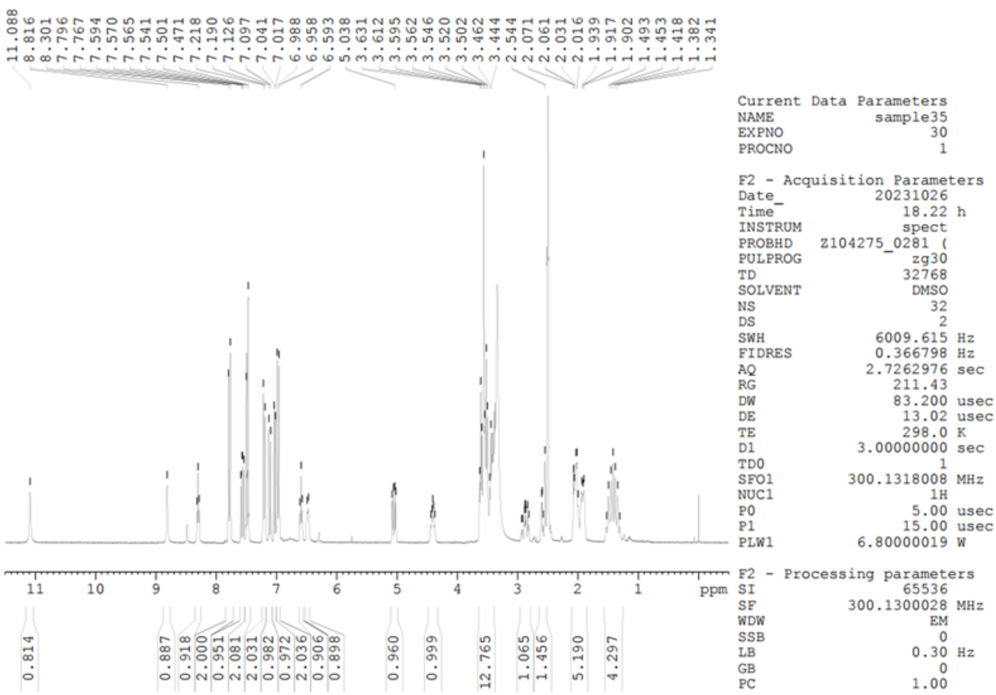


**$^{13}$C NMR (151 MHz, DMSO-$d_6$)**

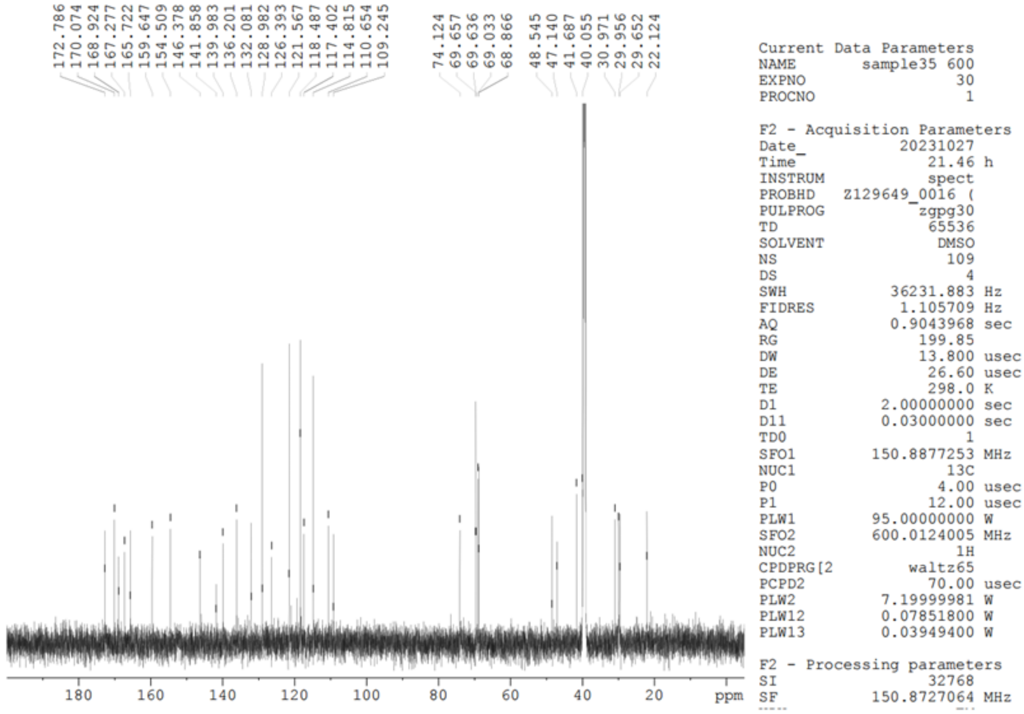

**Compound s1**

**$^{19}$F NMR (565 MHz, DMSO-$d_6$)** $CFCl_3$ was used as an internal standard.

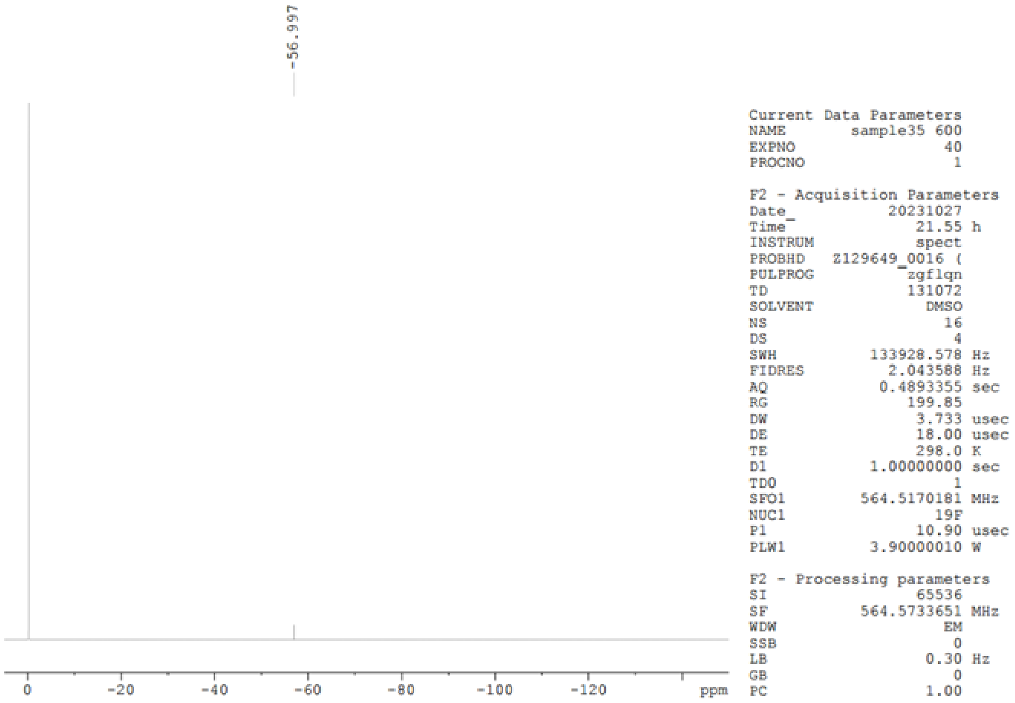

**Int 2**

**$^1H$ NMR (300 MHz, $CD_3OD$)**

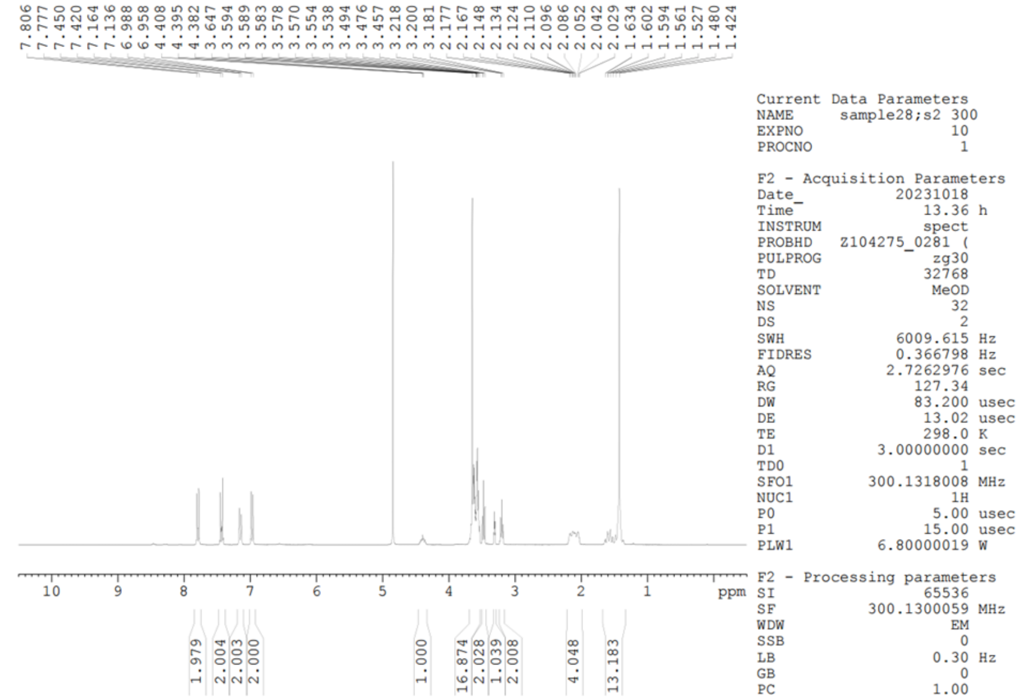

**$^{13}C$ NMR (75 MHz, $CD_3OD$)**

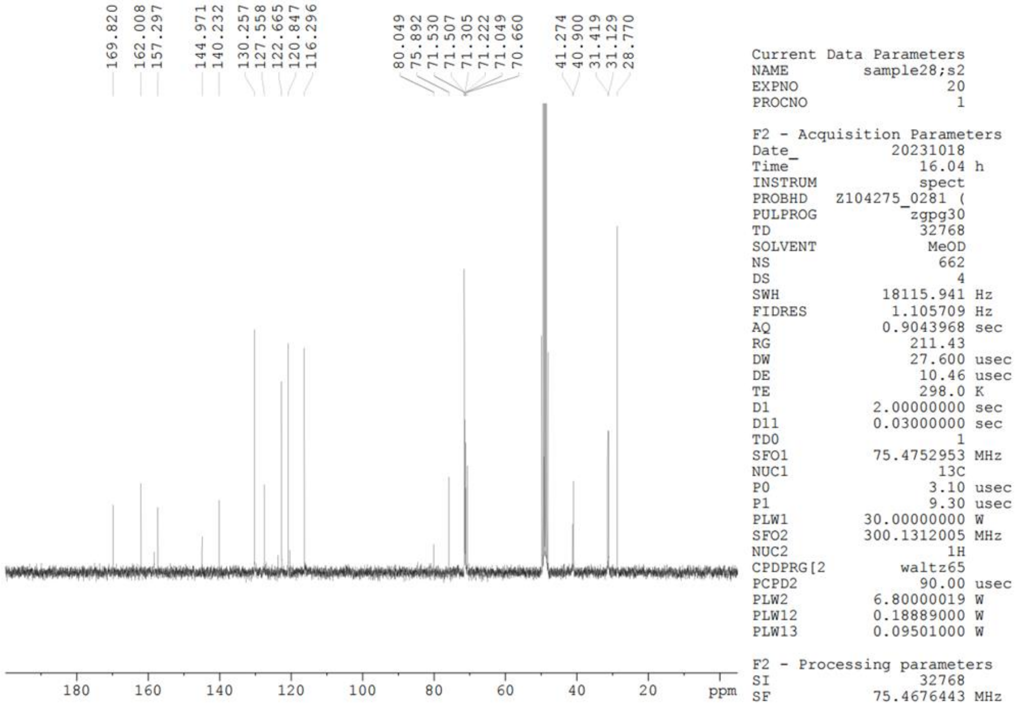

**Int 2**

**$^{19}$F NMR (565 MHz, $CD_3Cl$)** TFA was used as an internal standard.

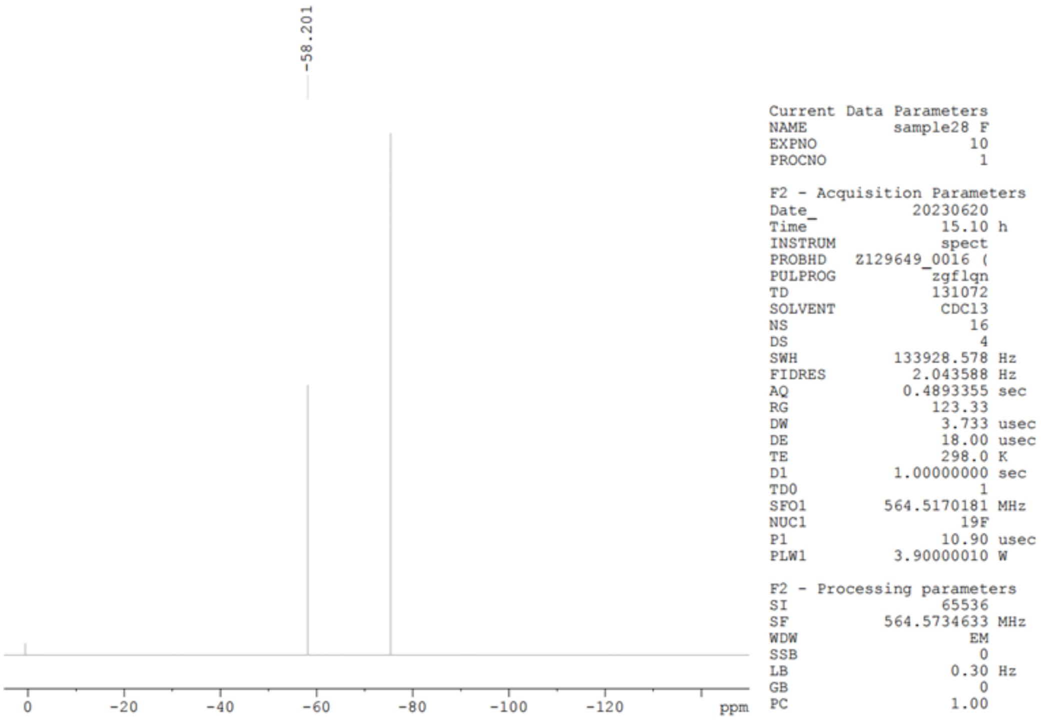

**Int 3**

**$^1$H NMR (300 MHz, $CD_3OD$)**

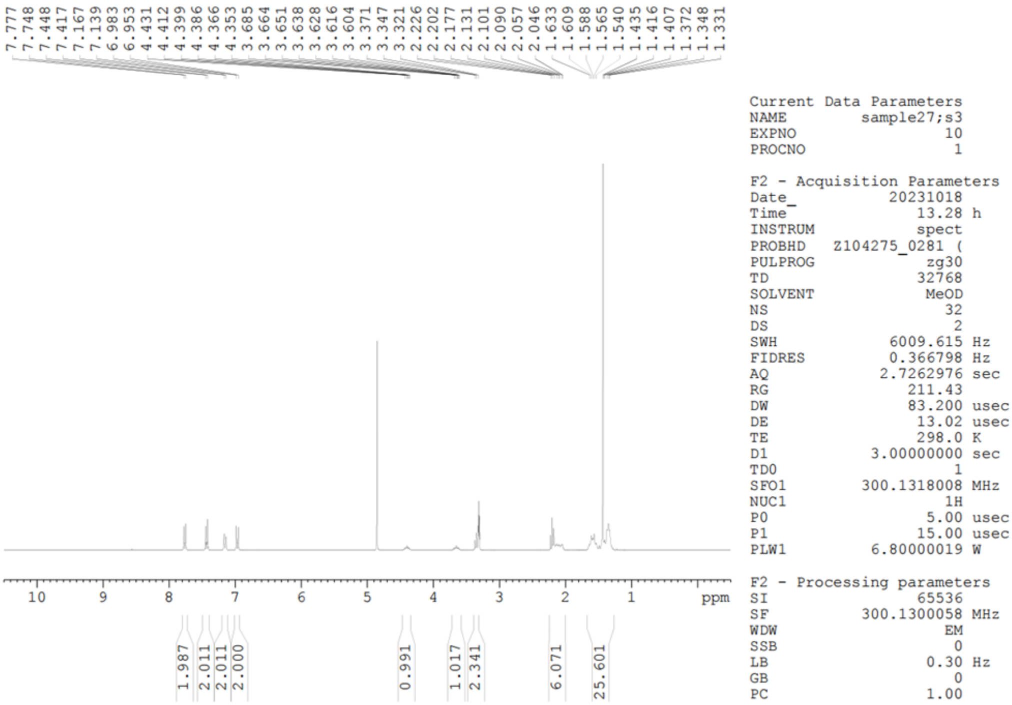


**$^{13}$C NMR (151 MHz, $CD_3OD$)**

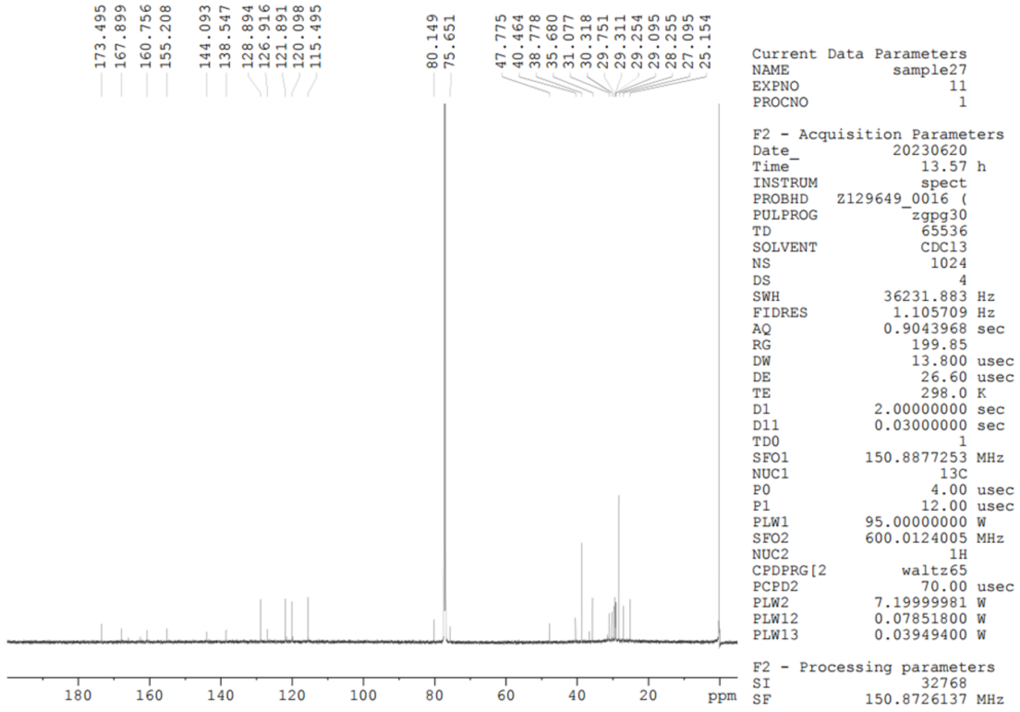

**Int 3**

**$^{19}$F NMR (282 MHz, $CD_3OD$)** $CFCl_3$ was used as an internal standard.

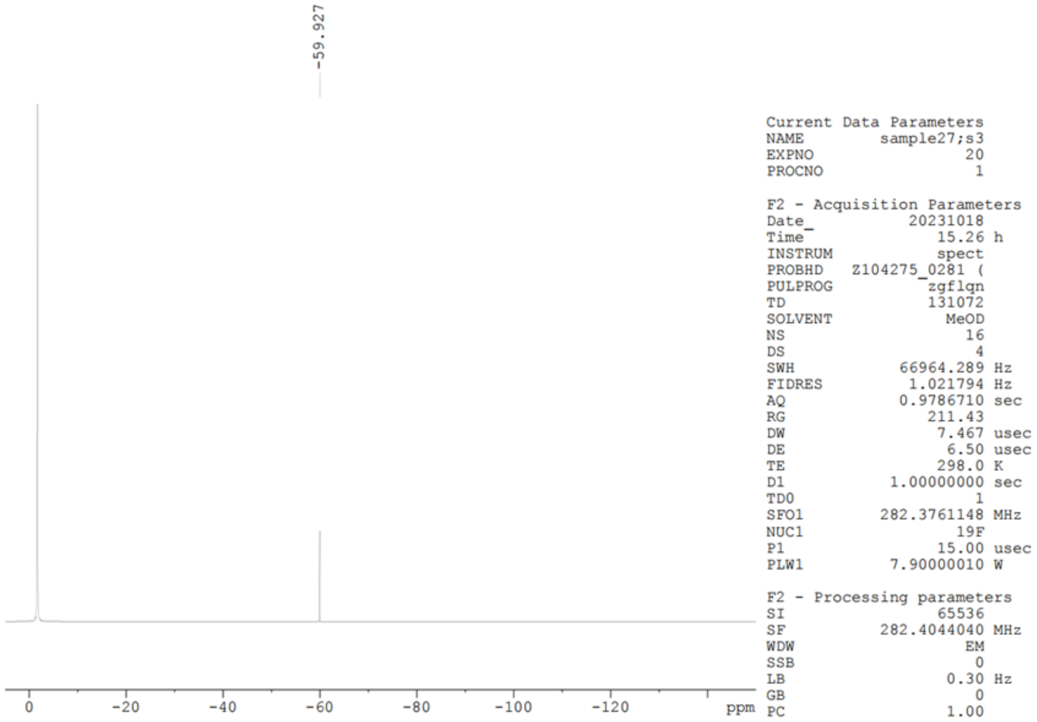

**Int 4**

**$^{1}$H NMR (600 MHz, $CD_3Cl$)**

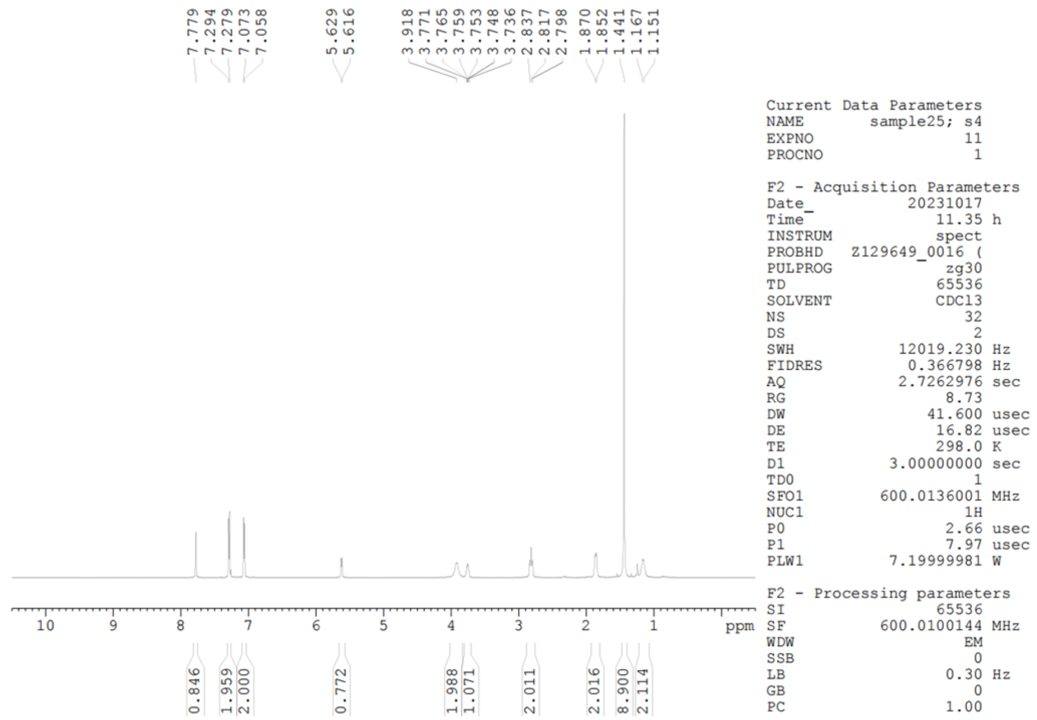


**$^{13}$C NMR (151 MHz, $CD_3Cl$)**

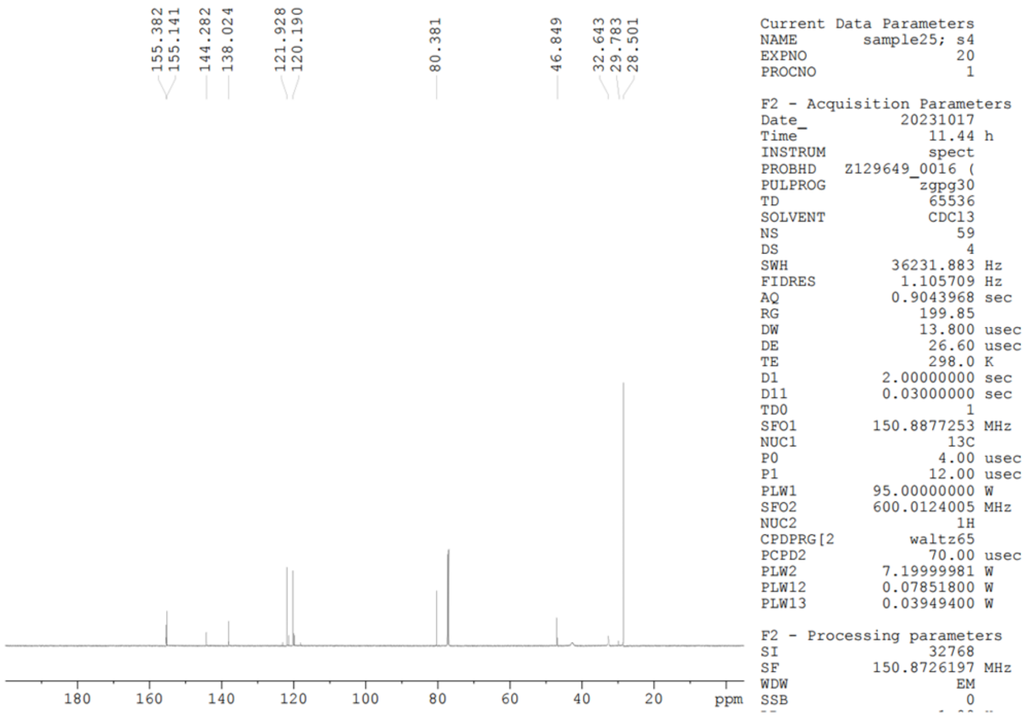

**Int 4**

**$^{19}$F NMR (565 MHz, $CD_3Cl$)** $CFCl_3$ was used as an internal standard.

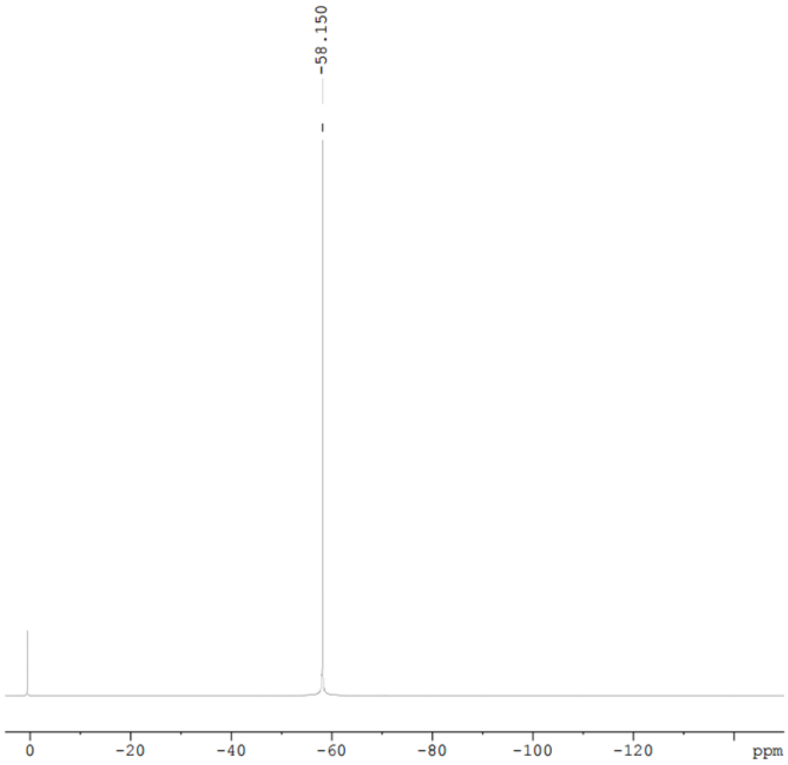


Current Data Parameters
NAME sample29 F
EXPNO 10
PROCNO 1

F2 - Acquisition Parameters
Date_ 20230620
Time 15.15 h
INSTRUM spect
PROBHD Z129649_0016 (
PULPROG zgflqn
TD 131072
SOLVENT CDCl3
NS 16
DS 4
SWH 133928.578 Hz
FIDRES 2.043588 Hz
AQ 0.4893355 sec
RG 31.64
DW 3.733 usec
DE 18.00 usec
TE 298.0 K
D1 1.00000000 sec
TD0 1
SFO1 564.5170181 MHz
NUC1 19F
P1 10.90 usec
PLW1 3.90000010 W

F2 - Processing parameters
SI 65536
SF 564.5734754 MHz
WDW EM
SSB 0
LB 0.30 Hz
GB 0
PC 1.00

**Int 5**

**$^1$H NMR (300 MHz, $CD_3Cl$)**

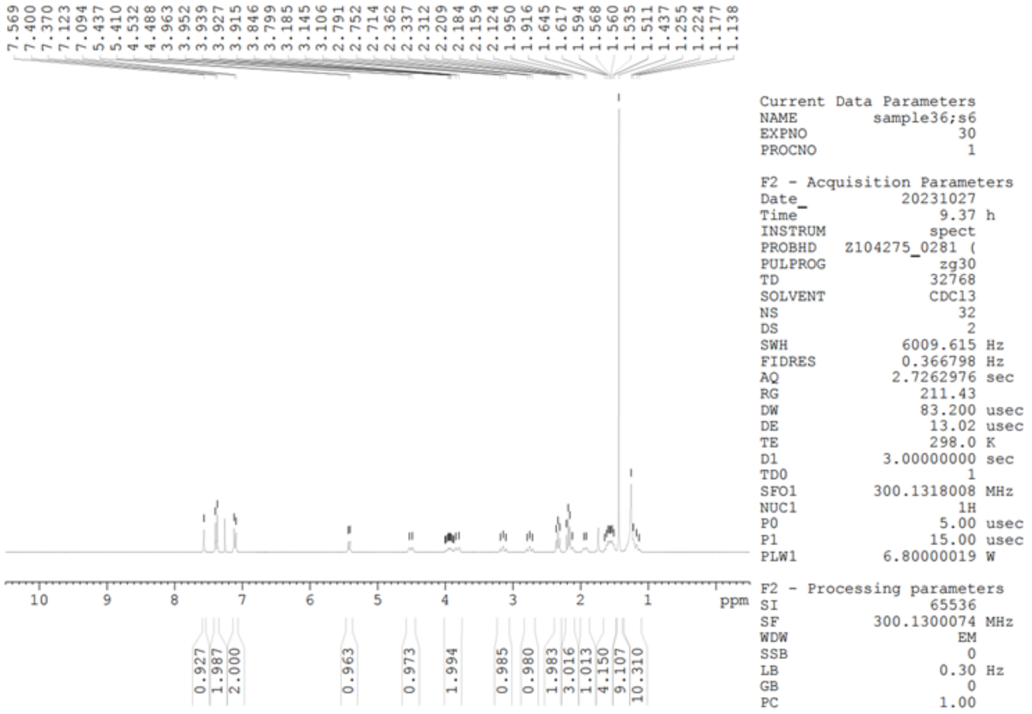

**$^{13}$C NMR (75 MHz, $CD_3Cl$)**

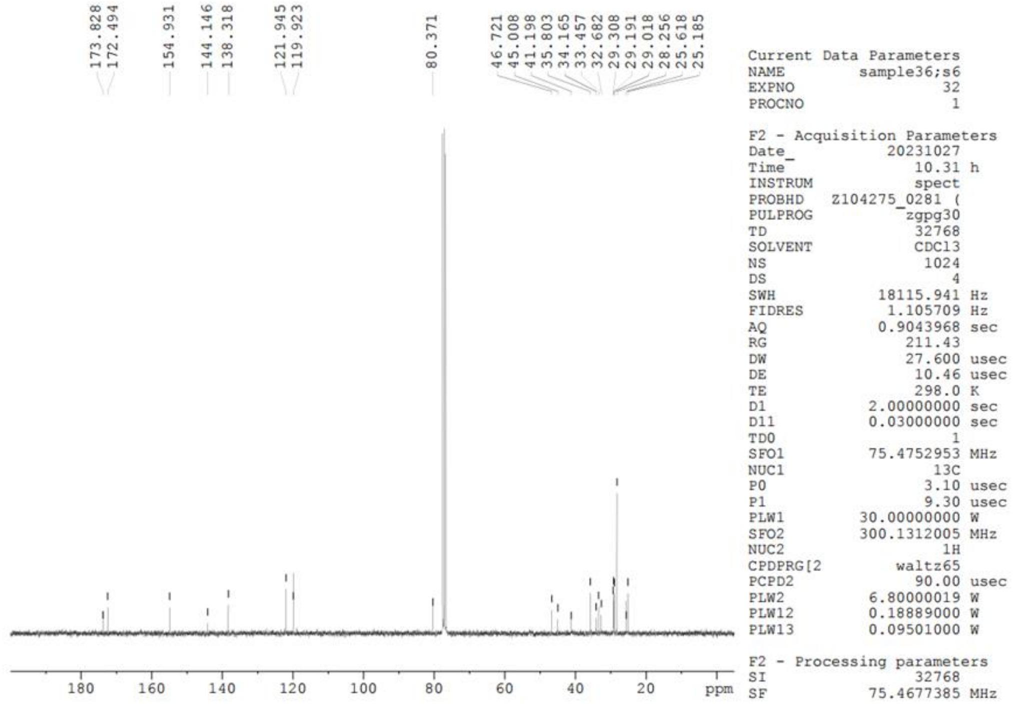

**Int 5**

**[19]F NMR (282 MHz, CD$_3$Cl)** CFCl$_3$ was used as an internal standard.

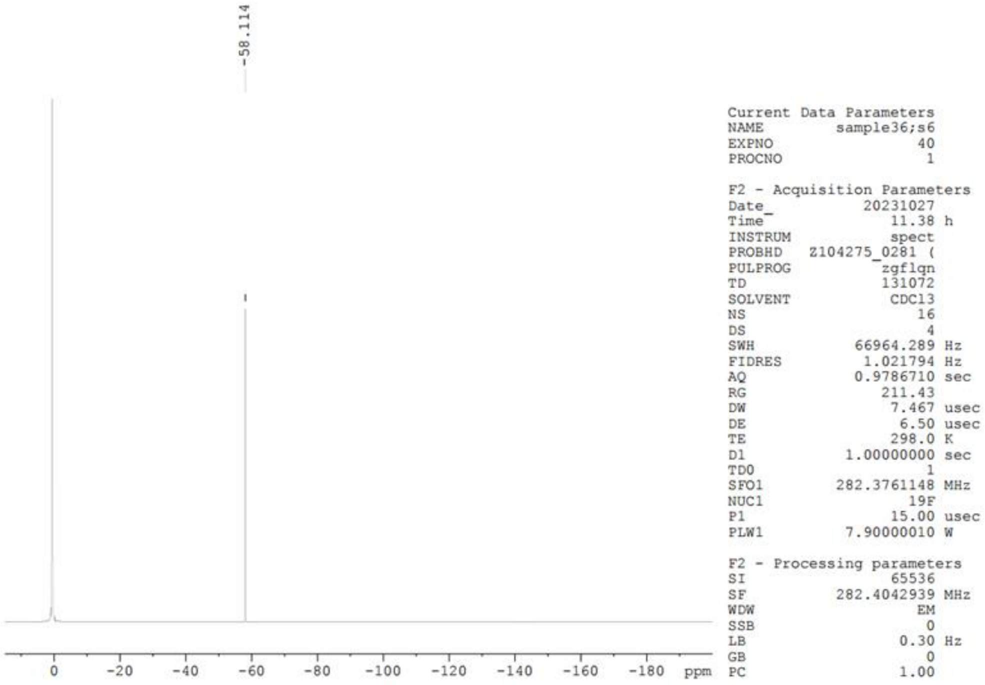

**HPLC traces**

Compound **8** (monitored at 254 nm)

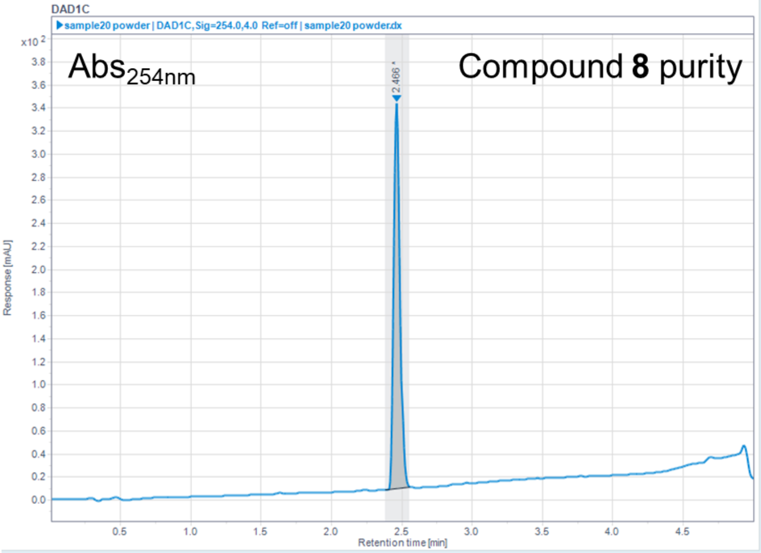


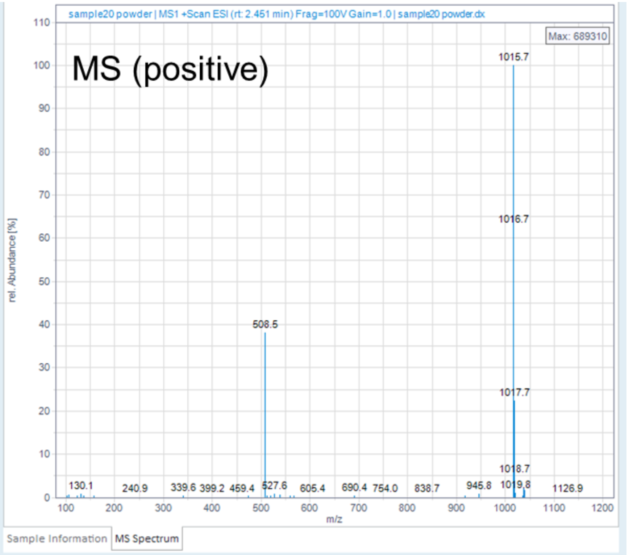


Injection Results

Peaks | Summary

| # | Name | Signal description | RT (min) | Area (mAU·s) | Area% | Height (mAU) | Height% | Amount | Concentration | Start time (min) | End time (min) |
|---|---|---|---|---|---|---|---|---|---|---|---|
| 1 | | DAD1C,Sig=254.0,4.0 | 2.466 | 1009.545 | 100.000 | 333.978 | 100.00 | | | 2.381 | 2.559 |

Compound **17** (monitored at 254 nm)

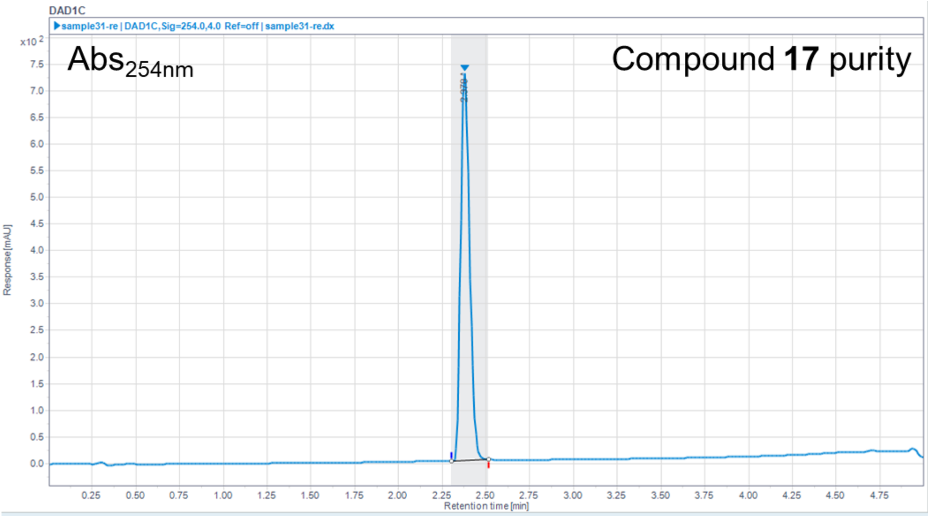


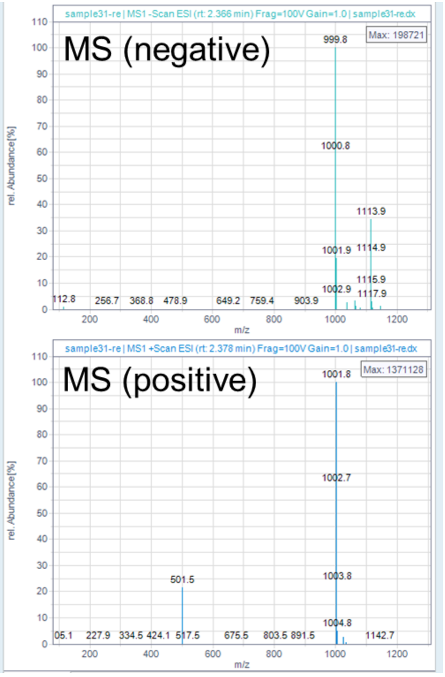


Injection Results

Peaks | Summary

| # | Name | Signal description | RT (min) | Area (mAU·s) | Area% | Height (mAU) | Height% | Amount | Concentration | Start time (min) | End time (min) |
|---|---|---|---|---|---|---|---|---|---|---|---|
| 1 | | DAD1C,Sig=254.0,4.0 | 2.378 | 2576.041 | 100.000 | 729.752 | 100.00 | | | 2.303 | 2.514 |

Compound **19** (monitored at 254 nm)

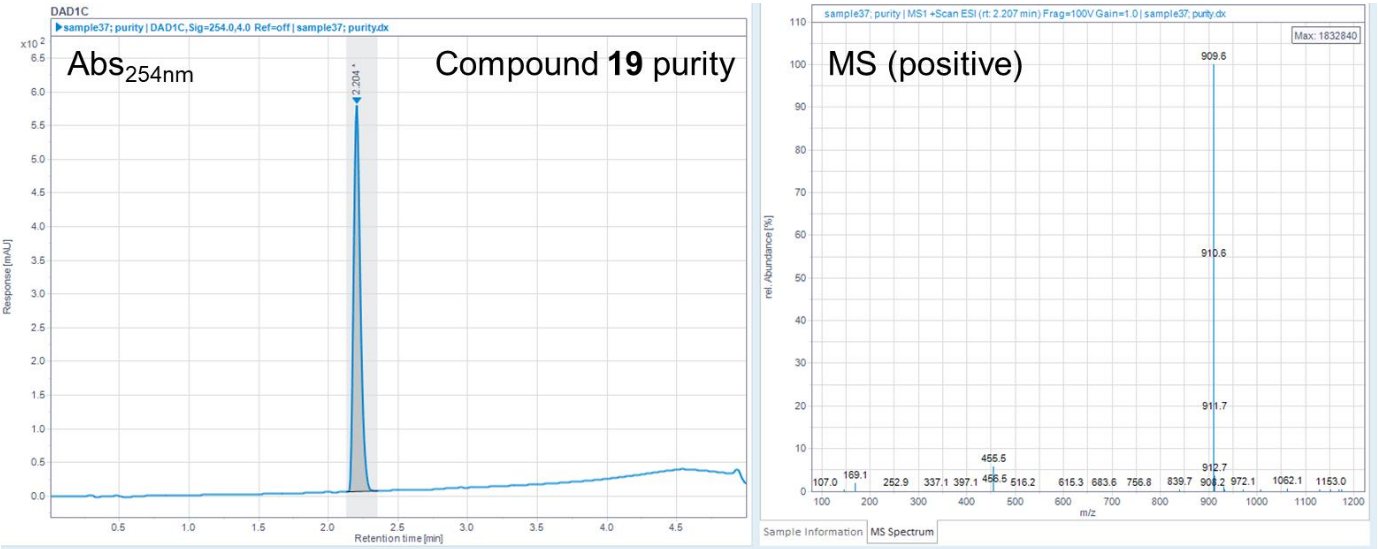


Injection Results

Peaks | Summary

| # | Name | Signal description | RT (min) | Area (mAU·s) | Area% | Height (mAU) | Height% | Amount | Concentration | Start time (min) | End time (min) |
|---|---|---|---|---|---|---|---|---|---|---|---|
| 1 | | DAD1C,Sig=254.0,4.0 | 2.204 | 1948.610 | 100.000 | 573.253 | 100.00 | | | 2.127 | 2.352 |

### Reference


(1) Schumacher, H.; Smith, R. L.; Williams, R. T. The Metabolism of Thalidomide: The Spontaneous Hydrolysis of Thalidomide in Solution. *Br. J. Pharmacol. Chemother.* **1965**, *25* (2), 324–337. https://doi.org/10.1111/j.1476-5381.1965.tb02053.x.

(2) Fulmer, G. R.; Miller, A. J. M.; Sherden, N. H.; Gottlieb, H. E.; Nudelman, A.; Stoltz, B. M.; Bercaw, J. E.; Goldberg, K. I. NMR Chemical Shifts of Trace Impurities: Common Laboratory Solvents, Organics, and Gases in Deuterated Solvents Relevant to the Organometallic Chemist. *Organometallics* **2010**, *29* (9), 2176–2179. https://doi.org/10.1021/om100106e.

(3) Rosenau, C. P.; Jelier, B. J.; Gossert, A. D.; Togni, A. Exposing the Origins of Irreproducibility in Fluorine NMR Spectroscopy. *Angew. Chem. Int. Ed Engl.* **2018**, *57* (30), 9528–9533. https://doi.org/10.1002/anie.201802620.

(4) Wang, Y.; Morisseau, C.; Takamura, A.; Wan, D.; Li, D.; Sidoli, S.; Yang, J.; Wolan, D. W.; Hammock, B. D.; Kitamura, S. PROTAC-Mediated Selective Degradation of Cytosolic Soluble Epoxide Hydrolase Enhances ER Stress Reduction. *ACS Chem. Biol.* **2023**, *18* (4), 884–896. https://doi.org/10.1021/acschembio.3c00017.

(5) Hwang, S. H.; Tsai, H.-J.; Liu, J.-Y.; Morisseau, C.; Hammock, B. D. Orally Bioavailable Potent Soluble Epoxide Hydrolase Inhibitors. *J. Med. Chem.* **2007**, *50* (16), 3825–3840. https://doi.org/10.1021/jm070270t.

(6) Peyman, M.; Barroso, E.; Turcu, A. L.; Estrany, F.; Smith, D.; Jurado-Aguilar, J.; Rada, P.; Morisseau, C.; Hammock, B. D.; Valverde, Á. M.; Palomer, X.; Galdeano, C.; Vázquez, S.; Vázquez-Carrera, M. Soluble Epoxide Hydrolase-Targeting PROTAC Activates AMPK and Inhibits Endoplasmic Reticulum Stress. *Biomed. Pharmacother.* **2023**, *168*, 115667. https://doi.org/10.1016/j.biopha.2023.115667.

(7) Lee, K. S. S.; Liu, J.-Y.; Wagner, K. M.; Pakhomova, S.; Dong, H.; Morisseau, C.; Fu, S. H.; Yang, J.; Wang, P.; Ulu, A.; Mate, C. A.; Nguyen, L. V.; Hwang, S. H.; Edin, M. L.; Mara, A. A.; Wulff, H.; Newcomer, M. E.; Zeldin, D. C.; Hammock, B. D. Optimized Inhibitors of Soluble Epoxide Hydrolase Improve *in Vitro* Target Residence Time and *in Vivo* Efficacy. *J. Med. Chem.* **2014**, *57* (16), 7016–7030. https://doi.org/10.1021/jm500694p.

(8) Jones, P. D.; Wolf, N. M.; Morisseau, C.; Whetstone, P.; Hock, B.; Hammock, B. D. Fluorescent Substrates for Soluble Epoxide Hydrolase and Application to Inhibition Studies. *Anal. Biochem.* **2005**, *343* (1), 66–75. https://doi.org/10.1016/j.ab.2005.03.041.

(9) Li, D.; Cui, Y.; Morisseau, C.; Gee, S. J.; Bever, C. S.; Liu, X.; Wu, J.; Hammock, B. D.; Ying, Y. Nanobody Based Immunoassay for Human Soluble Epoxide Hydrolase Detection Using Polymeric Horseradish Peroxidase (PolyHRP) for Signal Enhancement: The Rediscovery of PolyHRP? *Anal. Chem.* **2017**, *89* (11), 6248–6256. https://doi.org/10.1021/acs.analchem.7b01247.

(10) Li, D.; Morisseau, C.; McReynolds, C. B.; Duflot, T.; Bellien, J.; Nagra, R. M.; Taha, A. Y.; Hammock, B. D. Development of Improved Double-Nanobody Sandwich ELISAs for Human Soluble Epoxide Hydrolase Detection in Peripheral Blood Mononuclear Cells of Diabetic Patients and the Prefrontal Cortex of Multiple Sclerosis Patients. *Anal. Chem.* **2020**, *92* (10), 7334–7342. https://doi.org/10.1021/acs.analchem.0c01115.

(11) Gill, S. S.; Hammock, B. D. Distribution and Properties of a Mammalian Soluble Epoxide Hydrase. *Biochem. Pharmacol.* **1980**, *29* (3), 389–395. https://doi.org/10.1016/0006-2952(80)90518-3.

(12) Aguilan, J. T.; Kulej, K.; Sidoli, S. Guide for Protein Fold Change and P-Value Calculation for Non-Experts in Proteomics. *Mol. Omics* **2020**, *16* (6), 573–582. https://doi.org/10.1039/D0MO00087F.
