## Supplementary material for "*In vivo*–Active Soluble Epoxide Hydrolase–targeting PROTACs with Improved Potency and Stability": Original data of immunoblotting

**Figure 3**

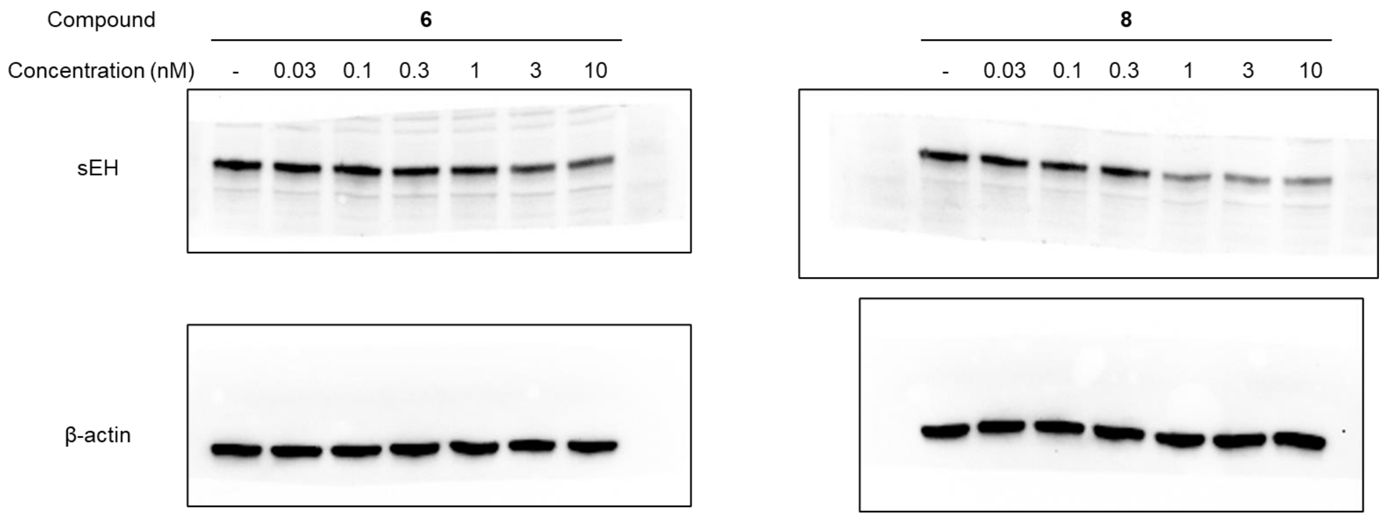


**Figure 3A**

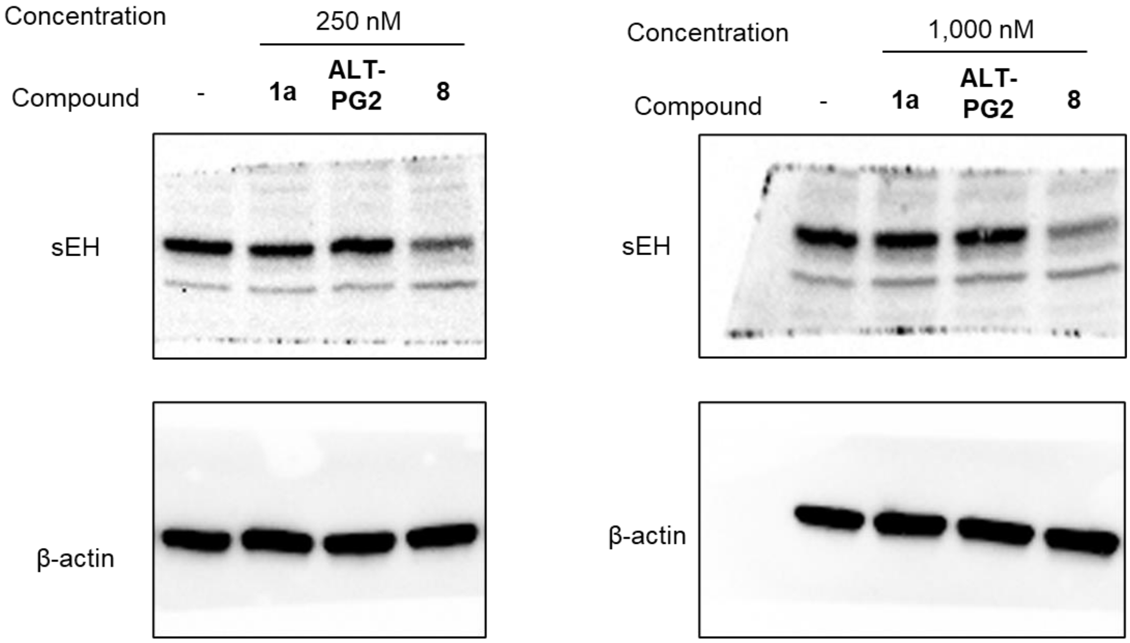


**Figure 3B**

Figure 4

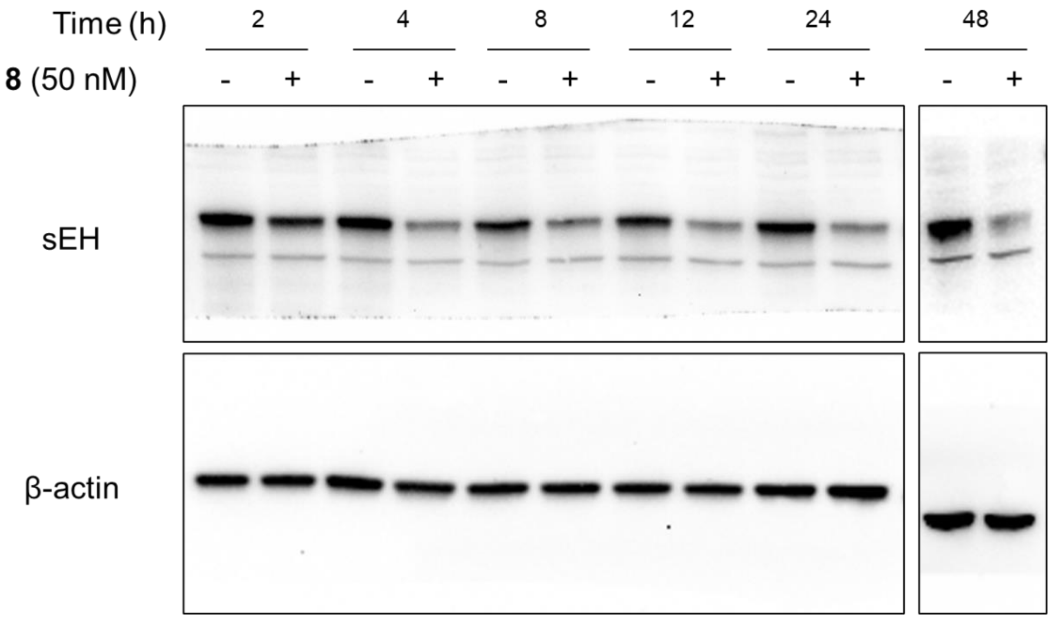


Figure 4A

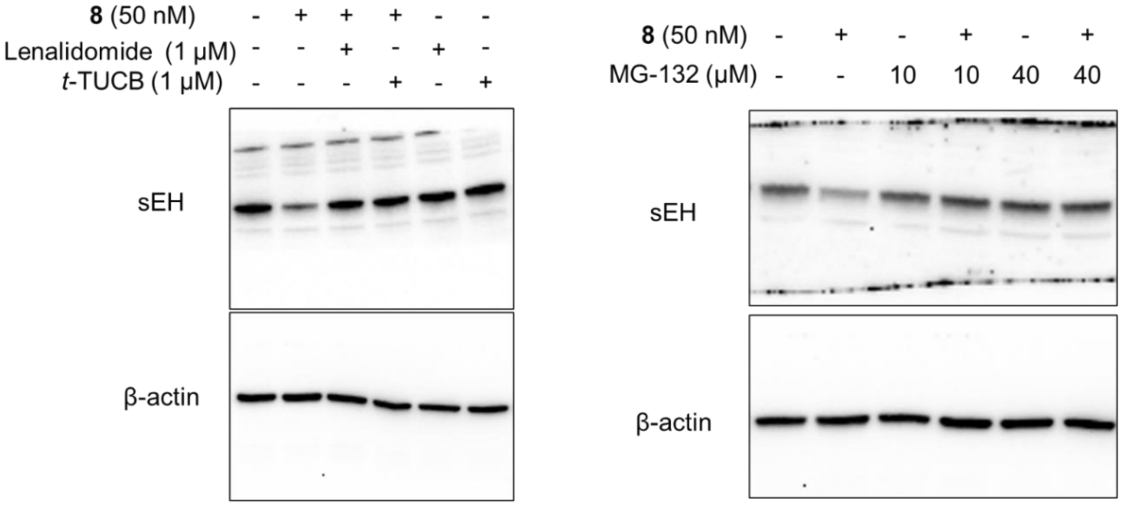


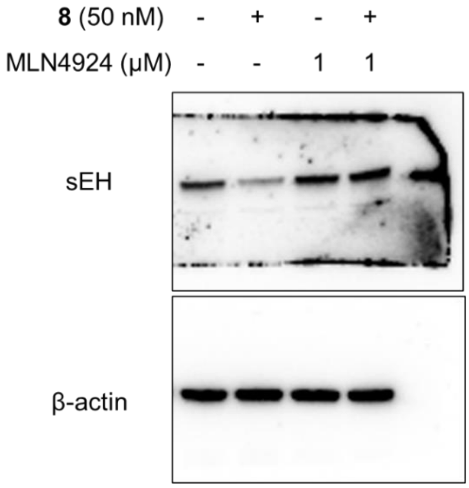


Figure 4B

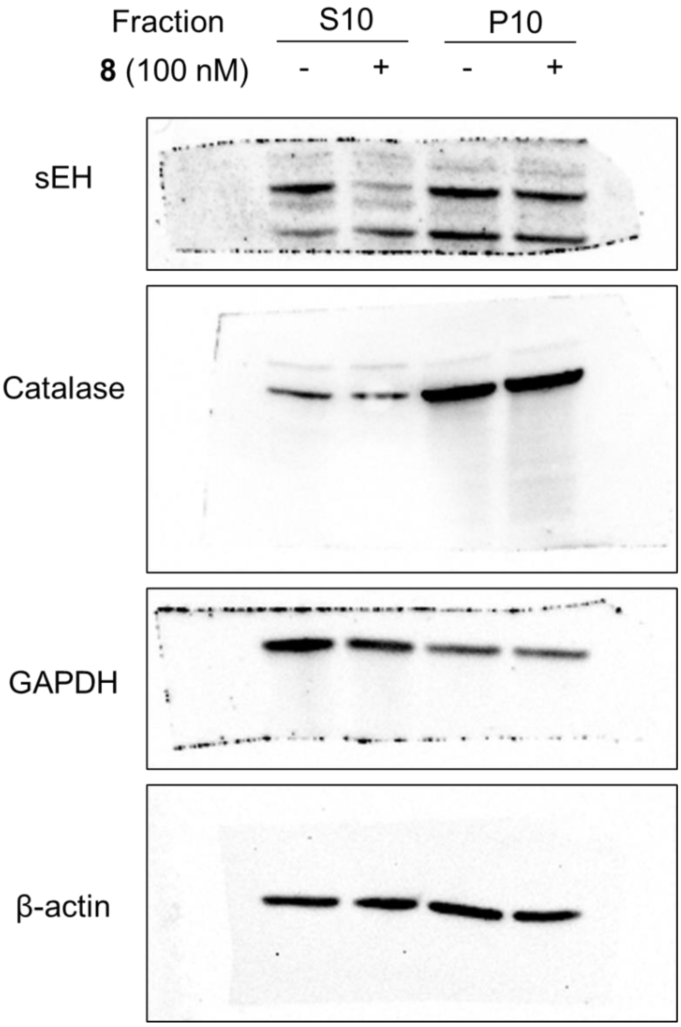


**Figure 4C**

**Figure 5 and Figure S9**

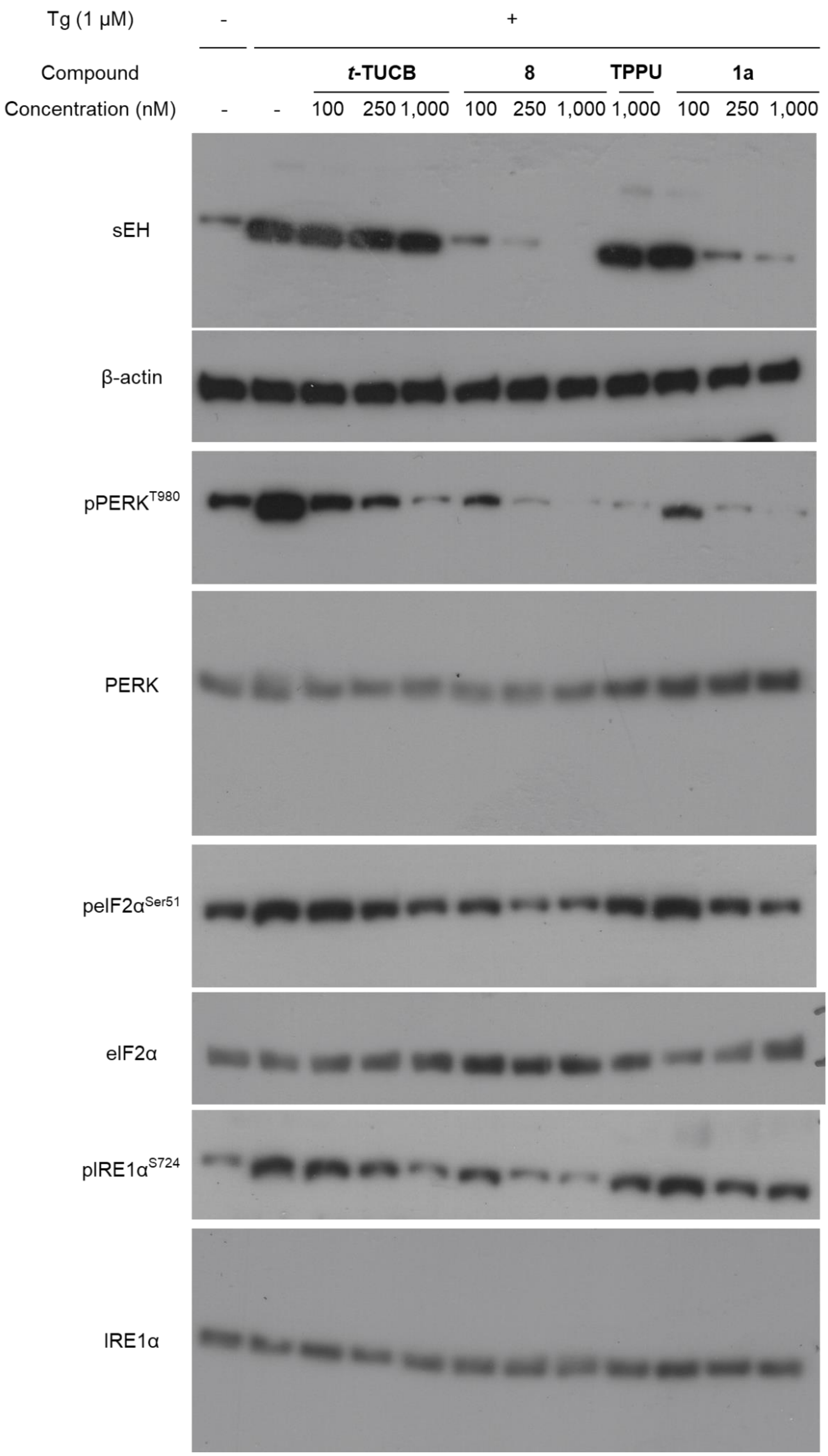

**Figure 6**

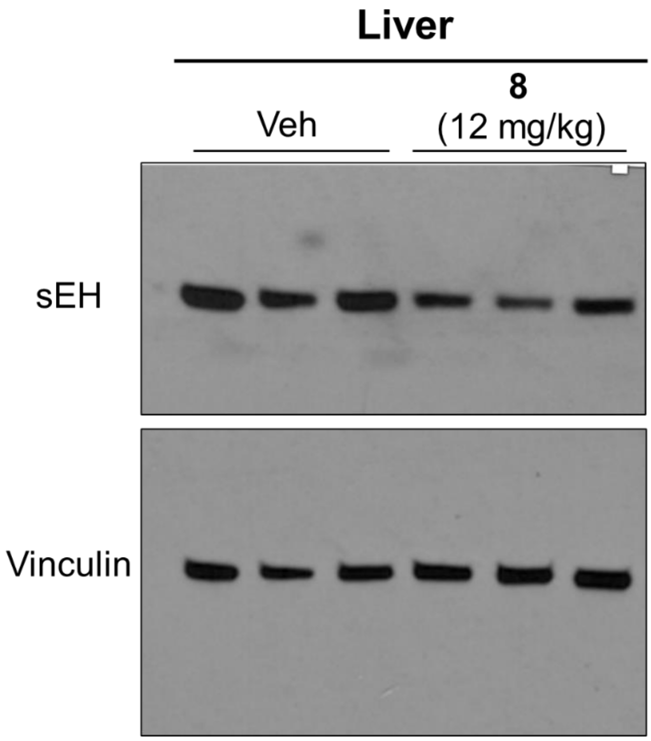


**Figure 6B**

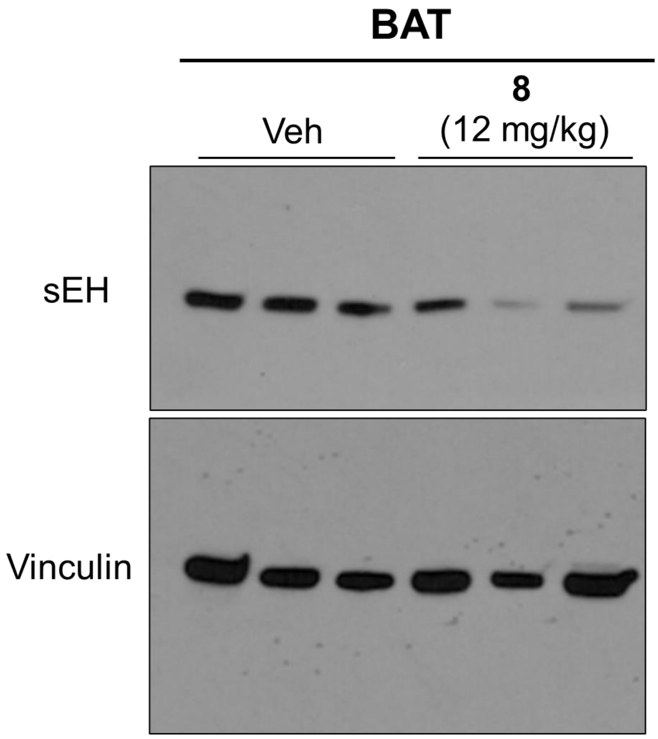


**Figure 6C**

**Figure S2**

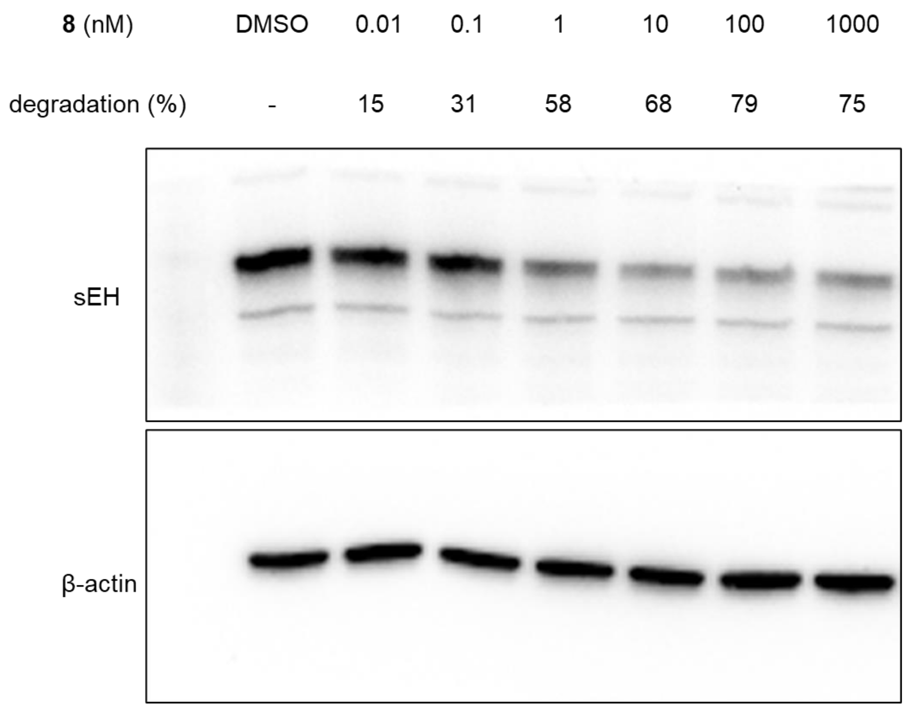

**8** (nM)
DMSO
0.01
0.1
1
10
100
1000
degradation (%)
-
15
31
58
68
79
75
sEH
β-actin

**Figure S3**

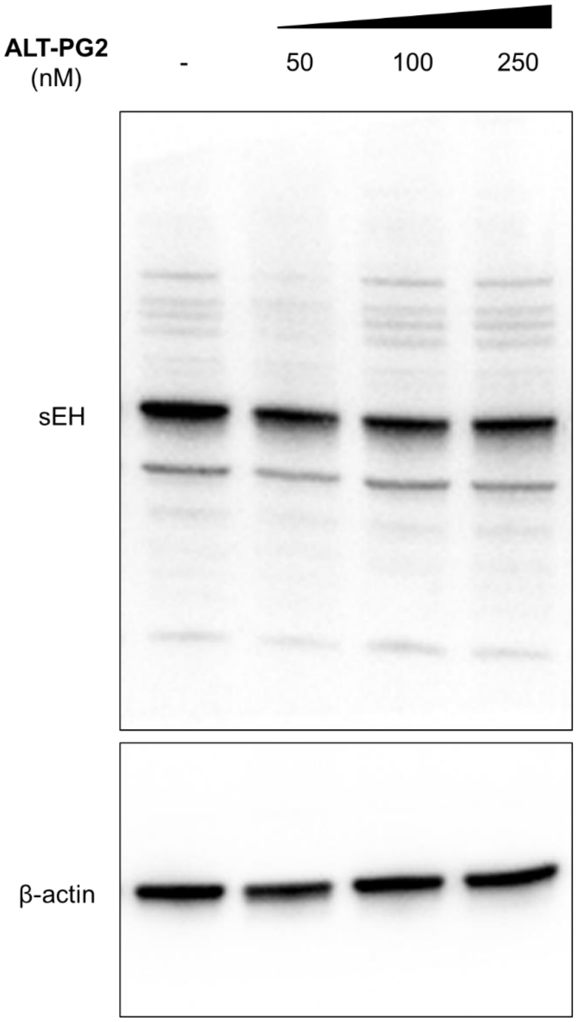

Figure S5

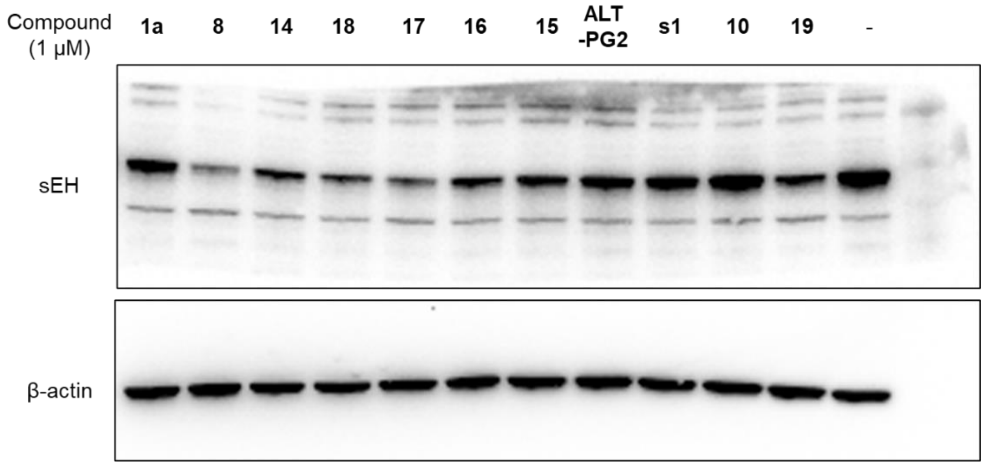


Figure S5A

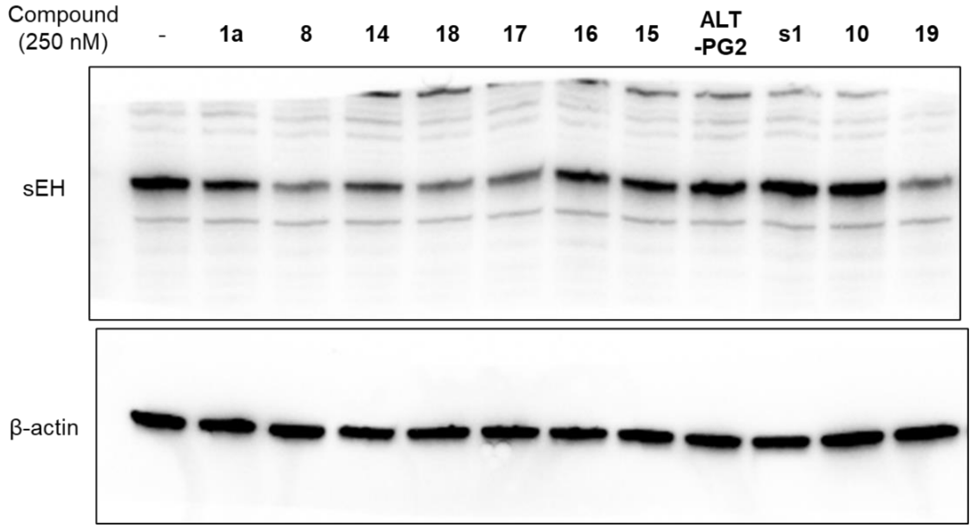


Figure S5B

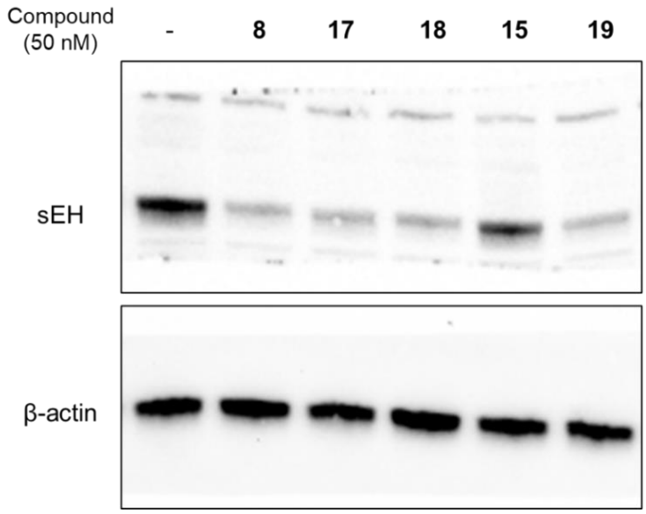


Figure S5C

**Figure S6**

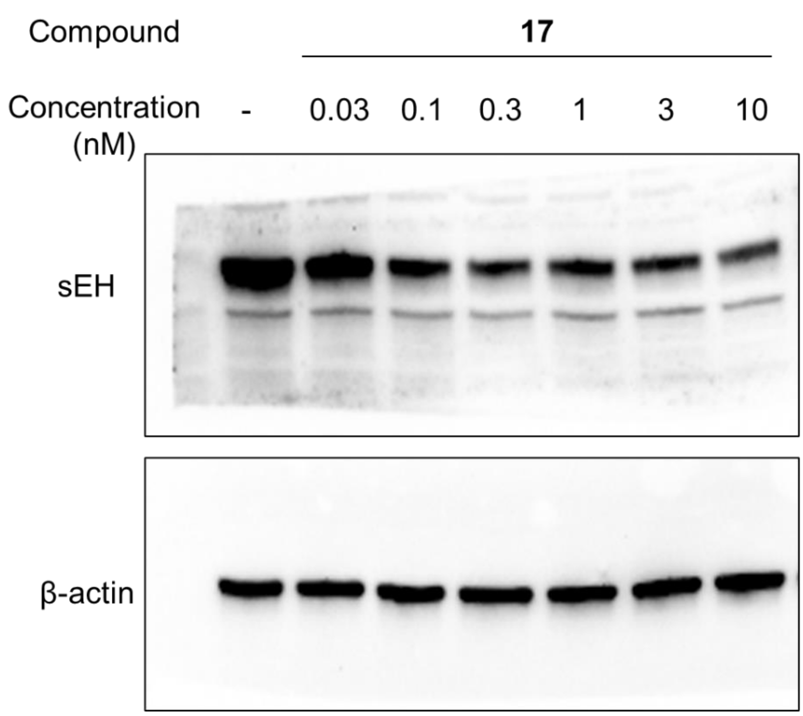


**Figure S6A**

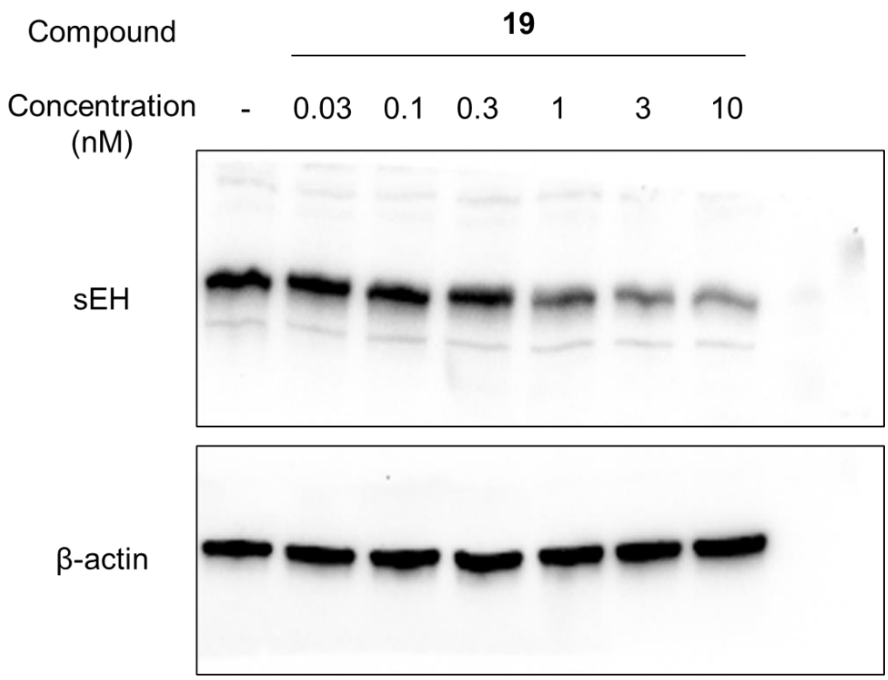


**Figure S6B**

**Figure S7**

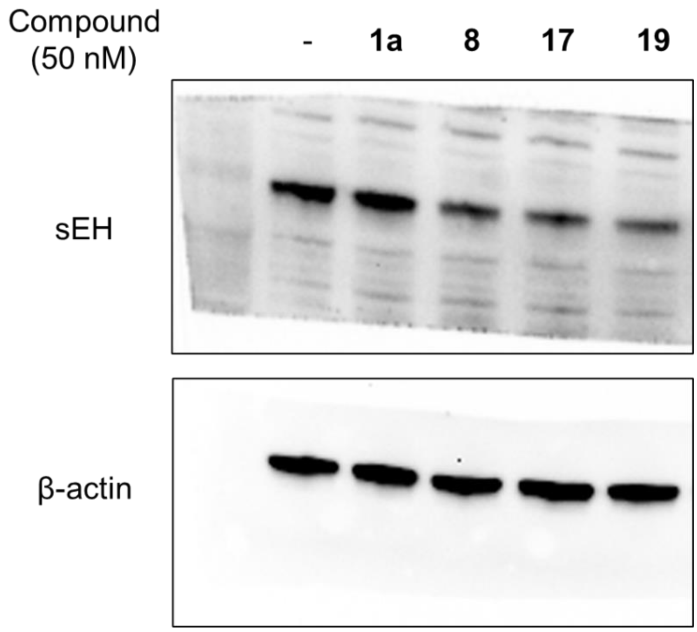

**Figure S8**

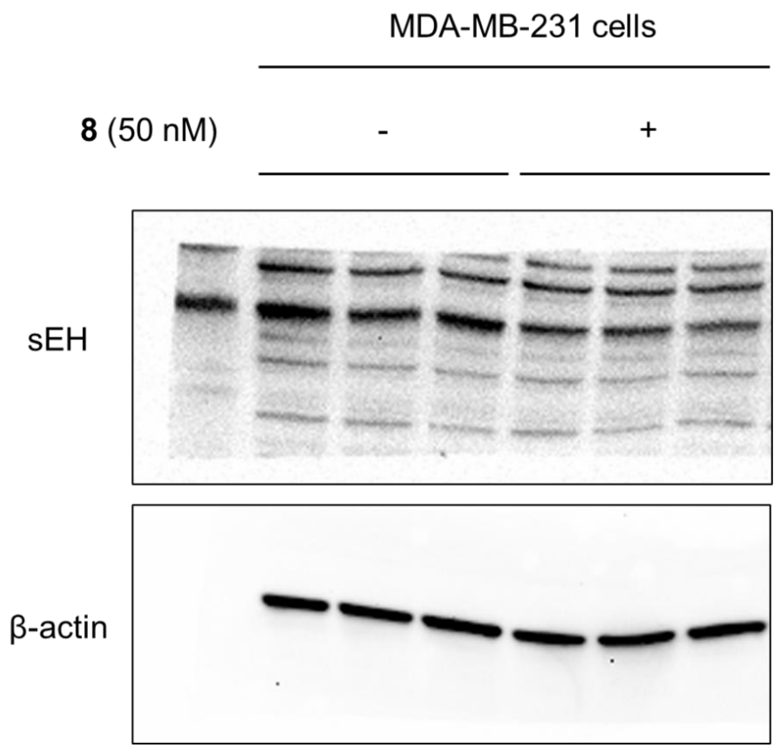


**Figure S8A**

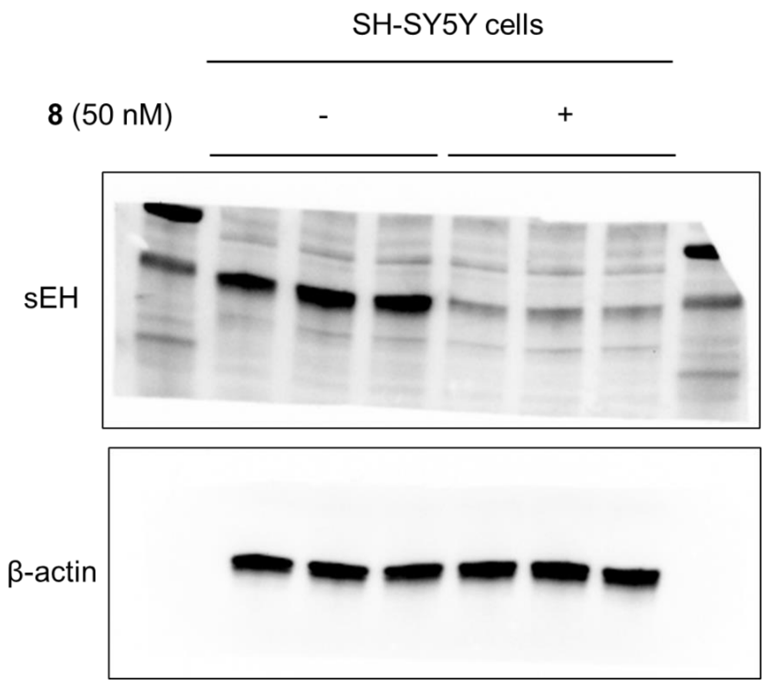


**Figure S8B**
